# Mechanistic and structural insights into a divergent PLP-dependent L-enduracididine cyclase from a toxic cyanobacterium

**DOI:** 10.1101/2023.03.21.533663

**Authors:** Jennifer L. Cordoza, Percival Yang-Ting Chen, Linnea R. Blaustein, Stella T. Lima, Marli F. Fiore, Jonathan R. Chekan, Bradley S. Moore, Shaun M. K. McKinnie

## Abstract

Cyclic arginine noncanonical amino acids (ncAAs) are found in several actinobacterial peptide natural products with therapeutically useful antibacterial properties. The preparation of ncAAs like enduracididine and capreomycidine currently takes multiple biosynthetic or chemosynthetic steps, thus limiting the commercial availability and applicability of these cyclic guanidine-containing amino acids. We recently discovered and characterized the biosynthetic pathway of guanitoxin, a potent freshwater cyanobacterial neurotoxin, that contains an arginine-derived cyclic guanidine phosphate within its highly polar structure. The ncAA L-enduracididine is an early intermediate in guanitoxin biosynthesis and is produced by GntC, a unique pyridoxal-5’-phosphate (PLP)-dependent enzyme. GntC catalyzes a cyclodehydration from a stereoselectively γ-hydroxylated L-arginine precursor via a reaction that functionally and mechanistically diverges from previously established actinobacterial cyclic arginine ncAA pathways. Herein, we interrogate L-enduracididine biosynthesis from the cyanobacterium *Sphaerospermopsis torques-reginae* ITEP-024 using spectroscopic, stable isotope labeling techniques, and X-ray crystal structure-guided site-directed mutagenesis. GntC initially facilitates the reversible deprotonations of the α- and β-positions of its substrate prior to catalyzing an irreversible diastereoselective dehydration and subsequent intramolecular cyclization. The comparison of *holo-* and substrate bound GntC structures and activity assays on sitespecific mutants further identified amino acid residues that contribute to the overall catalytic mechanism. These interdisciplinary efforts at structurally and functionally characterizing GntC enables an improved understanding of how Nature divergently produces cyclic arginine ncAAs and generates additional tools for their biocatalytic production and downstream biological applications.

## INTRODUCTION

Cyclic arginine noncanonical amino acids (ncAAs) are relatively rare in Nature but are highly represented in bioactive natural products.^1,2^ Traditionally isolated via bioactivity-guided fractionation efforts, antimicrobial non-ribosomal peptides like teixobactin,^3^ enduracidin,^4^ mannopeptimycin,^5^ and viomycin^6^ possess these cyclic ncAAs within their chemical structures (Figure 1A). Two isomeric forms currently exist wherein the guanidine moiety is cyclized at either the γ- or β-position of the arginine backbone, creating either the 5-membered enduracididine or 6-membered capreomycidine scaffolds, respectively. Once constructed, enduracididine and capreomycidine ncAAs can be directly incorporated into nonribosomal peptide synthetase (NRPS) products via specific adenylation domain activation,^7^ or further transformed by oxidative enzymes^8,9^ into other cyclic guanidine-containing natural products. Recent advances have shown that capreomycidines can be divergently biosynthesized on NRPSs via dehydrogenation and thioester-mediated Michael addition reactions in the faulknamycin and muraymycin biosynthetic pathways.^10,11^

**Figure 1:**
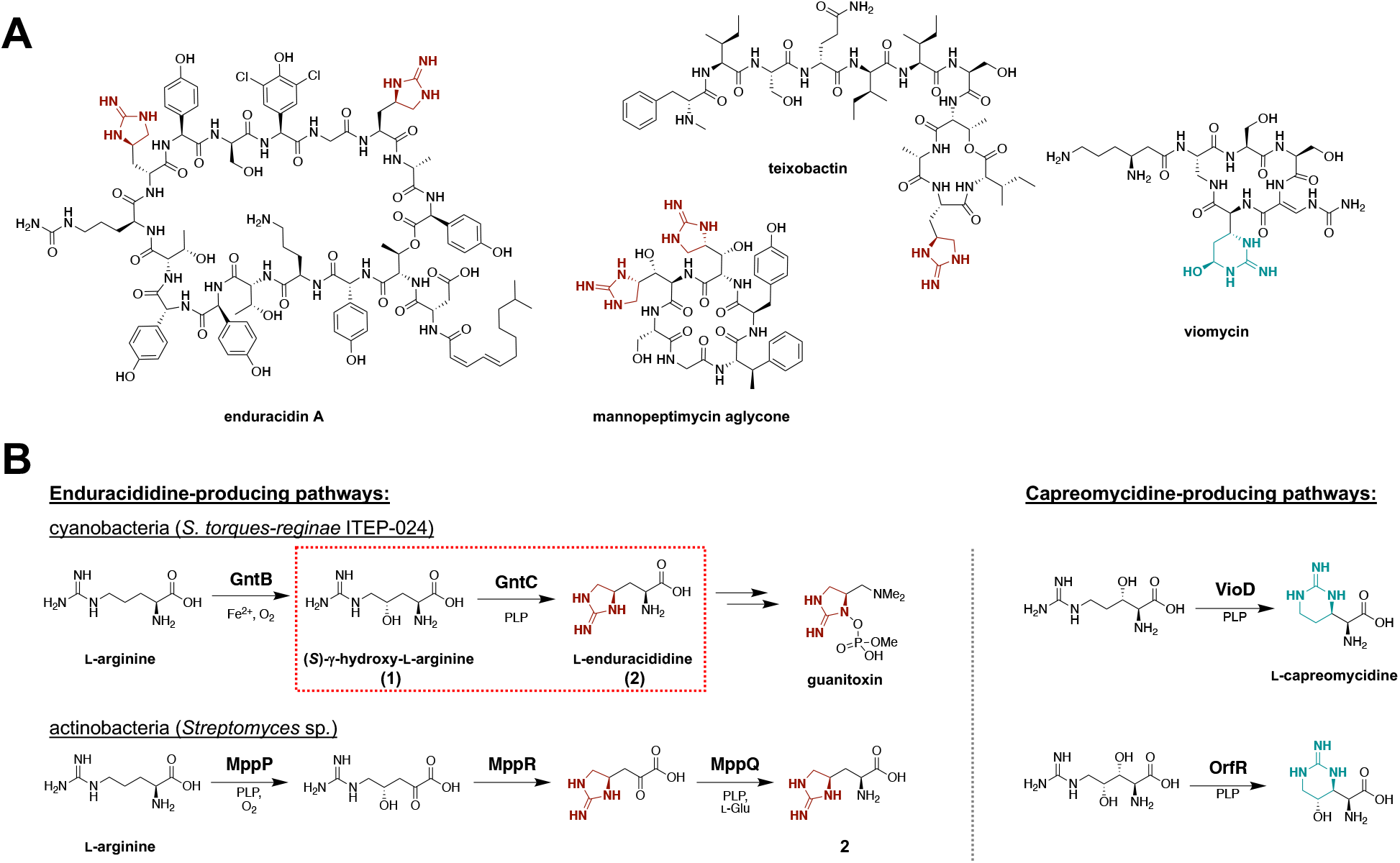
(A) Bioactive antimicrobial nonribosomal peptides containing enduracididine (red) or capreomycidine (blue) cyclic arginine noncanonical amino acids (ncAAs). (B) Characterized biosynthetic strategies towards the construction of cyclic arginine enduracididine (left) and capreomycidine (right) ncAAs. Guanitoxin biosynthesis in cyanobacterium *S. torques-reginae* ITEP-024 proceeds through cyclic arginine intermediate L-enduracididine (**2**) via the cyclodehydration activity of PLP-dependent enzyme GntC. Pyridoxal-5’-phosphate (PLP) enzymology involved in cyclic arginine biosynthesis from actinobacterial mannopeptimycin (MppP, MppQ), viomycin (VioD), and streptilodine (OrfR) pathways.

Currently, cyclic arginine ncAAs have limited commercial availability due to their multiple synthetic steps and heavy reliance on protecting groups.^2^ This has hampered an understanding of their role in establishing the bioactivities of their cognate NRPS products. Advances in small molecule biocatalysis, particularly in ncAA biosynthesis, have created ample opportunities to repurpose these enzymes for their efficient and scalable construction.^12,13^ Moreover, identifying novel methodologies for the biosynthetic construction of cyclic arginine ncAAs would serve as valuable genome mining hooks to discover new natural products with these moieties.

Recently, we discovered the biosynthetic pathway of guanitoxin, a potent freshwater neurotoxin from the cyanobacterium *Sphaerospermopsis torques-reginae* ITEP-024 which possesses a cyclic guanidine moiety in its highly polar structure (Figure 1B).^14^ Previously named anatoxin-a(s),^15^ guanitoxin exerts its neurotoxicity through the irreversible inhibition of acetylcholinesterase within the peripheral nervous system.^16^ If consumed via contaminated water, guanitoxin can lead to an overexcitation of cholinergic neurons and downstream muscle spasms, respiratory arrest, and potentially death. Within the nine-enzyme biosynthetic pathway that transforms L-arginine into the mature neurotoxin, the pyridoxal-5’-phosphate (PLP)-dependent enzyme GntC constructs the key cyclic guanidine moiety upon which the anticholinesterase *O*-methylphosphate pharmacophore is assembled. Specifically, GntC performs a cyclodehydration of substrate (*S*)-γ-hydroxy-L-arginine (**1**) to L-enduracididine (**2**), cleanly inverting the stereochemistry at the γ-position. This reaction is highly diastereoselective under *in vitro* enzyme assay conditions following 1-fluoro-2,4-dinitrophenyl-5-L-alanine amide (l-FDAA, Marfey’s reagent) derivatization and ultra-performance liquid chromatography mass spectrometry (UPLC-MS) analysis. In our initial characterization of GntC, we failed to observe any enzymatic production of *allo-L-en*duracididine diastereomer (**2’**), nor did we observe any conversion of substrate epimer (*Ä*)-γ-hydroxy-L-arginine (**1’**) within the limits of detection.^14^ While the GntC reaction occurs relatively early in the guanitoxin biogenesis, its validation was substantial for establishing the biosynthetic pathway. Both **1** and **2** were previously isolated from guanitoxin-producing *Anabaena flos-aquae* cyanobacteria, and the incorporation of backbone-deuterated **1** into the mature toxin was supported via *in vivo* feeding experiments.^17,18^ Although these stable isotope experiments directly implicated **1** in guanitoxin biogenesis, an intriguing reintroduction of hydrogen atoms at the former β-position implied that additional deprotonation/reprotonation chemistry was involved in toxin production (Figure S1).^18^ Following full pathway characterization,^14^ one of these β-position reprotonations could be justified using the retro-aldol (GntG) and transamination (GntE) reactions identified in this study. However, we propose that a mechanistic interrogation into GntC-mediated cyclization of **1** could help further rationalize this *in vivo* result.

PLP is a widely utilized cofactor that facilitates a diverse range of enzyme-catalyzed chemistry, including transaminations, decarboxylations, and aldol reactions.^19^ PLP-dependent enzymes frequently perform stereo- and regioselective reactions under ambient conditions to produce various molecules, including synthetically challenging ncAAs.^20^ The biosynthesis of **2** has previously been characterized in the actinobacterial mannopeptimycin pathway and employs two PLP-dependent enzymes (MppP, MppQ) and an acetoacetate decarboxylase-like superfamily homolog MppR to analogously generate this cyclic ncAA (Figure 1B).^21–24^ PLP-dependent oxidase MppP converts L-arginine to a γ-hydroxylated a-keto acid intermediate,^22,23^ upon which MppR uses an active site lysine to facilitate a stereoselective 5-membered ring cyclization while retaining the α-keto acid.^21^ A final PLP-dependent transamination by MppQ completes the biosynthesis of **2**.^24^ In contrast, **2** production in cyanobacterium *S. torques-reginae* ITEP-024 employs transmembrane enzyme GntB to generate linear **1**, which is directly cyclized by PLP-dependent GntC. In addition to the functional differences between the two biosynthetic routes to **2**, GntC has low sequence homology to any of the mannopeptimycin enzymes (30%, 34%, 5% sequence similarities to MppP, MppQ, MppR respectively), suggesting a divergent mechanistic and biochemical strategy towards constructing this ncAA.

At a functional level, GntC represents a unique fusion of characterized PLP-dependent enzymes from other biosynthetic pathways that transform amino acid-derived substrates. VioD and OrfR, from the viomycin^25,26^ and streptilodine^9^ pathways respectively, perform an intramolecular cyclization of the guanidine sidechain at the β-position of a hydroxylated arginine-precursor, generating the six-membered ring within the capreomycidine ncAA (Figure 1B). Intermolecular nucleophilic addition at the γ-position of a vinyl glycine ketimine intermediate is precedented in PLP-dependent enzymes like cystathionine-γ-synthase (CGS) from methionine primary metabolism.^27^ This ketimine intermediate has also been observed in specialized metabolite biosynthetic pathways, including: recently characterized fungal enzymes CndF,^28^ Fub7,^29^ and AnkD,^30^ actinobacterial Mur24 involved in antibiotic nucleoside muraymycin biogenesis,^31^ and indirectly through hydration in canavanine catabolism in select Gram-negative proteobacteria.^32^ Although it remains to be biochemically characterized, LolC from the loline biosynthetic pathway is believed to proceed through an analogous ketimine intermediate with an L-proline nucleophile.

Overall, due to its unique sequence homology, divergent chemical reaction, and important role in guanitoxin biosynthesis, we sought to determine the mechanism of GntC-mediated cyclization, rationalize its strict diastereoselectivity and understand structural features responsible for its catalytic function. Through this investigation into how Nature convergently biosynthesizes **2** in phylogenetically distinct microbes, we can better understand the role of this ncAA in divergent natural products and accelerate the biocatalytic application of GntC towards the construction of complex cyclic ncAAs.

## RESULTS AND DISCUSSION

### GntC PLP dependency

To fuel our investigation, substrate (**1**, **1’**) and product (**2**, **2’**) diastereomers were synthesized following previously established synthetic protocols.^14,33^ *S. torquesreginae* ITEP-024 GntC was heterologously expressed as an *E. coli* codon-optimized N-terminal His_6_-tagged construct and purified to near homogeneity using Ni-NTA affinity chromatography.^14^ *In vitro* GntC assays were set up in potassium phosphate (KPi) buffer, amine-containing components were derivatized with l-FDAA, and reaction mixtures were analyzed by UPLC-MS as previously reported.^14^ Because GntC co-purifies with its requisite PLP cofactor, we had yet to firmly confirm its dependency in **2** production. Treatment of purified GntC with PLP inhibitor hydroxylamine followed by buffer exchange abolished any catalytic activity with **1** in the *apo-GntC* form (Figure S2). Re-introduction of two molar equivalents of PLP to the apoenzyme restored **2** production, validating the cofactor dependence of the intramolecular cyclase GntC.

### GntC *in vitro* assay optimization

We initially investigated a range of pHs in KPi buffer to identify ideal conditions for **2** production. Under a 20:1 substrate:enzyme ratio, relative product formation was compared to an l-FDAA-derivatized glycine standard to correct for any derivatization discrepancies, and we determined that GntC behaved optimally at pH 8.0 (Figure S3). Previous reports identified that divalent metal cations can aid PLP-dependent enzyme catalysis through the promotion of electron displacement,^28^ and we wanted to determine if a similar observation occurred with GntC. Counterintuitively, **2** production was not enhanced in the presence of any 1 mM divalent metal salts; instead, incubation with chelator EDTA showed comparable activity to wild type GntC. Most other ionic additives (Mg^2+^, Ca^2+^, Mn^2+^, Fe^2+^, Zn^2+^, Co^2+^) showed similar or reduced *in vitro* **2** formation, with the addition of Cu^2+^ completely abolishing production (Figure S4). We finally investigated the impact of the His6 tag on *in vitro* **2** production, given the significance of the N-terminus on homodimerization, shaping the active site pocket, and establishing catalysis in other PLP-dependent enzymes.^34,35^ We recloned GntC as a C-terminal His6 construct and found a slight reduction in catalytic activity compared to the original N-terminal His6 construct (Figure S5). In spite of additional optimization efforts, *in vitro* GntC assays were done under identical reaction conditions to those previously described.^14^

### GntC spectroscopy

Spectroscopic analyses have been performed on various PLP-dependent enzymes and their conjugated cofactors to identify diagnostic quinonoid intermediates in rate limiting steps of the catalytic cycle,^36–38^ including methionine biosynthetic enzyme CGS^36^ and threonine aldolase enzymes.^39^ Following the incubation of GntC and substrate **1,** we did not observe any extended quinonoid intermediates. Instead, we identified a time-dependent accumulation of an intermediate with an absorbance at 325 nm at the expense of the resting PLP cofactor state at 420 nm (Figures 2, S6). The 325 nm chromophore represents the ketimine intermediate^40,41^ and is mechanistically formed from protonation of the quinonoid at the 4’ position. This is generally observed in canonical primary metabolic PLP-dependent aminotransferases and methionine biosynthetic enzyme CGS. Intriguingly, when substrate diastereomer **1’** was mixed with GntC under identical reaction conditions, a very modest increase of this 325 nm intermediate was observed with minimal perturbations to the PLP system itself, even after prolonged incubation (Figure S6). These data imply that an extended conjugated state is not involved in the rate-limiting step of the GntC mechanism, and that the substrate hydroxyl group stereochemistry plays an important role in not only product output, but spectroscopic features of catalysis.

**Figure 2:**
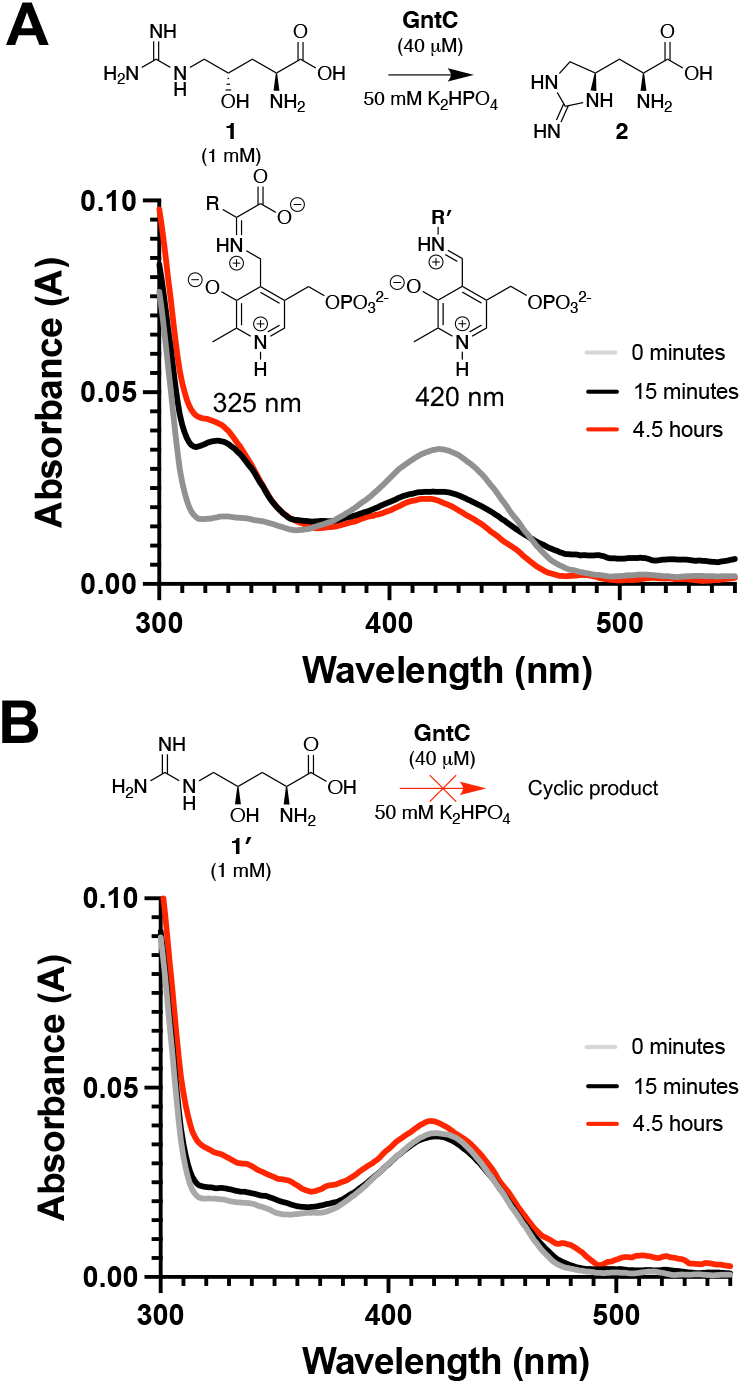
*In vitro* GntC assays with (A) native substrate **1** and (B) diastereomer **1’** show divergent UV-visible absorption spectra at 325 nm and 420 nm, corresponding to the ketimine and aldimine intermediates respectively. R’ represents either a conserved lysine residue (internal aldimine) or an amine substrate (external aldimine).

### Mechanistic studies

To assess the mechanism of enduracididine synthesis by GntC, we initially investigated whether any deprotonation/reprotonation or oxidation events occur at the γ-position of substrate **1**. We modified our established synthetic scheme to selectively reduce the *in situ* ketone intermediate with sodium borodeuteride and install one deuterium atom at the γ-position (Figure 3A).^14^ The deuterated (*S*)-alcohol was resolved from its diastereomer via flash chromatography, and subsequent nitro reduction, guanylation, and deprotection reactions generated the desired γ-deuterated substrate (**1-***d*_z_). *In vitro* incubation of ***1-d_γ_*** with GntC followed by l-FDAA derivatization and UPLC-MS analysis showed that the resultant cyclic product (**2-***d*_z_) was one additional Da heavier than derivatized **2** (Figure 3B) and mirrored the isotopic distribution of synthetic substrate ***1**-d_γ_*. The retention of the γ deuterium atom suggests that no deprotonations occur at this position during the GntC mechanism.

**Figure 3:**
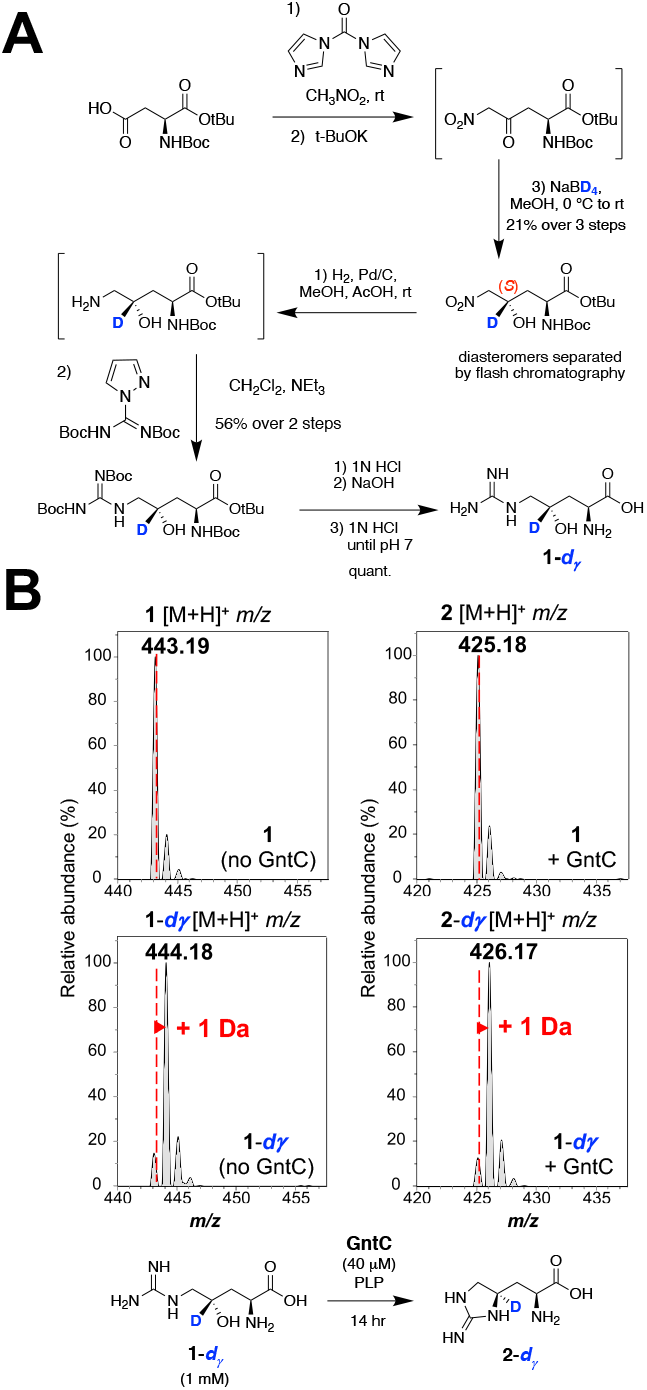
Selective deuteration reveals that GntC does not deprotonate the g-position of substrate **1**. (A) Synthesis of selectively g-deuterated substrate ***1**-d_γ_*. (B) UPLC-MS spectra for l-FDAA derivatized substrates (**1** and **1**-*d_γ_*; left) and products (**2** and **2**-*d_γ_*; right) in the absence and presence of *in vitro* GntC. The increase of +1 Da in product **2-***d_γ_* supports that the g-deuterium atom is retained throughout the GntC mechanism.

To complement the **1**-*d_γ_* assay, we next aimed to identify which substrate hydrogen atoms were reversibly solvent exchanged during the GntC mechanism via buffered deuterium oxide (D_2_O) assay conditions and mass spectrometry analyses. After exchanging GntC into deuterated KPi buffer following previous literature examples,^42,43^ we employed *in vitro* enzymatic assays under 100% D_2_O conditions. Following the assay, l-FDAA derivatization and UPLC-MS analyses were conducted in regular protic solvents to ensure that only deuterium atoms incorporated to the carbon backbone were retained. Moreover, assays were compared to controls lacking GntC to correct for non-enzymatic deuterium incorporation. Analyses of *in vitro* D_2_O reactions identified an increase of 3 Da in product **2** (***2-**d_3_*) in the presence of GntC (Figure 4A), implying that up to 3 deprotonation/reprotonation events occur during the catalytic mechanism. A similar 3 Da increase was observed in substrate **1** (Figure 4A) and inactive diastereomer **1’** (Figure 4B) in the presence of deuterated GntC. This was intriguing as **1’** failed to generate any observable products by UPLC-MS following incubation with GntC.^14^ In combination with the **1-***d_g_* assay results, these data suggested that D-incorporations occurred at the a and β carbons within the GntC active site.

**Figure 4:**
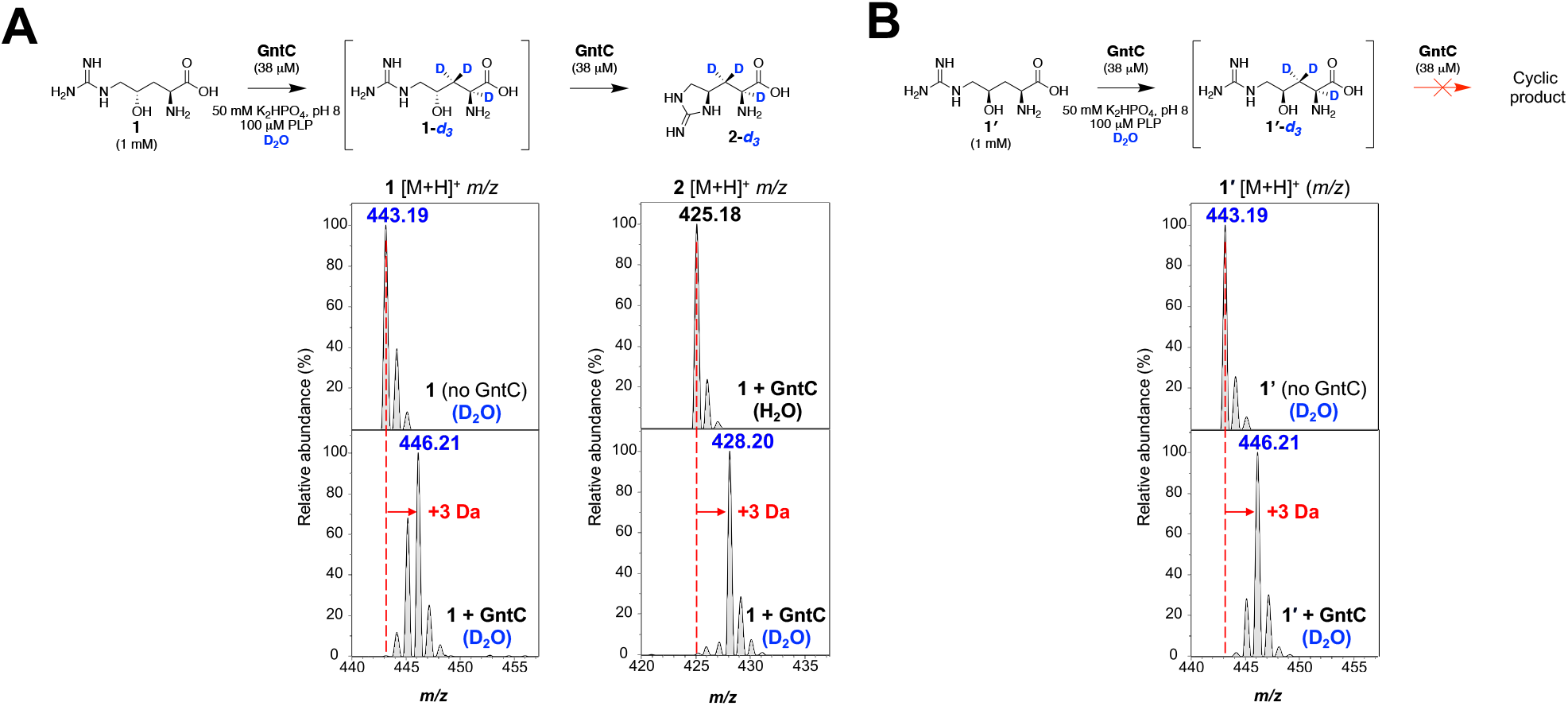
UPLC-MS analyses of l-FDAA derivatized *in vitro* GntC D_2_O assays identify the incorporation of up to three deuterium atoms in g-hydroxy-L-arginine substrates. (A) An increase of +3 Da is observed in linear substrate **1** and cyclic product **2** (**1**-*d_3_* and ***2**-d_3_* respectively) in the presence of D_2_O and GntC. (B) Substrate diastereomer ***1’*** also shows an increase of +3 Da (**1’**-*dj*) in the presence of D_2_O buffered GntC despite any cyclic product formation.

To orthogonally corroborate these results, we employed ^1^H NMR to conclusively identify which hydrogen atoms were solvent exchanged through the course of the GntC reaction. We initially established our assay in protic solvent conditions under a diluted 100:1 substrate: enzyme ratio within a 0.5 mm NMR tube. Following a 14-hour reaction endpoint, we identified the diagnostic a-hydrogen (δ 3.69) and β-hydrogen (δ 2.09) signals that mirrored the synthetic **2** standard (Figure S7). After exchanging GntC into KPi buffered D_2_O and employing analogous *in vitro* conditions, we tracked reaction progression and D-incorporation over time via ^1^H NMR. Under D_2_O conditions, the a-hydrogen of **1** (δ 3.92) was lost after 30 minutes, while the β-hydrogen signals (δ 1.94) began to change in multiplicity over the course of the reaction, finally diminishing after 24 hours (Figure 5). There was an aberrant decrease in one of the multiplexed diastereotopic β-hydrogens, implying a facial preference for this reversible D-incorporation. Consistent with the loss of **1** hydrogen signals, characteristic **2** signals at δ 4.24, 3.85, and 3.38 appeared after the 14-hour mark in the ^1^H-NMR time course, with the notable omission of α- and β-hydrogen atoms. Cumulatively, these results corroborate the 3 Da increase from UPLC-MS analyses under D_2_O conditions and support that deuterium atoms were present at the α- and β-hydrogens within **2**. A similar trend was observed when interrogating substrate diastereomer **1 ‘**, in which α- and β-hydrogen signals were lost in the presence of D_2_O-exchanged GntC, however with the notable absence of any dehydrated or cyclic products (Figure S8). This provides further insight into the GntC mechanism by suggesting that α- and β-deprotonations occur prior to the dehydration and cyclization steps necessary for **2** formation. Moreover, this implies that the stereospecific recognition of the γ-hydroxy group by GntC enables the elimination to proceed.

**Figure 5:**
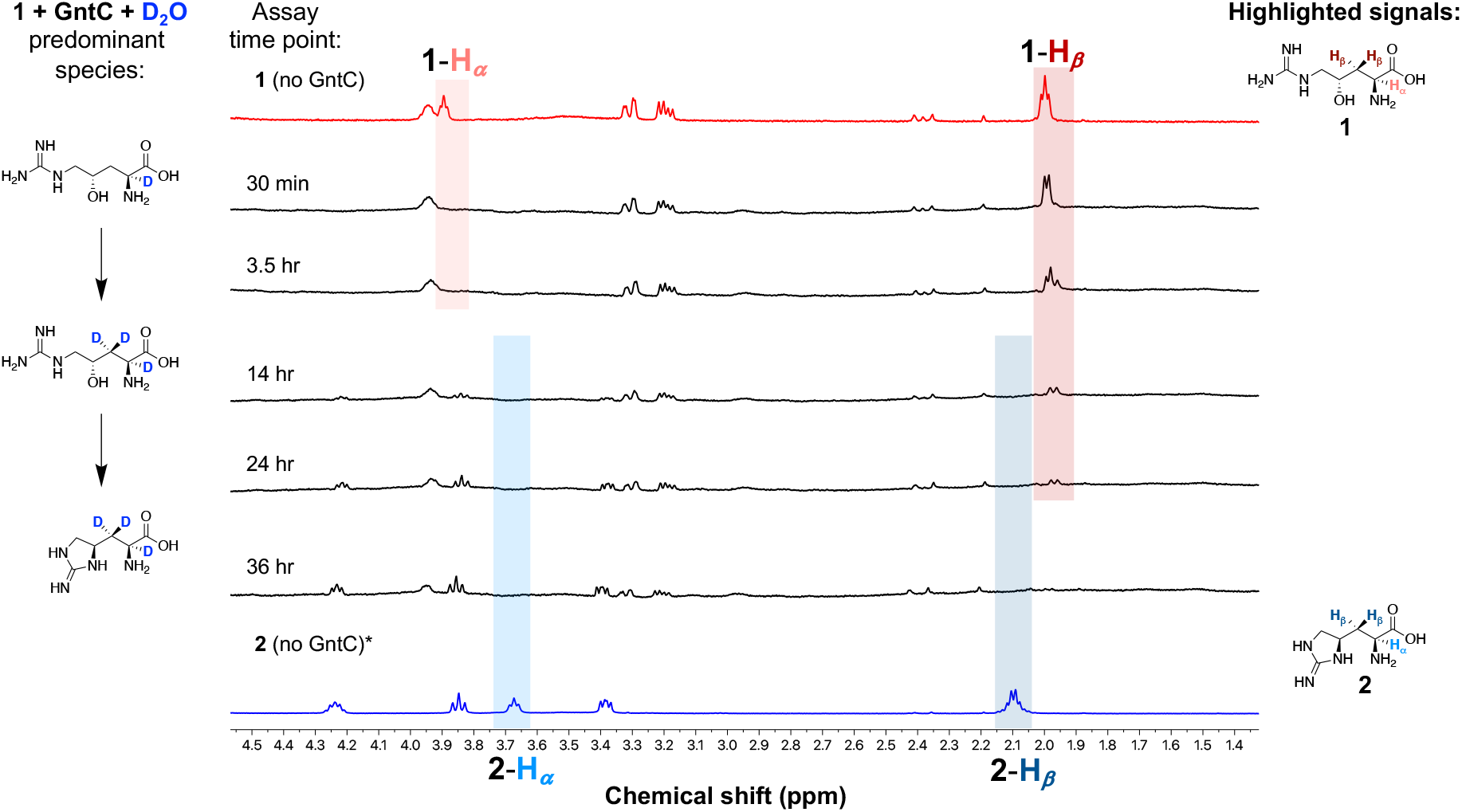
^1^H NMR time course experiments under D_2_O conditions identify that both the α- and β-hydrogen atoms of substrate **1** and product **2** are solvent-exchanged during *in vitro* GntC assays. The predominant chemical species are depicted to the left of GntC reaction time points and compared to **1** (top, red trace) and **2** (bottom, blue trace) standards.

Many PLP transaminases and aldolases exhibit a level of reversibility which can regenerate the starting materials depending on the equilibrium or metabolic flux of the system. To assess the overall reversibility of the GntC reaction, we incubated product **2** with GntC in buffered D_2_O conditions and analyzed the reaction mixture by UPLC-MS after 14 h. While we did observe an increase in 3 Da in product **2** (Figure S9), we did not identify any substrate **1** with any degree of D incorporation nor any other L-arginine-derived products (Figure S10). To further assess the reversibility of the dehydration step, we incubated GntC with substrate **1** under KPi buffered H_2_^18^O. Following a 14-hour assay, UPLC-MS analyses of l-FDAA derivatized **1** showed no difference in the isotopic distribution in the presence or absence of GntC, implying that heavy water is not being incorporated (Figure S11). This reaction deviates from that of the streptolidine PLP-dependent OrfR capreomycidine cyclase, in which H_2_^18^O was reversibly incorporated into its β-hydroxylated arginine substrate.^9^ These cumulative results show that the GntC-catalyzed cyclodehydration of substrate **1** is unidirectional and that the dehydration is an irreversible mechanistic step under the physiological conditions tested.

In summary, our heavy isotope labeling experiments indicate that up to three deprotonations occur at the α- and β-positions of substrate **1**, these occur prior to the elimination of the γ-hydroxyl group and cyclization, and the dehydration and overall GntC reaction is not reversible under *in vitro* conditions. Moreover, these results highlight a divergence of this cyanobacterial PLP-dependent arginine cyclase enzyme in contrast to its actinobacterial orthologs. Given the D_2_O results of the substrate diastereomer **1’**, we hypothesized the enzyme active site was responsible for the diastereoselectivity of GntC and next pursued enzyme structural studies to further understand the GntC mechanism and the biochemical source of this selectivity.

### GntC sequence homology to PLP-dependent cyclases

After gaining mechanistic insights into GntC catalytic activity, we compared its amino acid sequence with validated PLP-dependent arginine cyclases to bioinformatically determine any conserved residues that may contribute to the overall mechanism. In addition to the active site lysine (K219) that binds the PLP cofactor, sequence alignments of GntC to PLP-dependent capreomycidine cyclases VioD^25,26^ and OrfR^9^ revealed a conserved glutamate residue (E9) and two consecutive serine residues (S25, S26) towards the N-terminus of all three enzymes (Figure S12). OrfR and VioD shared a slightly higher percent sequence similarity to GntC (38% and 44%, respectively) than other PLP-dependent enzymes. Outside of PLP-dependent cyclases, we also compared the amino acid sequence of GntC to functionally analogous PLP-dependent enzymes that perform divergent intermolecular γ-substitutions (CndF, LolC, CGS, Figure S13). Additionally, as GntC bioinformatically annotates as a Type I/II PLP-dependent aminotransferase, we independently compared its sequence to *E. coli* AspC (UniProt ID: P00509). The subset of PLP-dependent intermolecular γ-substitution enzymes and aminotransferases displayed comparatively lower (21 – 31%) amino acid sequence similarities. CGS exhibited the highest percent similarity to GntC and shared residues for PLP anchoring such as the conserved lysine (K219) for internal aldimine formation and aspartate (D186) for pyridine ring nitrogen protonation (Figure S13). These observations suggest there are minimal similar residues at a sequence level between GntC and other γ-substitution enzymes that would bioinformatically contribute to product conversion.

### GntC crystal structure and mutagenesis

To gain further insight into the structural features that facilitate catalysis, we obtained crystal structures of GntC with bound PLP (PDB ID: 8FFT). Following heterologous expression and purification, GntC crystals were obtained using a hanging drop method and the structure was solved to a 2.10-Å resolution using molecular replacement with a PLP-dependent aminotransferase from *Streptococcus suis* (PDB ID: 3OP7, 29% sequence identity to GntC) (Figure S14). GntC crystallized as a dimer of homodimers and adopted a largely a-helical structural fold. PLP was bound to K219 in each monomer as an internal aldimine; this was consistent with sequence alignment results that suggested this lysine was responsible for anchoring the cofactor in the active site.^44^ We subsequently set up individual crystallization trials with PLP-bound GntC and each available substrate (**1**, **1’**) and product (**2**, **2’**) diastereomer. Only **1** successfully cocrystallized with GntC (80% occupancy in one of the monomers) and this external aldimine-containing structure (PLP-**1**) was solved to 2.04-Å resolution (PDB ID: 8FFU) (Figure 6A). PLP-bound and PLP-**1** GntC structures were highly similar (0.31 Å root-mean-square deviation (RMSD) for backbone atom overlap) with the major structural difference being an additional N-terminal ordered helix that helps form the active site in the PLP-**1** bound complex (Figure S14). When comparing PLP-bound and PLP-**1** internal and external aldimine substrates, there was a 30° downwards rotation of the pyridine ring, orienting the amino acid substrate towards a new ordered N-terminal helix that was not present in the first structure. Consistent with many type I/II PLP dependent enzymes,^19,44^ the GntC active site is found at the homodimer interface. Although it is predominantly composed of residues from one of the monomers, important contributions from the corresponding homodimer occur on at least one face of the active site pocket.

**Figure 6:**
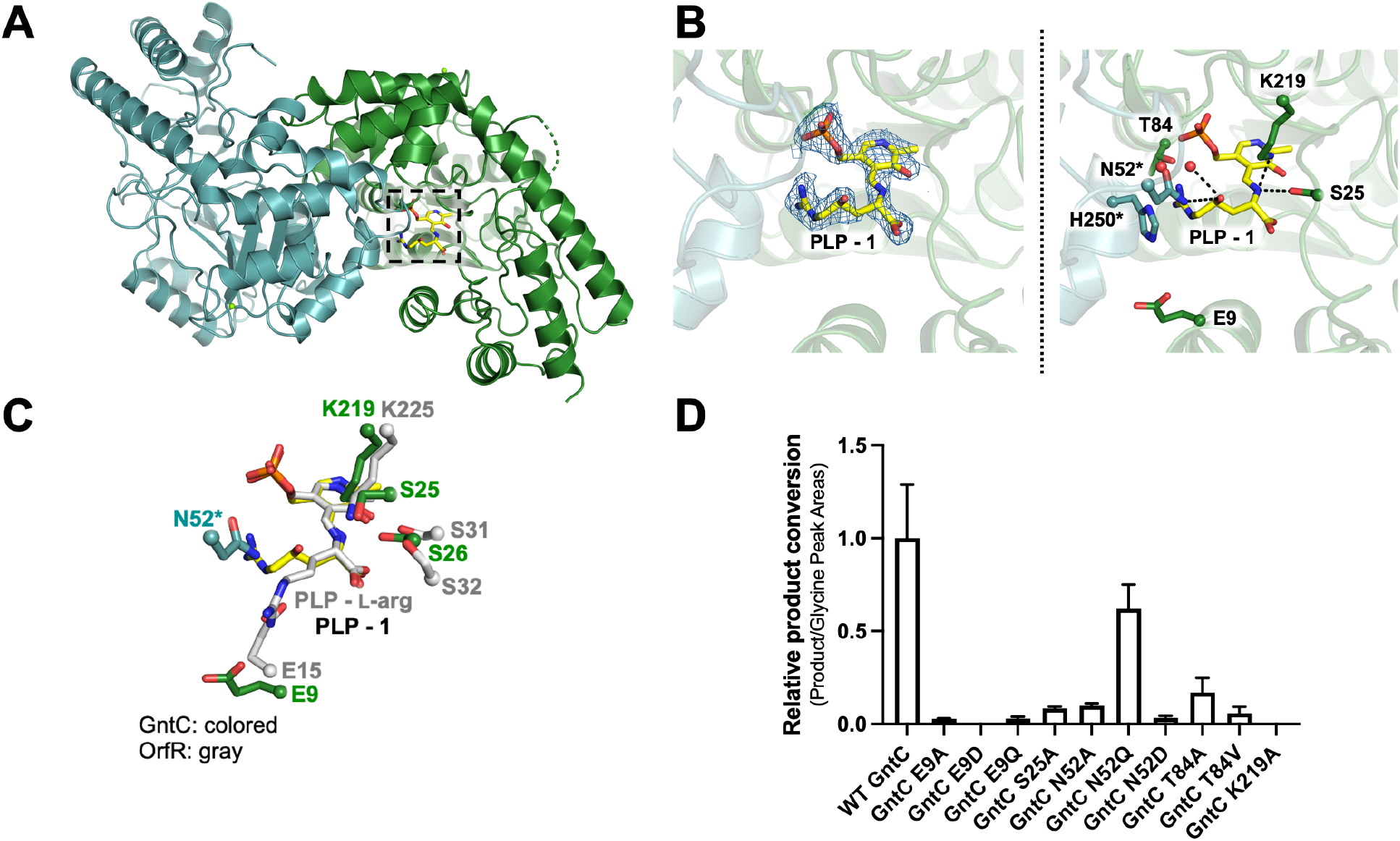
GntC X-ray crystal structure and site-directed mutagenesis identify important amino acid residues for catalysis. (A) GntC crystal structure with bound PLP-**1** substrate (yellow) crystallized as a dimer of homodimers at 2.04 Å resolution (PDB ID: 8FFU, only one homodimer depicted for simplicity). The active site is at the interface of the homodimer between the green and teal monomers, consistent with other PLP-dependent enzyme structures. (B) GntC active site with Fo-Fc electron density map for bound PLP-**1** (blue mesh, σ = 1.0) imposed on the ligand (left) and amino acid residues proposed to participate in substrate recognition and catalysis (right), including E9, S25, T84, K219, (green monomer) and N52*, H250* (teal monomer). (C) Simplified active site comparison of GntC (green/teal) and capreomycidine cyclase OrfR residues (gray, PDB ID: 4M2M) with bound PLP-**1** (yellow) and PLP-L-arg (gray) substrates respectively. (D) *In vitro* enzyme assays for **2** production with soluble GntC mutants in comparison to wild type (WT) GntC.

We next sought to identify GntC active site residues within hydrogen bonding distances of the PLP-**1** adduct that could contribute to catalysis and diastereoselectivity (Figure 6B). Conserved active site K219 lies 2.8 Å from the a-amine forming the external aldimine, T84 is located 2.4 Å away from the PLP phosphate and 3.3 Å away from the guanidinium moiety of **1**, H250 putatively shapes the guanidinium binding pocket, and S25 is within close proximity to the a-carboxylate. Given the mechanistic importance of the γ-hydroxyl group of **1**, we searched for residues of interest near this moiety to help dictate the diastereospecificity of GntC cyclization. The only amino acid side chain in proximity was N52, which was 4.1 Å away and contributed from the other subunit in the dimer (defined as N52* herein). Other residues involved in PLP binding (Figure S14) include D186 (2.8 Å from the pyridoxal ring nitrogen), H188 (2.8 Å from the PLP phosphate group), N158 and Y189 (both within 3 Å of the PLP hydroxyl group), and F108 (π stacking with PLP cofactor). Additionally, R346 was positioned 2.9 Å from the a-carboxylate of substrate PLP-**1**, and presumably contributes to amino acid substrate recognition. GntC requires an active site base to support the α- and β-deprotonation results obtained; based on the PLP-**1** structure, the best candidate is K219.

When the crystal structures of GntC and OrfR (PDB ID: 4M2M) were aligned (1.7 Å RMSD for overlap), S25 and S26, K219, and E9 in the GntC crystal structure overlaid with OrfR residues identified by sequence alignment (Figure 6C). These analogous amino acid residues were previously biochemically validated to have important mechanistic roles in the OrfR catalyzed PLP-dependent formation of γ-hydroxylated capreomycidine ncAA.^9^ Briefly, the two continuous serine residues (OrfR S31 and S32) coordinate with active site lysine (OrfR K225) to facilitate the initial a-deprotonation of dihydroxylated L-arginine via an activated water molecule, and the conserved glutamate residue, (OrfR E15), putatively helped tune the nucleophilicity of the guanidine prior to cyclization. Given the similar functions of the two enzymes, we hypothesized these residues could serve similar purposes in GntC. However, the divergent cyclic ncAA connectivity and heavy isotope incorporation mechanistic differences fail to provide conclusive identification to the source of γ-hydroxy recognition and diastereoselectivity through this OrfR structural comparison.

The most chemically analogous enzymatic reaction to GntC is arguably MppR, which catalyzes the intramolecular stereoselective cyclization en route to **2** biosynthesis.^21^ However, MppR significantly diverges as it does not use a PLP-cofactor and works upon keto-acid containing starting materials and products. Unsurprisingly, very minimal amino acid sequence or structure alignment similarities (PDB ID: 4JME) were observed between these topologically distinct enzymes with different quaternary structures. Negligible active site homology exists between these two enzymes, and additional comparisons were not pursued despite the similar cyclization functions. Although it does not participate in arginine cyclization biochemistry, we performed a structure alignment of GntC to known crystal structure of type I PLP-dependent enzyme aspartate aminotransferase *E. coli* AspC^45^ (PDB ID: 1ARS); this analysis revealed few conserved residues outside of those needed for PLP binding activity and a-carboxylate recognition (Figure S15).

Outside of known protein crystal structures, we also aligned GntC to AlphaFold 2.0 models^46^ of VioD^25,26^ and Fgm3^47^ from the viomycin and fusaoctaxin biosynthetic pathways respectively. VioD catalyzes the PLP-dependent formation of a 6-membered capreomycidine ring analogous to OrfR but with a singly β-hydroxylated L-arginine precursor. Structural alignment with modeled VioD supported the role of previously identified GntC amino acid residues (K219, R346, S25, S26, E9, F108) implicated in PLP-**1** interaction and catalysis (Figure S16). Unique similarities to the GntC and VioD comparative model were R227 and Y189, however both interact with the PLP cofactor instead of the bound substrate. Lastly, Fgm3 is a PLP-dependent enzyme that uses **1** as a substrate to catalyze a retro-aldol reaction.^47^ Aligning modeled Fgm3 and GntC identified the conservation of active site K219 and T84 that coordinates with the guanidine group of **1** (Figure S17). Fgm3 contains a histidine residue instead of F108 in GntC for π stacking above the pyridine ring of PLP. Intriguingly, Fgm3 has a tyrosine residue (Y225) in proximity to the γ-hydroxyl group of PLP-**1** that is absent in GntC. Y225 in Fgm3 putatively could serve as the general base needed for the retro-aldol activity of this enzyme and biochemically rationalize the divergent chemistries these two enzymes exhibit upon the same substrate.

Using these cumulative structural comparisons, we selected six residues of interest hypothesized to contribute to GntC catalysis (E9, S25, N52, T84, K219, and H250). We generated site specific alanine mutants at these six positions and additionally prepared GntC variants with more conservative mutations to address specific features about that amino acid side chain. E9D and E9Q were engineered to investigate the impact of chain length and hydrogen bond donor ability of this previously established catalytic residue in OrfR, T84V and T84S were targeted to assess if pocket shaping or hydrogen bonding are important for guanidinium recognition, and N52D and N52Q mutants were designed to probe the impact of hydrogen bond donation ability and chain length respectively on putative γ-hydroxyl recognition and elimination. GntC mutants were heterologously expressed and purified using previously described procedures except H250A and T84S which were insoluble and not pursued further. Given the proposed influence of homodimerization on shaping the GntC active site and role of N52* in γ-hydroxyl recognition, all purified mutants were assessed via analytical size-exclusion chromatography and displayed comparable levels of dimerization to wild-type GntC (Figure S18).

The five soluble GntC alanine mutants (E9A, S25A, N52A, T84A, K219A) were assayed for **2** production in triplicate and normalized to wild-type GntC using a 500 μM glycine internal standard following l-FDAA derivatization and UPLC-MS analysis (Figure 6D). In all cases, a reduction of **2** formation was observed, indicating the importance of these residues in GntC catalysis. We hypothesized that K219A would disrupt the external aldimine resting state of GntC and remove the covalent tether retaining PLP in the active site. The complete loss of **2** production *in vitro* validated that K219 is an essential residue. Mutating nearby S25 to an alanine reduced **2** production by an order of magnitude, suggesting that organization within this area is beneficial for GntC catalysis. E9 mutation also showed a substantial loss of **2** production regardless of the amino acid residue mutation (E9A, E9D, E9Q). This glutamate residue was identified from the sequence alignment of OrfR and VioD and was hypothesized to help the cyclization mechanism. Within GntC, E9 is 7.7 Å away from the hydroxyl group in the PLP-**1** bound structure, so we propose that it plays a stabilizing role in the second-shell coordination of the substrate or construction of the active site. The T84A and T84V mutants were designed to disrupt any hydrogen bonding interactions that might coordinate the incoming guanidinium nucleophile; decreased **2** production with both mutants provides support for this claim. Finally, the N52 mutant series provided the largest variations in **2** formation. The N52A mutant reduced product by approximately ten-fold, with even a further decrease by the N52D mutation. However, we observed a significant retention of activity with the N52Q mutant, suggesting that the amide side chain of N52 plays an important role in GntC catalysis. We hypothesize the amide hydrogen bonds with the γ-hydroxyl of **1** to make it a better leaving group during the GntC-catalyzed dehydration step of the mechanism either directly or via a hydrogen-bonded network of water. Although the N52 side chain is 4.1 Å away from the γ-hydroxyl within the PLP-**1** bound GntC structure, there could be additional conformational flexibility during catalysis or substrate reorientation following α- and β-deprotonations. The reduction of **2** production during the N52A and N52D mutants further supports this hypothesis (Figure S19). Amide-containing amino acid residues have been found to play important catalytic roles in PLP-dependent enzymes, such as Q52 in dialkylglcyine decarboxylase, which facilitates favorable stereoelectronic interactions during the reaction. ^48^ These data give insight into the inner biochemical workings of GntC catalysis and enable its biocatalytic application towards ncAA syntheses.

### GntC mechanistic proposal

Based on cumulative spectroscopic, UPLC-MS, NMR, and X-ray crystallography-derived site-directed mutagenesis results, we have proposed the following GntC mechanism (Scheme 1). Initially, substrate **1** forms an external aldimine (I) with the PLP cofactor, displacing the active site K219 tether. Subsequent a-hydrogen deprotonation, either by K219 or a coordinated water molecule, generates the canonical quinonoid species (II), which can be reprotonated to rearomatize the PLP cofactor and form ketimine III. Reversible deprotonation of the β-hydrogens facilitates the formation of enamine IV. The side chain amide of N52* next forms a diastereoselective hydrogen bond either directly or through a coordinated water molecule with the γ-hydroxy group to power the irreversible dehydration and subsequent formation of a,β-unsaturated imine intermediate V. Afterwards, the T84-coordinated guanidine moiety performs an irreversible intramolecular Michael addition to form the cyclic ring and resultant enamine (VI). Reprotonation at the β-position regenerates ketimine VII, which can be subsequently deprotonated to quinonoid (VIII) and reprotonated at the a-position to form PLP- **2** (intermediate IX). Final replacement with K219 liberates product **2** and regenerates the internal aldimine to complete the catalytic cycle.

**Scheme 1:**
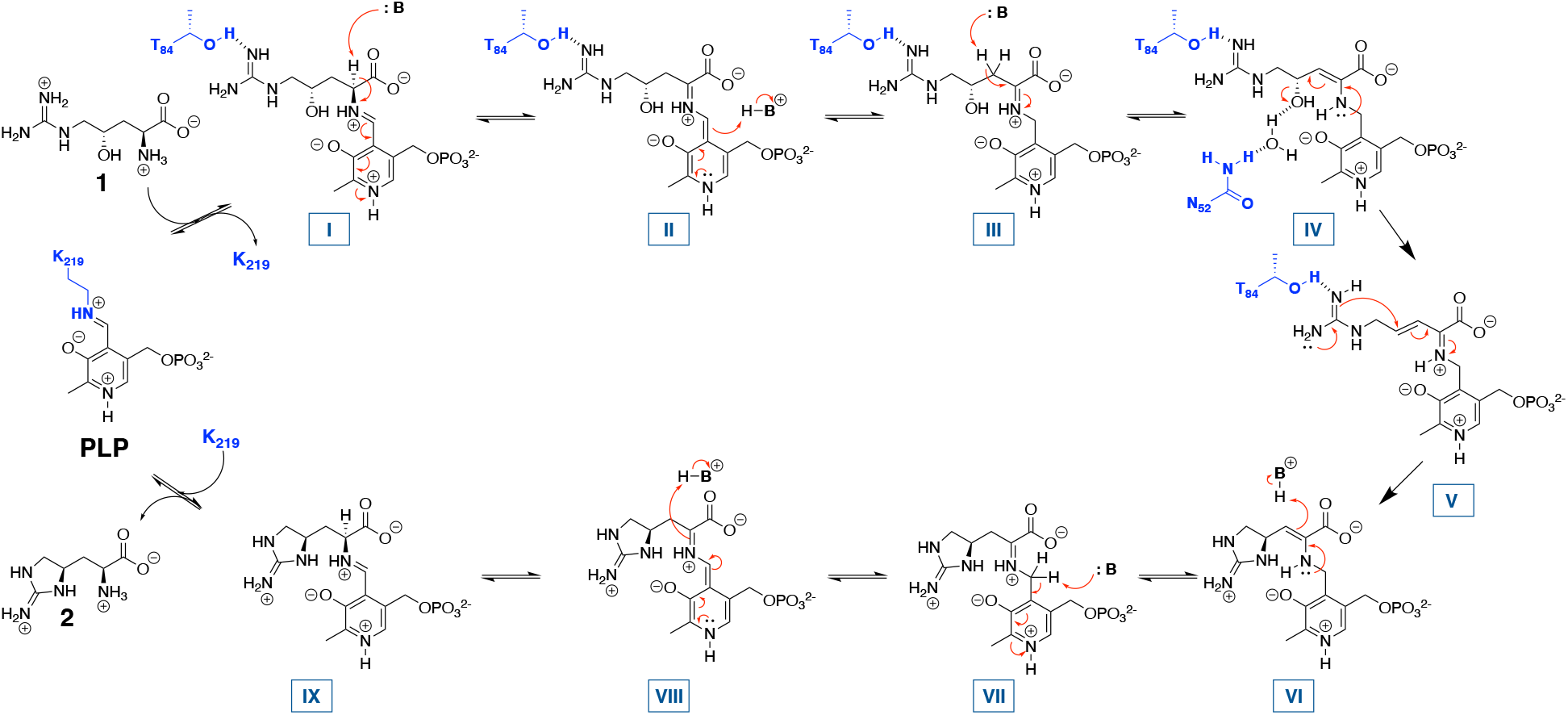
Proposed GntC mechanism based on cumulative spectroscopic, heavy-isotope labeling, NMR, and X-ray crystallography and site-directed mutagenesis results.

While the early and late mechanistic steps are consistent with many canonical PLP-dependent enzymes, the novelty around the hydroxyl group elimination and subsequent intramolecular cyclization are what distinguish this cyanobacterial homolog from its more distant orthologs. Capreomycidine cyclases VioD and OrfR use the nucleophilicity of the quinonoid intermediate to catalyze the β-hydroxyl group elimination, and then adapt the extended conjugation and electrophilicity of the resulting pyridinium cofactor to facilitate the intramolecular guanidine cyclization (Figure S20). The absence of an extended chromophore in GntC spectroscopic analyses immediately distinguished this proposed mechanism from known actinobacterial β-hydroxyarginine cyclases. Intermolecular PLP-dependent γ-substitution enzymes, including CGS, CndF, Fub7, AnkD, Mur24, and putative LolC, follow a Michael-type addition of diverse nucleophiles onto a vinyl-glycine ketimine derived from a γ-functionalized amino acid substrate (Figure S21). In contrast, GntC sets up ketimine intermediate V with only a γ-alcohol as a leaving group and uses an intramolecular guanidine moiety as a nucleophile instead. Through the mechanism, GntC uses a γ-substitution strategy (like CndF and CGS) to perform the intramolecular cyclization in the formation of a 5-membered ring ncAA (like non-PLP-dependent enzyme MppR). This cyanobacterial biosynthetic strategy represents a hybrid of two different approaches to generate cyclic arginine ncAA **2.**

GntC adds to the repertoire of PLP-dependent enzymes that have been recently discovered that catalyze unconventional reactions. Select examples include: BesB from the β-ethynylserine pathway that creates a terminal alkyne;^49^ SbzP which catalyzes a γ-substitution of β-nicotinamide adenine dinucleotide onto *S*-adenosylmethionine in the altemicidin biosynthetic pathway;^50^ Claisen chemistry in saxitoxin^51^ and ketomemicin^52^ biosyntheses; the decarboxylative aldol reaction facilitated by UstD that has seen exceptional biocatalytic application;^53^ and the diverse range of PLP-dependent oxidases that have been uncovered in recent years.^22,54^ Despite the ubiquitous presence of PLP in primary and secondary metabolic enzymes, Nature continues to adapt this cofactor in unique ways to generate diverse and biocatalytically useful chemical transformations.

## CONCLUSION

In conclusion, GntC is the first characterized example of an L-enduracididine cyclase from a cyanobacterium. Through its interdisciplinary interrogation using spectroscopic, stable isotope labeling, NMR, MS, and X-ray crystal structure-function studies, we conclude that GntC follows a distinct mechanism from characterized actinobacterial PLP-dependent cyclases and behaves more like an intermolecular γ-substitution PLP-dependent enzyme. This study seeks to better understand the biosynthesis of cyclic arginine ncAAs from diverse microbial species. Moreover, this improved understanding creates opportunities to apply GntC as a biocatalyst for the scalable production of known and novel ncAAs.

## Supporting information

GntC-SI-PDF

## ASSOCIATED CONTENT

### Supporting Information

This material is available free of charge on the ACS Publications website at http://pubs.acs.org.

General materials and methods, Figures S1-S21, Tables S1-S4, chemical synthesis and characterization, and NMR spectra (PDF)

GntC + PLP PDB ID: 8FFT

GntC + PLP-**1** PDB ID: 8FFU

## AUTHOR INFORMATION

### Present Addresses

*†If an author’s address is different than the one given in the affiliation line, this information may be included here.*

*Present Address for P.Y.T.C.*

Morphic Therapeutics, Waltham, Massachusetts 02541, United States

Present Address for S.T.L.

Department of Chemistry and Biochemistry, University of North Carolina at Greensboro, North Carolina 27402, United States

### Author Contributions

J.L.C., P.Y-.T.C., J.R.C., B.S.M., and S.M.K.M. designed this study. Small molecule organic synthesis, characterization, site-directed mutagenesis, and enzyme assays were conducted by J.L.C. and L.R.B. Protein crystallization was performed by P.Y-.T.C., S.T.L., J.R.C., and B.S.M. GntC genomic information and plasmids were provided by S.T.L., and M.F.F. The manuscript was prepared by J.L.C. and S.M.K.M. with input from all authors, and everyone has given approval to the final version of the manuscript.

## ACKNOWLEDGMENTS

We acknowledge Prof. L. M. Sanchez and H. J. Lusk for assistance with obtaining high-resolution mass spectrometry data, Dr. H.-W. Lee for maintenance of nuclear magnetic resonance spectroscopy facilities, and P. Ngoi for help with analytical size exclusion chromatography (all University of California, Santa Cruz). We also thank Prof. J. P. Noel and Dr. G. Louie (Salk Institute for Biological Studies) for beamtime coordination.

## ABBREVIATIONS

l-FDAA: 1-fluoro-2,4-dinitrophenyl-5-L-alanine amide, Marfey’s reagent
ncAA: noncanonical amino acid
NRPS: non-ribosomal peptide synthetase
PLP: pyridoxal-5’-phosphate
RMSD: rootmean-square deviation.

## For Table of Contents Only

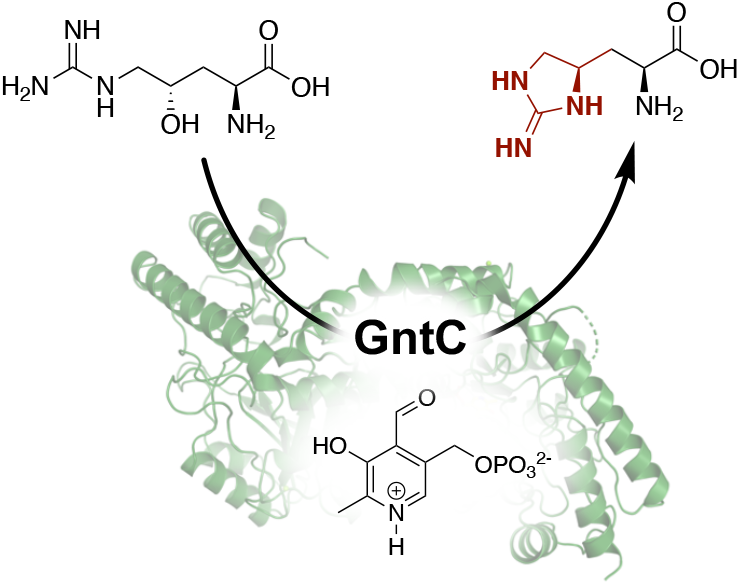

## REFERENCES

(1) Berlinck, R. G. S.; Bertonha, A. F.; Takaki, M.; Rodriguez, J. P. G. The Chemistry and Biology of Guanidine Natural Products. Nat. Prod. Rep. 2017, 34 (11), 1264–1301. https://doi.org/10.1039/C7NP00037E.

(2) Atkinson, D. J.; Naysmith, B. J.; Furkert, D. P.; Brimble, M. A. Enduracididine, a Rare Amino Acid Component of Peptide Antibiotics: Natural Products and Synthesis. Beilstein J. Org. Chem. 2016, 12, 2325–2342. https://doi.org/10.3762/bjoc.12.226.

(3) Ling, L. L.; Schneider, T.; Peoples, A. J.; Spoering, A. L.; Engels, I.; Conlon, B. P.; Mueller, A.; Schäberle, T. F.; Hughes, D. E.; Epstein, S.; Jones, M.; Lazarides, L.; Steadman, V. A.; Cohen, D. R.; Felix, C. R.; Fetterman, K. A.; Millett, W. P.; Nitti, A. G.; Zullo, A. M.; Chen, C.; Lewis, K. A New Antibiotic Kills Pathogens without Detectable Resistance. Nature 2015, 517 (7535), 455–459. https://doi.org/10.1038/nature14098.

(4) Higashide, E.; Hatano, K.; Shibata, M.; Nakazawa, K. Enduracidin, a New Antibiotic. J. Antibiot. (Tokyo) 1968, 21 (2), 126–137. https://doi.org/10.7164/antibiotics.21.126.

(5) Singh, M. P.; Petersen, P. J.; Weiss, W. J.; Janso, J. E.; Luckman, S. W.; Lenoy, E. B.; Bradford, P. A.; Testa, R. T.; Greenstein, M. Mannopeptimycins, New Cyclic Glycopeptide Antibiotics Produced by *Streptomyces hygroscopicus* LL-AC98: Antibacterial and Mechanistic Activities. Antimicrob. Agents Chemother. 2003, 47 (1), 62–69. https://doi.org/10.1128/AAC.47.1.62-69.2003.

(6) Finlay, A. C.; Hobby, G. L.; Hochstein, F.; Lees, T. M.; Lenert, T. F.; Means, J. A.; P’an, S. Y.; Regna, P. P.; Routien, J. B.; Sobin, B. A.; Tate, K. B.; Kane, J. H. Viomycin, a New Antibiotic Active against Mycobacteria. Am. Rev. Tuberc. 1951, 63 (1), 1–3.

(7) Chen, I.; Cheng, T.; Wang, Y.; Huang, S.; Hsiao, Y.; Lai,Y.; Toh, S.; Chu, J.; Rudolf, J. D.; Chang, C. Characterization and Structural Determination of CmnG-A, the Adenylation Domain that Activates the Nonproteinogenic Amino Acid Capreomycidine in Capreomycin Biosynthesis. ChemBioChem 2022, 23 (24). https://doi.org/10.1002/cbic.202200563.

(8) Haltli, B.; Tan, Y.; Magarvey, N. A.; Wagenaar, M.; Yin, X.; Greenstein, M.; Hucul, J. A.; Zabriskie, T. M. Investigating β-Hydroxyenduracididine Formation in the Biosynthesis of the Mannopeptimycins. Chem. Biol. 2005, 12 (11), 1163–1168. https://doi.org/10.1016/j.chem-biol.2005.09.013.

(9) Chang, C.-Y.; Lyu, S.-Y.; Liu, Y.-C.; Hsu, N.-S.; Wu, C.-C.; Tang, C.-F.; Lin, K.-H.; Ho, J.-Y.; Wu, C.-J.; Tsai, M.-D.; Li, T.-L. Biosynthesis of Streptolidine involved two unexpected Intermediates produced by a Dihydroxylase and a Cyclase through Unusual Mechanisms. Angew. Chem. Int. Ed. 2014, 53 (7), 1943–1948. https://doi.org/10.1002/anie.201307989.

(10) Tryon, J. H.; Rote, J. C.; Chen, L.; Robey, M. T.; Vega, M. M.; Phua, W. C.; Metcalf, W. W.; Ju, K.-S.; Kelleher, N. L.; Thomson, R. J. Genome Mining and Metabolomics Uncover a Rare D-Capreomycidine containing Natural Product and its Biosynthetic Gene Cluster. ACS Chem. Biol. 2020, 15 (11), 3013–3020. https://doi.org/10.1021/acschem-bio.0c00663.

(11) Cui, Z.; Nguyen, H.; Bhardwaj, M.; Wang, X.; Büschleb, M.; Lemke, A.; Schütz, C.; Rohrbacher, C.; Junghanns, P.; Koppermann, S.; Ducho, C.; Thorson, J. S.; Van Lanen, S. G. Enzymatic Cβ-H Functionalization of L-Arg and L-Leu in Nonribosomally Derived Peptidyl Natural Products: A Tale of Two Oxidoreductases. J. Am. Chem. Soc. 2021, 143(46), 19425–19437. https://doi.org/10.1021/jacs.1c08177.

(12) Bell, E. L.; Finnigan, W.; France, S. P.; Green, A. P.; Hayes, M. A.; Hepworth, L. J.; Lovelock, S. L.; Niikura, H.; Osuna, S.; Romero, E.; Ryan, K. S.; Turner, N. j.; Flitsch, S. L. Biocatalysis. Nat. Rev. Methods Primer 2021, 1.

(13) Pyser, J. B.; Chakrabarty, S.; Romero, E. O.; Narayan, A. R. H. State-of-the-Art Biocatalysis. ACS Cent. Sci. 2021, 7 (7), 1105–1116. https://doi.org/10.1021/acscentsci.1c00273.

(14) Lima, S. T.; Fallon, T. R.; Cordoza, J. L.; Chekan, J. R.; Delbaje, E.; Hopiavuori, A. R.; Alvarenga, D. O.; Wood, S. M.; Luhavaya, H.; Baumgartner, J. T.; Dörr, F. A.; Etchegaray, A.; Pinto, E.; McKinnie, S. M. K.; Fiore, M. F.; Moore, B. S. Biosynthesis of Guanitoxin Enables Global Environmental Detection in Freshwater Cyanobacteria. J. Am. Chem. Soc. 2022, 144 (21), 9372–9379. https://doi.org/10.1021/jacs.2c01424.

(15) Fiore, M. F.; de Lima, S. T.; Carmichael, W. W.; McKinnie, S. M. K.; Chekan, J. R.; Moore, B. S. Guanitoxin, Re-Naming a Cyanobacterial Organophosphate Toxin. Harmful Algae 2020, 92, 101737. https://doi.org/10.1016/j.hal.2019.101737.

(16) Carmichael, W. W. Health Effects of Toxin-Producing Cyanobacteria: “The CyanoHABs.” Hum. Ecol. Risk Assess. Int. J. 2001, 7 (5), 1393–1407. https://doi.org/10.1080/20018091095087.

(17) Matsunaga, S.; Moore, R. E.; Niemczura, W. P.; Carmichael, W. W. Anatoxin-a(*s*), a Potent Anticholinesterase from *Anabaena flos-aquae*. J. Am. Chem. Soc. 1989, 111(20), 8021–8023. https://doi.org/10.1021/ja00202a057.

(18) Hemscheidt, T.; Burgoyne, D. L.; Moore, R. E. Biosynthesis of Anatoxin-a(*s*). (2*S*,4*S*)-4-Hydroxyarginine as an Intermediate. J. Chem. Soc. Chem. Commun. 1995, No. 2, 205–206. https://doi.org/10.1039/c39950000205.

(19) Du, Y.-L.; Ryan, K. S. Pyridoxal Phosphate-Dependent Rections in the Biosynthesis of Natural Products. Nat. Prod. Rep. 2019, 36 (3), 430–457. https://doi.org/10.1039/C8NP00049B.

(20) Rocha, J. F.; Pina, A. F.; Sousa, S. F.; Cerqueira, N. M. F. S. A. PLP-Dependent Enzymes as Important Biocatalysts for the Pharmaceutical, Chemical and Food Industries: A Structural and Mechanistic Perspective. Catal. Sci. Technol. 2019, 9 (18), 4864–4876. https://doi.org/10.1039/C9CY01210A.

(21) Burroughs, A. M.; Hoppe, R. W.; Goebel, N. C.; Sayyed, B. H.; Voegtline, T. J.; Schwabacher, A. W.; Zabriskie, T. M.; Silvaggi, N. R. Structural and Functional Characterization of MppR, an Enduracididine Biosynthetic Enzyme from *Streptomyces hygroscopicus:* Functional Diversity in the Acetoacetate Decarboxylase-like Superfamily. Biochemistry 2013, 52 (26), 4492–4506. https://doi.org/10.1021/bi400397k.

(22) Han, L.; Schwabacher, A. W.; Moran, G. R.; Silvaggi, N. R. S. *wadayamensis* MppP Is a Pyridoxal 5’-Phosphate-Dependent L-Arginine α-Deaminase, γ-Hydroxylase in the Enduracididine Biosynthetic Pathway. Biochemistry 2015, 54(47), 7029–7040. https://doi.org/10.1021/acs.bio-chem.5b01016.

(23) Han, L.; Vuksanovic, N.; Oehm, S. A.; Fenske, T. G.; Schwabacher, A. W.; Silvaggi, N. R. *Streptomyces wadayamensis* MppP Is a PLP-Dependent Oxidase, not an Oxygenase. Biochemistry 2018, 57 (23), 3252–3264. https://doi.org/10.1021/acs.biochem.8b00130.

(24) Vuksanovic, N.; Serrano, D. A.; Schwabacher, A. W.; Silvaggi, N. R. Structural and Preliminary Biochemical Characterization of MppQ, a PLP-Dependent Aminotransferase from *Streptomyces hygroscopicus*; preprint; bioRxiv, 2022. https://doi.org/10.1101/2022.04.03.486910.

(25) Ju, J.; Ozanick, S. G.; Shen, B.; Thomas, M. G. Conversion of (2*S*)-Arginine to (2*S*,3*R*)-Capreomycidine by VioC and VioD from the Viomycin Biosynthetic Pathway of *Streptomyces* sp. Strain ATCC11861. ChemBioChem 2004, 5 (9), 1281–1285. https://doi.org/10.1002/cbic.200400136.

(26) Yin, X.; McPhail, K. L.; Kim, K.; Zabriskie, T. M. Formation of the Nonproteinogenic Amino Acid 2*S*,3*R*-Capre-omycidine by VioD from the Viomycin Biosynthesis Pathway. ChemBioChem 2004, 5 (9), 1278–1281. https://doi.org/10.1002/cbic.200400187.

(27) Clausen, T.; Huber, R.; Prade, L.; Wahl, M. C.; Messerschmidt, A. Crystal Structure of *Escherichia coli* Cystathionine γ-Synthase at 1.5 Å Resolution. EMBO J. 1998, 17(23), 6827–6838. https://doi.org/10.1093/em-boj/17.23.6827.

(28) Chen, M.; Liu, C.-T.; Tang, Y. Discovery and Biocatalytic Application of a PLP-Dependent Amino Acid γ-Substitution Enzyme that Catalyzes C–C Bond Formation. J. Am. Chem. Soc. 2020, 142 (23), 10506–10515. https://doi.org/10.1021/jacs.0c03535.

(29) Hai, Y.; Chen, M.; Huang, A.; Tang, Y. Biosynthesis of Mycotoxin Fusaric Acid and Application of a PLP-Dependent Enzyme for Chemoenzymatic Synthesis of Substituted L-Pipecolic Acids. J. Am. Chem. Soc. 2020, 142 (46), 19668–19677. https://doi.org/10.1021/jacs.0c09352.

(30) Yee, D. A.; Niwa, K.; Perlatti, B.; Chen, M.; Li, Y.; Tang, Y. Genome Mining for Unknown–Unknown Natural Products. Nat. Chem. Biol. 2023. https://doi.org/10.1038/s41589-022-01246-6.

(31) Cui, Z.; Overbay, J.; Wang, X.; Liu, X.; Zhang, Y.; Bhardwaj, M.; Lemke, A.; Wiegmann, D.; Niro, G.; Thorson, J. S.; Ducho, C.; Van Lanen, S. G. Pyridoxal-5’-Phosphate-Dependent Alkyl Transfer in Nucleoside Antibiotic Biosynthesis. Nat. Chem. Biol. 2020, 16 (8), 904–911. https://doi.org/10.1038/s41589-020-0548-3.

(32) Hauth, F.; Buck, H.; Stanoppi, M.; Hartig, J. S. Canavanine Utilization via Homoserine and Hydroxyguanidine by a PLP-Dependent γ-Lyase in *Pseudomonadaceae* and *Rhizobiales*. RSC Chem. Biol. 2022, 3 (10), 1240–1250. https://doi.org/10.1039/D2CB00128D.

(33) Giltrap, A. M.; Dowman, L. J.; Nagalingam, G.; Ochoa, J. L.; Linington, R. G.; Britton, W. J.; Payne, R. J. Total Synthesis of Teixobactin. Org. Lett. 2016, 18 (11), 2788–2791. https://doi.org/10.1021/acs.orglett.6b01324.

(34) Montioli, R.; Fargue, S.; Lewin, J.; Zamparelli, C.; Danpure, C. J.; Borri Voltattorni, C.; Cellini, B. The N-Terminal Extension Is Essential for the Formation of the Active Dimeric Structure of Liver Peroxisomal Alanine: Glyoxylate Aminotransferase. Int. J. Biochem. Cell Biol. 2012, 44 (3), 536–546. https://doi.org/10.1016/j.biocel.2011.12.007.

(35) Edayathumangalam, R.; Wu, R.; Garcia, R.; Wang, Y.; Wang, W.; Kreinbring, C. A.; Bach, A.; Liao, J.; Stone, T. A.; Terwilliger, T. C.; Hoang, Q. Q.; Belitsky, B. R.; Petsko, G. A.; Ringe, D.; Liu, D. Crystal Structure of *Bacillus subtilis* GabR, an Autorepressor and Transcriptional Activator of *GabT*. Proc. Natl. Acad. Sci. U. S. A. 2013, 110 (44), 17820–17825. https://doi.org/10.1073/pnas.1315887110.

(36) Brzovic, P.; Holbrook, E. L.; Greene, R. C.; Dunn, M. F. Reaction Mechanism of *Escherichia coli* Cystathionine γ-Synthase: Direct Evidence for a Pyridoxamine Derivative of Vinylgloxylate as a Key Intermediate in Pyridoxal Phosphate Dependent γ–Elimination and γ-Replacement Reactions. Biochemistry 1990, 29 (2), 442–451. https://doi.org/10.1021/bi00454a020.

(37) Drewe, W. F.; Dunn, M. F. Characterization of the Reaction of L-Serine and Indole with *Escherichia coli* Tryptophan Synthase via Rapid-Scanning Ultraviolet-Visible Spectroscopy. Biochemistry 1986, 25 (9), 2494–2501. https://doi.org/10.1021/bi00357a032.

(38) Herger, M.; van Roye, P.; Romney, D. K.; Brinkmann-Chen, S.; Buller, A. R.; Arnold, F. H. Synthesis of β-Branched Tryptophan Analogues Using an Engineered Subunit of Tryptophan Synthase. J. Am. Chem. Soc. 2016, 138(27), 8388–8391. https://doi.org/10.1021/jacs.6b04836.

(39) Kumar, P.; Meza, A.; Ellis, J. M.; Carlson, G. A.; Bingman, C. A.; Buller, A. R. L-Threonine Transaldolase Activity is Enabled by a Persistent Catalytic Intermediate. ACS Chem. Biol. 2021, 16 (1), 86–95. https://doi.org/10.1021/acschem-bio.0c00753.

(40) Phillips, R. S.; Sundararaju, B.; Koushik, S. V. The Catalytic Mechanism of Kynureninase from *Pseudomonas fluorescens*: Evidence for Transient Quinonoid and Ketimine Intermediates from Rapid-Scanning Stopped-Flow Spectrophotometry. Biochemistry 1998, 37 (24), 8783–8789. https://doi.org/10.1021/bi980066v.

(41) Ronda, L.; Bazhulina, N. P.; Morozova, E. A.; Revtovich, S. V.; Chekhov, V. O.; Nikulin, A. D.; Demidkina, T. V.; Mozzarelli, A. Exploring Methionine γ-Lyase Structure-Function Relationship via Microspectrophotometry and X-Ray Crystallography. Biochim. Biophys. Acta, Proteins Proteomics 2011, 1814 (6), 834–842. https://doi.org/10.1016/j.bbapap.2010.06.017.

(42) Chun, S. W.; Narayan, A. R. H. Biocatalytic, Stereoselective Deuteration of α-Amino Acids and Methyl Esters. ACS Catal. 2020, 10 (13), 7413–7418. https://doi.org/10.1021/acscatal.0c01885.

(43) Doyon, T. J.; Buller, A. R. Site-Selective Deuteration of Amino Acids through Dual-Protein Catalysis. J. Am. Chem. Soc. 2022, 144 (16), 7327–7336. https://doi.org/10.1021/jacs.2c00608.

(44) Liang, J.; Han, Q.; Tan, Y.; Ding, H.; Li, J. Current Advances on Structure-Function Relationships of Pyridoxal 5’-Phosphate-Dependent Enzymes. Front. Mol. Biosci. 2019, 6, 4. https://doi.org/10.3389/fmolb.2019.00004.

(45) Okamoto, A.; Higuchi, T.; Hirotsu, K.; Kuramitsu, S.; Kagamiyama, H. X-Ray Crystallographic Study of Pyridoxal 5’-Phosphate-Type Aspartate Aminotransferases from *Escherichia coli* in Open and Closed Form. J. Biochem. 1994, 116 (1), 95–107.

(46) Jumper, J.; Evans, R.; Pritzel, A.; Green, T.; Figurnov, M.; Ronneberger, O.; Tunyasuvunakool, K.; Bates, R.; Žídek, A.; Potapenko, A.; Bridgland, A.; Meyer, C.; Kohl, S. A. A.; Ballard, A. J.; Cowie, A.; Romera-Paredes, B.; Nikolov, S.; Jain, R.; Adler, J.; Back, T.; Petersen, S.; Reiman, D.; Clancy, E.; Zielinski, M.; Steinegger, M.; Pacholska, M.; Berghammer, T.; Bodenstein, S.; Silver, D.; Vinyals, O.; Senior, A. W.; Kavukcuoglu, K.; Kohli, P.; Hassabis, D. Highly Accurate Protein Structure Prediction with AlphaFold. Nature 2021, 596 (7873), 583–589. https://doi.org/10.1038/s41586-021-03819-2.

(47) Tang, Z.; Tang, H.; Wang, W.; Xue, Y.; Chen, D.; Tang, W.; Liu, W. Biosynthesis of a New Fusaoctaxin Virulence Factor in *Fusarium graminearum* Relies on a Distinct Path to Form a Guanidinoacetyl Starter Unit Priming Nonribosomal Octapeptidyl Assembly. J. Am. Chem. Soc. 2021, 143 (47), 19719–19730. https://doi.org/10.1021/jacs.1c07770.

(48) Fogle, E. J.; Liu, W.; Woon, S.-T.; Keller, J. W.; Toney, M. D. Role of Q52 in Catalysis of Decarboxylation and Transamination in Dialkylglycine Decarboxylase. Biochemistry 2005, 44 (50), 16392–16404. https://doi.org/10.1021/bi051475b.

(49) Marchand, J. A.; Neugebauer, M. E.; Ing, M. C.; Lin, C.-I.; Pelton, J. G.; Chang, M. C. Y. Discovery of a Pathway for Terminal-Alkyne Amino Acid Biosynthesis. Nature 2019, 567 (7748), 420–424. https://doi.org/10.1038/s41586-019-1020-y.

(50) Barra, L.; Awakawa, T.; Shirai, K.; Hu, Z.; Bashiri, G.; Abe, I. β-NAD as a Building Block in Natural Product Biosynthesis. Nature 2021, 600 (7890), 754–758. https://doi.org/10.1038/s41586-021-04214-7.

(51) Chun, S. W.; Hinze, M. E.; Skiba, M. A.; Narayan, A. R. H. Chemistry of a Unique Polyketide-like Synthase. J. Am. Chem. Soc. 2018, 140 (7), 2430–2433. https://doi.org/10.1021/jacs.7b13297.

(52) Kawata, J.; Naoe, T.; Ogasawara, Y.; Dairi, T. Biosynthesis of the Carbonylmethylene Structure Found in the Ketomemicin Class of Pseudotripeptides. Angew. Chem. Int. Ed. 2017, 56 (8), 2026–2029. https://doi.org/10.1002/anie.201611005.

(53) Ellis, J. M.; Campbell, M. E.; Kumar, P.; Geunes, E. P.; Bingman, C. A.; Buller, A. R. Biocatalytic Synthesis of Non-Standard Amino Acids by a Decarboxylative Aldol Reaction. Nat. Catal. 2022, 5 (2), 136–143. https://doi.org/10.1038/s41929-022-00743-0.

(54) Hoffarth, E. R.; Caddell Haatveit, K.; Kuatsjah, E.; MacNeil, G. A.; Saroya, S.; Walsby, C. J.; Eltis, L. D.; Houk, K. N.; Garcia-Borràs, M.; Ryan, K. S. A Shared Mechanistic Pathway for Pyridoxal Phosphate–Dependent Arginine Oxidases. Proc. Natl. Acad. Sci. U. S. A. 2021, 118 (40), e2012591118 (1-10). https://doi.org/10.1073/pnas.2012591118.

