## Supplementary material for "Mechanistic and structural insights into a divergent PLP-dependent L-enduracididine cyclase from a toxic cyanobacterium": GntC-SI-PDF

**This PDF file includes:**

General Materials and Methods
Figures S1 to S21
Tables S1 to S4
Chemical Synthesis
NMR and Compound Characterization
References

### Table of Contents

1. **General Methods and Materials** **3**
2. **Molecular Biology and Biochemical Methods** **4**
3. **Enzyme Assay Methods** **7**
4. **Protein Crystallography Methods** **8**
5. **Supplementary Figures** **10**
   Fig. S1. Previously reported *in vivo* guanitoxin stable isotope labeling results. 10
   Fig. S2. *In vitro* GntC PLP dependency assay. 11
   Fig. S3. *In vitro* GntC pH preference assay. 12
   Fig. S4. *In vitro* GntC divalent metal cation preference assay. 13
   Fig. S5. *In vitro* GntC assays with N- or C-terminally $His_6$-tagged enzyme. 14
   Fig. S6. UV-Vis spectra of GntC assays with substrate **1** and diastereomer **1′**. 15
   Fig. S7. *In vitro* $^1$H NMR GntC assay with substrate **1** in $H_2O$ and $D_2O$. 16
   Fig. S8. *In vitro* $^1$H NMR GntC assay with substrate diastereomer **1′** in $D_2O$. 17
   Fig. S9. *In vitro* GntC assay with product L-enduracididine (**2**) in $D_2O$. 18
   Fig. S10. *In vitro* GntC assay to assess overall reaction reversibility. 19
   Fig. S11. *In vitro* GntC assay in 50% $H_2{}^{18}O$ to assess dehydration reversibility. 20
   Fig. S12. Sequence alignment of GntC to PLP-dependent capreomycidine cyclases. 21
   Fig. S13. Sequence alignment of GntC to PLP-dependent intermolecular γ-substitution enzymes and aspartate aminotransferase AspC. 22
   Fig. S14. Structural comparison of GntC with PLP and PLP-**1** substrates. 23
   Fig. S15. Active site comparison of PLP-**1** GntC to aspartate aminotransferase. 24
   Fig. S16. Active site comparison of PLP-**1** GntC and VioD AlphaFold model. 25
   Fig. S17. Active site comparison of PLP-**1** GntC and Fgm3 AlphaFold model. 26
   Fig. S18. Analytical size exclusion chromatograms of wild-type GntC and mutants. 27
   Fig. S19. Proposed rationale for how GntC N52* enhances γ-hydroxy elimination. 28
   Fig. S20. OrfR capreomycidine cyclase is mechanistically distinct from GntC. 29
   Fig. S21. PLP-dependent intermolecular γ-substitution reactions from diverse biosynthetic pathways are functionally similar to the intramolecular GntC reaction. 30
6. **Tables** **32**
   Table S1. NCBI accession codes for sequences used in this study 32
   Table S2. Data collection and refinement statistics of GntC 33
   Table S3. Residues modeled in each chain of GntC (residues 1-370) 34
   Table S4. GntC primers used in this study 35
7. **Chemical Synthesis** **36**
   Synthesis of **1**-$d_\gamma$ 36
8. **NMR and Compound Characterization** **38**
   NMR spectra for: SI-1 38
   SI-2 41
   **1**-$d_\gamma$ 45
9. **References** **48**

### General Methods and Materials

#### Synthesis and analytical methods

For chemical synthesis, solvents were bought from Thermo Fisher Scientific and Millipore Sigma, and reagents were used without further purification unless otherwise noted. Anhydrous reactions were performed in oven-dried glassware under an argon atmosphere. Reactions were monitored via precoated plates thin-layer chromatography (TLC, Merck, 60 $F_{254}$), and visualized via $KMnO_4$ staining solution (1.5 g $KMnO_4$, 10 g $K_2CO_3$, 1.25 mL 10% NaOH in 200 mL water) and with UV at 254 nm, with TLC $R_f$ values rounded to the nearest 0.05. Silica gel (Alfa Aesar, 60 (215-400 mesh)) was used to purify products via flash column chromatography, and rotary evaporation was used to concentrate samples under reduced pressure. Nuclear magnetic resonance (NMR) spectra were obtained using an Avance III HD spectrometer (Bruker) equipped with a BBFO SmartProbe at 500 MHz ($^1$H NMR) or 125 MHz ($^{13}$C NMR) using $CDCl_3$ or $D_2O$ as solvents. Chemical shifts for synthetic reactions are reported in ppm ($\delta$) and referenced either to methanol as an internal standard for $D_2O$ ($\delta$=4.79 ppm for $^1$H NMR), or the $CDCl_3$ solvent signal ($\delta$=7.26 ppm for $^1$H, $\delta$=77.2 ppm for $^{13}$C NMR). NMR data are reported as follows: s=singlet, d=doublet, t=triplet, q=quartet, m=multiplet, J=coupling constant (Hz). High resolution mass spectrometer (HRMS) data were obtained via MALDI-qTOF on a timsTOFfleX (Bruker) in positive mode with a laser power of 50%, frequency of 1000 Hz, and 1000 shots per sample.

#### LCMS method

General LCMS conditions were followed from a literature reference.[1] In summary, assays were measured on an Elute UHPLC system (Bruker) coupled to an ESI-ion trap (Bruker) mass spectrometer in positive mode. Compound separation was achieved using reversed-phase chromatography using the eluents water+0.1% formic acid (Solvent A) and acetonitrile+0.1% formic acid (Solvent B) with a Bruker Intensity Solo C18(2), 2 mm – 2 x 100 mm column. The LC method is adapted from a literature reference,[1] with the modification of a hold at 10% B for 4 minutes.

#### Reagents for $D_2O$ assays

Premade stock solutions of pyridoxal-5′-phosphate (PLP) and $K_2HPO_4$ buffer were lyophilized overnight, then resuspended in $D_2O$ and lyophilized a second time. Solutions were washed and lyophilized in $D_2O$ an additional time before being stored under Ar gas at -20 °C until needed.

#### General Marfey's derivatization for LCMS analysis

The reaction procedure for Marfey's derivatization of enzymatic assays was followed from a previous reference.[1] Briefly, saturated sodium bicarbonate (20 $\mu$L) was added to the enzyme assay (50 $\mu$L). A 1% w/v solution of Marfey's reagent was prepared in acetone, and 100 $\mu$L of the Marfey's solution was added to the crude enzyme assay to begin the derivatization reaction. After 90 minutes of incubation at 37 °C, the reaction was neutralized with 1N HCl (25 $\mu$L). The neutralized reaction was centrifuged at 14,000 *x g* for 10 minutes, and the supernatant was extracted for LCMS analysis.

### Molecular Biology and Biochemical Methods

#### *gntC* cloning

*gntC* C-terminus $His_6$ tag cloning: Cloning of *gntC* with a C-terminal $His_6$ tag was achieved with the primers listed in Table S3 with HiFi Assembly polymerase mix (New England Biolabs Inc.) with the following amplification conditions to amplify the gene insert:

Initial denaturation at 98 °C (1:40 minutes), 30 cycles of 98 °C (10 seconds), 60 °C (10 seconds), 72 °C (1:40 minutes), a final elongation of 72 °C (3 minutes) and hold at 4 °C until removal of product from PCR thermocycler.

To prepare linearized pET28a(+) backbone, a restriction enzyme digest was performed on empty pET28a(+) with the restriction enzymes NcoI and XhoI at 37 °C for 30 minutes, then 80 °C for 20 minutes to heat inactivate the restriction enzymes.

Once the gene insert and linear pET28a(+) were prepared, a Gibson assembly was performed and incubated at 55 °C for 30 minutes, and transformed into *E. coli* DH10β cells. Sequencing of *gntC* was confirmed via Sanger sequencing (Azenta Life Sciences).

*gntC* mutant cloning: Using the pET28a(+) plasmid of WT *gntC* as a template, site-directed mutagenesis of *gntC* was achieved by PCR using the primers listed in Table S3 with Q5 polymerase (New England Biolabs Inc.), and the following amplification conditions:

General PCR protocol for each primer set: initial denaturation at 98 °C (30 seconds), 35 cycles of 98 °C (5 seconds), 68 °C (10 seconds),* and 72 °C (1:40 minutes), and a final elongation of 72 °C (1:30 minutes).
*temperature varied based on melting temperature of primer set, further details listed in Table S3

PCR products were run on a 10% agarose gel to check for amplification prior to reaction with KLD (kinase, ligase, DpnI) enzyme reaction mix (New England Biolabs Inc.) to remove parental DNA. After the reaction with the KLD mixture, the reaction product was transformed into E. coli DH10β cells. Sequencing of mutants were confirmed via Sanger sequencing (Azenta Life Sciences).

#### Transformation into *E. coli*

Transformation protocol was adapted from work previously done in the lab.[1] Plasmids used in this study were transformed into *E. coli* DH10β chemically competent cells for plasmid storage and BL21(DE3) cells for plasmid expression by heat shock. After thawing cell aliquots from -80 °C storage on ice for 10 minutes, 1.0 μL of plasmid was added to cell aliquots and incubated on ice for 30 minutes. Cells were heat shocked at 42 °C for 55 seconds, then incubated on ice again for 2 minutes. Aliquots were added to 750 μL of LB media and then incubated at 37 °C at 200 rpm for 1 hour. Afterword, cells were plated on LB media plates with 50 mg/mL kanamycin and incubated at 37 °C overnight, then checked for colony formation the next day.

#### Plasmid purification

Plasmid purification was achieved using the QIAprep Spin Miniprep Kit (QIAGEN) with the manufacturer's protocol. DNA concentrations were measured via NanoDrop.

**GntC heterologous protein expression**

A general method from a literature reference[1] was followed for GntC and each GntC mutant. After sequence confirmation, purified plasmid was transformed into *E. coli* DH10β and BL21(DE3) cells. From these cultures, glycerol stocks were prepared, and a colony was picked from the *E. coli* BL21(DE3) plate for protein expression starter cultures for the GntC mutants. Starter cultures contained 15 mL LB media, 50 mg/mL kanamycin, and a colony of the appropriate GntC mutant, and were incubated overnight at 37 °C at 200 rpm. After overnight incubation, 10 mL of the starter culture was inoculated into 1 L of Terrific Broth media containing 50 mg/mL kanamycin and was incubated at 37 °C and 200 rpm until the culture reached an $OD_{600}$ of 0.8. Afterword, the 1 L culture was cooled to 18 °C with 200 rpm for 1 hour prior to the addition of 100 μM isopropyl-β-D-thiogalactopyranoside (IPTG) to induce protein expression. Cultures were incubated overnight at 18 °C and 200 rpm prior to cell harvesting. Cultures were harvested by centrifugation at 2500 x g at 4 °C for 30 minutes, then resuspended in 30 mL cell lysis buffer (1 M NaCl, 20 mM Tris-HCl pH 8.0, 20 mM imidazole, 10% glycerol) and stored at -80 °C until protein purification.

**GntC protein purification**

Protein purification conditions adapted from a literature reference.[1] *E. coli* BL21 (DE3) cell pellets containing $(His)_6$-*gntC* variants were thawed, then sonicated on ice for a total of 6 minutes at 40% amplitude with 15 seconds on, and 45 seconds off (FisherBrand Model 505 Sonic Dismembrator, 3.2 mm microtip). The lysate was clarified by centrifugation at 15,000 *x g* for 30 minutes at 4 °C. Proteins were purified at 4 °C using an AKTAGo FPLC system with a HisTrap FF (5 mL) column (GE Healthcare Life Sciences) preequilibrated with at least 25 mL Buffer A (30 mM Tris pH 8.0, 30 mM imidazole, 300 mM NaCl) at a maximum flow rate of 2 mL/minute. Buffers used for FPLC purification were filtered through a 0.22 μM nitrocellulose membrane prior to usage. After loading the clarified cell lysate onto the column, the column was rinsed with Buffer A until the UV absorbance returned to baseline, then the column was washed with 10% Buffer B (20 mM Tris pH 8.0, 1 M NaCl, and 250 imidazole) to remove non-specifically bound proteins using either 25 mL Buffer B or until the UV absorbance returned to baseline. $His_6$-GntC was eluted with a linear gradient to 100% B over 60 mL in 30 minutes, while collecting 5 mL fractions. The eluate fractions were assessed for purity using a 10% SDS-PAGE gel, and fractions with over 90% GntC, were combined and concentrated to less than 2.5 mL using by Amicon ultra centrifugal filters (30 kDa molecular weight cut-off (MWCO), EMD-Millipore). Proteins were buffer exchanged into GF Buffer (50 mM HEPES pH 8.0, 300 mM KCl) using a preequilibrated PD-10 column (GE Healthcare Life Sciences). Protein concentrations were estimated using the Bradford assay with a bovine serum albumin standard and, if necessary, protein was further concentrated with the 30 kDa Amicon ultra centrifugal filter. After obtaining purified protein, the protein was aliquoted into 50 μL aliquots and stored at -80 °C.

The following pure protein concentrations were obtained: WT GntC (37 mg/L), GntC N52A (9 mg/L), GntC N52Q (16 mg/L), GntC N52D (13 mg/L), GntC T84A (8 mg/L), GntC T84V (6 mg/L), GntC K219A (31 mg/L), GntC S25A* (4 mg/L).

*During FPLC elution, protein eluted with the absence of PLP and appeared to aggregate in a concentration-dependent manner, in which 500 μL of 2.00 mM PLP was added to each fraction to stabilize the enzyme. After elution, aggregate was removed via centrifugation at 20,000 *x g* at 4 °C for 10 minutes to remove aggregates.

**Protein oligomerization assessment**

To assess oligomerization of site-directed GntC variants, a Superdex 200 size exclusion column (GE Healthcare Life Sciences) was equilibrated with 40 mL of GF buffer. Protein samples were diluted to 4 mg/mL, and 60 μL was directly injected to the FPLC. Using a flow rate of 0.20 mL/minute, fractions were eluted, and the main dimer peak was collected.

**GntC buffer exchange for $D_2O$ assays**

Enzyme buffer exchange procedures were adapted from a literature reference.[2] A 500 μL aliquot of GntC was thawed on ice from storage at -80 °C. The aliquot would be pipetted into a 30 kDa MWCO Amicon centrifugal filter with 50 mM $K_2HPO_4$ at pH 8.0 (5 mL) in $D_2O$ and centrifuged at 3,400 *x g* for 20 minutes. The GntC aliquot would be washed with two additional $K_2HPO_4$ $D_2O$ washes prior to Bradford assay to determine enzyme concentration.

**GntC apoenzyme preparation**

GntC coelutes with the cofactor PLP during protein purification. To obtain apo-GntC, a 1 mL aliquot of GntC was thawed on ice from storage at -80 °C. GntC was added to GF buffer and 5 mM hydroxylamine to a volume of 10 mL. The solution was incubated on ice for 4 hours prior to adding to a 30 kDa MWCO Amicon centrifugal filter with GF buffer and centrifuged at 3,400 *x g* for 20 minutes. The solution was centrifuged with two additional GF buffer washes prior to Bradford assay to determine enzyme concentration.

### Enzyme Assay Methods

#### $H_2O$ enzyme assays

Wild-type (WT) and mutant GntC assays were conducted in a similar manner to previously reported.[1] Assays were conducted in 50 mM $K_2HPO_4$ (pH 8.0), using 1 mM substrate, 100 μM PLP, and 50 μM purified GntC enzyme. Total reaction volumes were brought to a total volume of 50 μL using MilliQ water and incubated at room temperature overnight. After incubation, reactions were derivatized using the detailed protocol above for Marfey's derivatization, then subjected to LCMS analysis.

#### Apoenzyme GntC assay

GntC PLP-stripped assays were conducted in 50 mM $K_2HPO_4$ (pH 8.0) using 1 mM substrate, and 50 μM apoenzyme GntC. Reaction volumes were brought to 50 μL with MilliQ water and incubated overnight at room temperature. Reactions with derivatized with Marfey's reagent (protocol detailed above) prior to LCMS analysis.

#### $D_2O$ enzyme assays

GntC deuterium incorporation assays were conducted in 50 mM $K_2HPO_4$ (pH 8.0) using 1 mM substrate, 100 μM PLP, and 38 μM of purified GntC enzyme that was buffer exchanged into $D_2O$ and 50 mM $K_2HPO_4$. Reaction volumes were brought to their total volume of 50 μL with $D_2O$. Assays were incubated at room temperature overnight, then derivatized with Marfey's reagent prior to LCMS analysis.

#### $H_2{}^{18}O$ enzyme assays

To help establish the irreversibility of a step of the GntC mechanism, GntC assays were conducted in 50% $H_2{}^{18}O$ (Cambridge Isotope Laboratories) by using 50 mM $K_2HPO_4$ (pH, 8.0), 1 mM substrate, 100 μM PLP, and 50 μM GntC. All reagents were stored in MilliQ water, and enzyme assays were diluted to a final volume of 50 μL using $H_2{}^{18}O$. After 16 hours of incubation at room temperature, the reactions were derivatized with Marfey's reagent for LCMS analysis.

#### NMR enzyme assays

GntC NMR assays were conducted in 50 mM $K_2HPO_4$ (pH 8.0), 2.6 mM substrate (unless otherwise stated), 100 μM PLP, and 30 μM purified WT GntC. GntC enzyme was buffer exchanged into $D_2O$ using the protocol above. Reaction volumes were brought to 500 μL with $D_2O$. Reactions placed into a clean, oven-dried NMR tube and analyzed using $^1H$ NMR obtained on a Bruker 500 MHz NMR instrument using the default settings except for using 256 scans rather than 16 scans.

#### Spectroscopy assays

GntC UV-Vis assays were adapted from a literature reference.[3] Assays were conducted in 50 mM $K_2HPO_4$ (pH 8.0), 1 mM substrate, and 40 μM purified WT GntC enzyme. No additional PLP was added to the reaction, and reaction volumes were brought to 500 μL using MilliQ water. Assays were analyzed in a 500 μL, 10 mm quartz cuvette cell (VWR International, LLC) using an Eppendorf BioSpectrometer between 300 and 550 nm.

### Protein Crystallography Methods

#### Protein expression and purification for crystallography experiments

For crystallography studies, GntC was expressed and purified in a similar manner above with the following changes: A single colony was inoculated into a 10 mL of LB starter culture, grown at 37 °C and 220 rpm shaking overnight. The starter culture was then inoculated into 1 L LB media, grown at 37 °C and 220 rpm shaking until $OD_{600}$ reaches 1.0. An hour after the temperature adjustment, the protein expression was induced by IPTG at 1 mM final concentration. Cells were harvested by centrifugation (8,000 x g, 4 °C, 10 min) 16 hours after induction.

For protein purification the following changes were made from the protocol above: The column was further equilibrated with 40 mL of Buffer A (25 mM Tris pH 8.0, 300 mM NaCl, 10 mM imidazole, 100 μM PLP). The column was eluted with a linear gradient to 100% Buffer B (25 mM Tris pH 8.0, 300 mM NaCl, and 250 mM imidazole) over 40 mL volume while collecting 5 mL fractions. The protein was further purified on a Superdex 75 16/60 size exclusion column (GE Healthcare Life Sciences) in the gel filtration buffer (100 mM MOPS pH 6.7 and 20 μM PLP). The dimeric peak from gel filtration was collected and concentrated by Amicon ultra centrifugal filters (30 kD molecular weight cut-off) to a final concentration of 10.6 mg/mL. The protein was flash-frozen in 25 μL aliquots and stored in -80 °C freezer.

#### Crystallization of GntC and cocrystallization of GntC with 1

GntC was crystallized by hanging drop crystallization at 8 °C. 1.0 μL of 10.6 mg/mL GntC in the gel filtration buffer was mixed with 1.0 μL well solution (19% (w/v) PEG 3350, 0.50 M $MgCl_2$, 0.10 M Tris pH 8.5, and 1 mM PLP) to make a 2-μL hanging drop in a sealed well with 500 μL well solution. Yellow rod crystals grew overnight. The crystals were then transferred to a cryogenic solution (19% (w/v) PEG 3350, 0.50 M $MgCl_2$, 0.10 M Tris pH 8.5, 1 mM PLP, and 15 % (v/v) ethylene glycol) and flash-cooled in liquid nitrogen.

Purified **1** was dissolved in water and titrated with 1 M NaOH to a final concentration of 100 mM at approximately pH 7. GntC with **1** bound was crystallized by sitting drop crystallization at 4 °C. 1.0 μL of 9.6 mg/mL GntC in the gel filtration buffer supplemented with 10 mM **1** was mixed with 1.0 μL well solution (24-27% (w/v) PEG 3350, 0.40-0.44 M $MgCl_2$, 0.10 M Tris pH 8.5, and 1 mM PLP) to make a 2-μL sitting drop in a sealed well with 500 μL well solution. Yellow rod crystals grew overnight. The crystals were then transferred to a cryogenic solution (24% (w/v) PEG 3350, 0.44 M $MgCl_2$, 0.10 M Tris pH 8.5, and 20% (v/v) ethylene glycol) and flash-cooled in liquid nitrogen.

#### Data collection and processing

A data set of GntC was collected at Advanced Light Source (Berkeley, California, USA) on beamline 8.2.1 using a ADSC Q315R detector at a temperature of 100 K. Resolution cutoffs were chosen based on $I/\sigma$ limit of 2. Data were indexed, integrated, and scaled in AutoProc.[4]

A data set of GntC with **1** bound was collected at Stanford Synchrotron Radiation Lightsource (Menlo Park, California, USA) on beamline 12-2 using a Pilatus 6M PAD detector at a temperature of 100 K. Resolution cutoffs were chosen as $CC_{1/2} \sim 0.7$. Data were indexed, integrated, and scaled in XDS.[5] The crystal of GntC with **1** bound was determined by Phenix[6] Xtriage to have merohedral twinning, with a twin law of h, - k, - l. The twin fraction was refined in Phenix Refine to 48%, and the structure was refined to account for twinning.

#### Structure determination and refinement

The structure of GntC was determined to 2.10-Å resolution by molecular replacement (MR) using MRage[7] implemented in Phenix. A PLP-dependent aminotransferase (ZP_03625122.1) from *Streptococcus suis* 89/1591 (PDB ID: 3OP7), which shared 28.9% identity with GntC, was trimmed by Sculptor[8] and used as a model for molecular replacement. The MR gives one solution containing two GntC dimers per asymmetric unit (ASU) with a log-likelihood gain (LLG) value of 655 and a final translational function Z score (TFZ) of 20.6. The MR solution was used as an input for AutoBuild[9] implemented in Phenix for initial model building. After the AutoBuild run, the atomic coordinates and B-factors were iteratively refined in Phenix Refine with model building and manual adjustment of model in Coot.[10] TLS parameters were refined in the last few rounds of refinement in which each GntC monomer is a TLS group. Water molecules were added manually throughout real space refinements using $F_o$-$F_c$ electron density contoured to 3.0$\sigma$ as criteria. Four-fold non-crystallographic symmetry (NCS) restraints were used throughout refinement. Restraints for Lys-PLP adduct were generated from Grade Web Server (Global Phasing).

The structure of GntC with **1** bound was determined to 2.04-Å resolution by rigid-body refinement from 2.10-Å resolution GntC structure. The atomic coordinates and B-factors was iteratively refined in Phenix Refine with model building and manual adjustment of model in Coot. Water molecules were added manually throughout real space refinements using $F_o$-$F_c$ electron density contoured to 3.0$\sigma$ as criteria. Four-fold non-crystallographic symmetry (NCS) restraints were used throughout refinement. Restraints for the PLP-**1** adduct were generated from Grade Web Server (Global Phasing). A composite-omit electron density map calculated by Phenix Composite_omit_map was used to verify the model. Restraints for Lys-PLP adduct were generated from Grade Web Server (Global Phasing).

### Supplementary figures and tables:

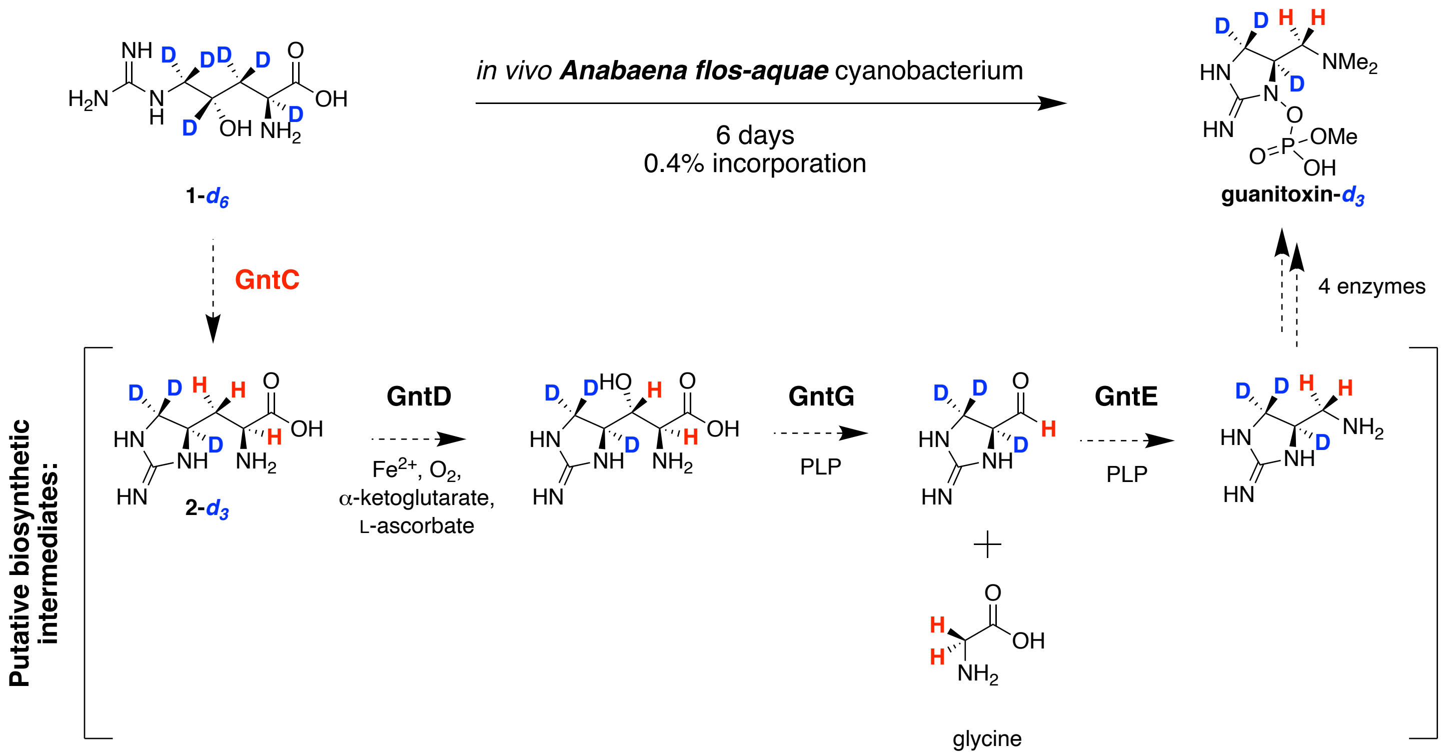

Figure S1: Previously reported *in vivo* guanitoxin stable isotope labeling results.[11] Heavy isotope labeled **1-*d*₆** was successfully incorporated into guanitoxin when fed to toxin-producing *Anabaena flos-aquae* cyanobacterium. Only three of the original deuterium atoms were retained during this biotransformation as depicted in the guanitoxin-***d*₃** structure above. Based on our previous biochemical characterization of GntD, GntE, and GntG[1] and GntC mechanistic results from this study, we propose the following biosynthetic intermediates (beginning with **2-*d*₃**), which would rationalize the *in vivo* re-introduction of the two hydrogen atoms adjacent to the dimethylamine moiety within isolated guanitoxin-***d*₃**.

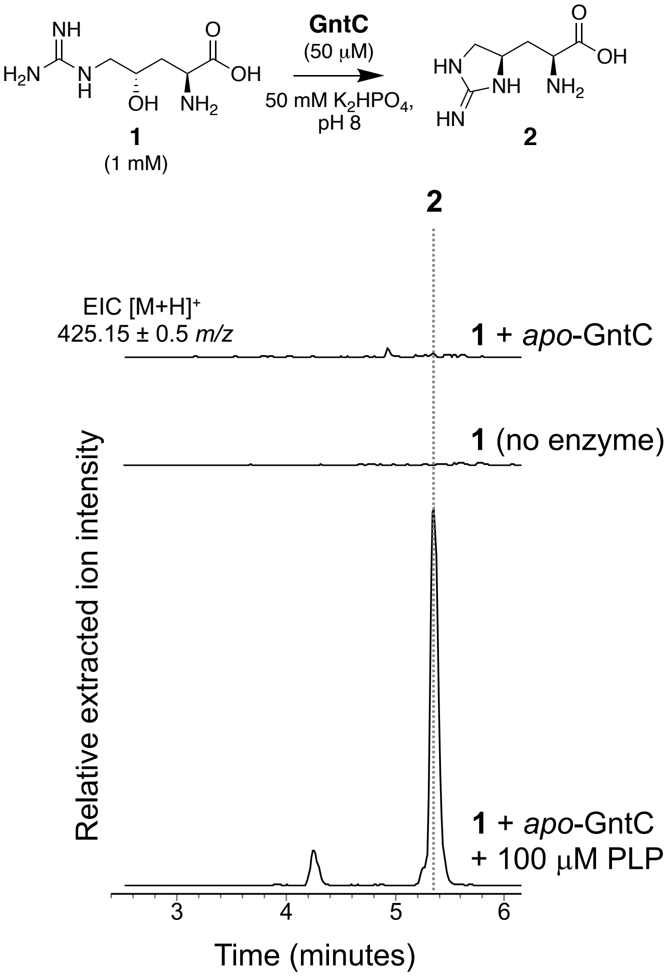

Figure S2 *In vitro* GntC PLP dependency assay. To remove the co-purifying PLP cofactor and generate the *apo*-enzyme, GntC was treated with hydroxylamine following the procedures above. Enzyme assays with *apo*-GntC (top trace), no enzyme (middle trace), and *apo*-GntC with 100 μM PLP exogenously added (bottom trace) were incubated at room temperature for 16 hours prior to Marfey's derivatization and UPLC-MS analysis. Relative intensities of positive mode extracted ion chromatograms from UPLC-MS traces were extracted for derivatized product **2** ($[M+H]^+$ 425.15 ± 0.50 *m/z*).

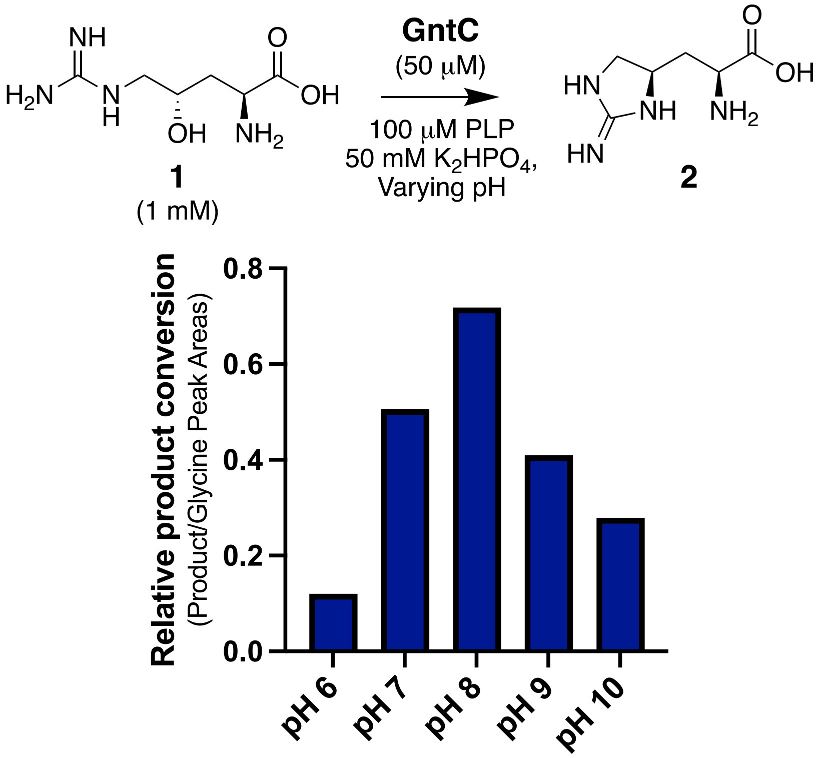

Figure S3: *In vitro* GntC pH preference assay. *In vitro* GntC assays were set up as previously described for 14 hours under buffered potassium phosphate at pHs 6 - 10. An internal standard of 500 μM glycine was added prior to Marfey's derivatization to correct for any discrepancies during derivatization and for relative product conversion measurements via LCMS analysis. Relative intensities of positive mode extracted ion chromatograms from UPLC-MS traces were extracted for derivatized **2** and glycine ($[M+H]^+$ 425.15, 328.08 ± 0.50 *m/z* respectively). The **2**:glycine peak areas ratios at the 5 pHs tested were compared and identified pH 8 as optimal for **2** production.

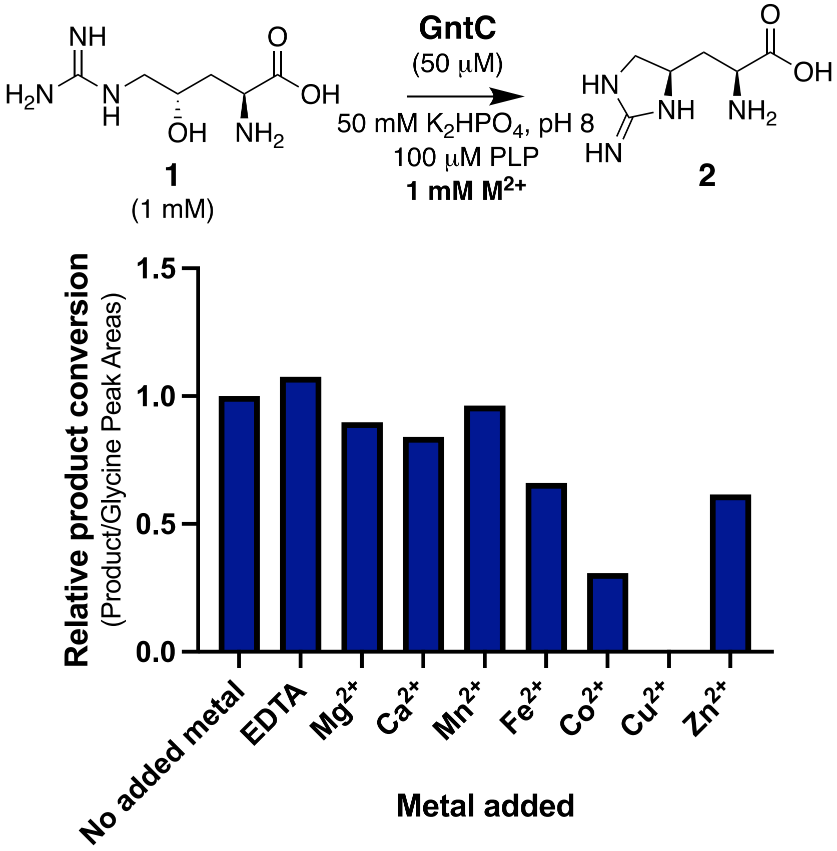

Figure S4: *In vitro* GntC divalent metal cation preference assay. *In vitro* GntC assays were set up as previously described for 16 hours with the addition of 1 mM EDTA or divalent metal salt ($MgCl_2 \cdot 6H_2O$, $CaCl_2$, $MnSO_4 \cdot H_2O$, $FeSO_4 \cdot 7H_2O$, $CoBr_2$, $CuSO_4 \cdot 5H_2O$, $ZnCl_2$) following adaption from literature resources.[12] An internal standard of 1 mM glycine was added prior to Marfey's derivatization to correct for any discrepancies during derivatization and for relative product conversion measurements via LCMS analysis. Relative intensities of positive mode extracted ion chromatograms from UPLC-MS traces were extracted for derivatized **2** and glycine ($[M+H]^+$ 425.15, 328.08 ± 0.50 *m/z* respectively). The **2**:glycine peak area ratios were compared to no added metal and showed negligible enhancement of **2** production by GntC.

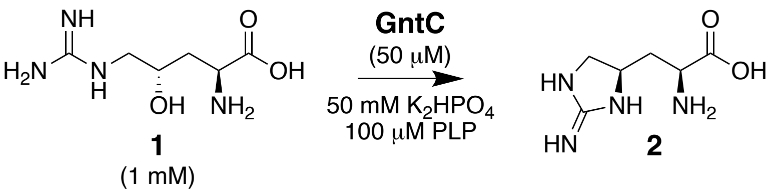

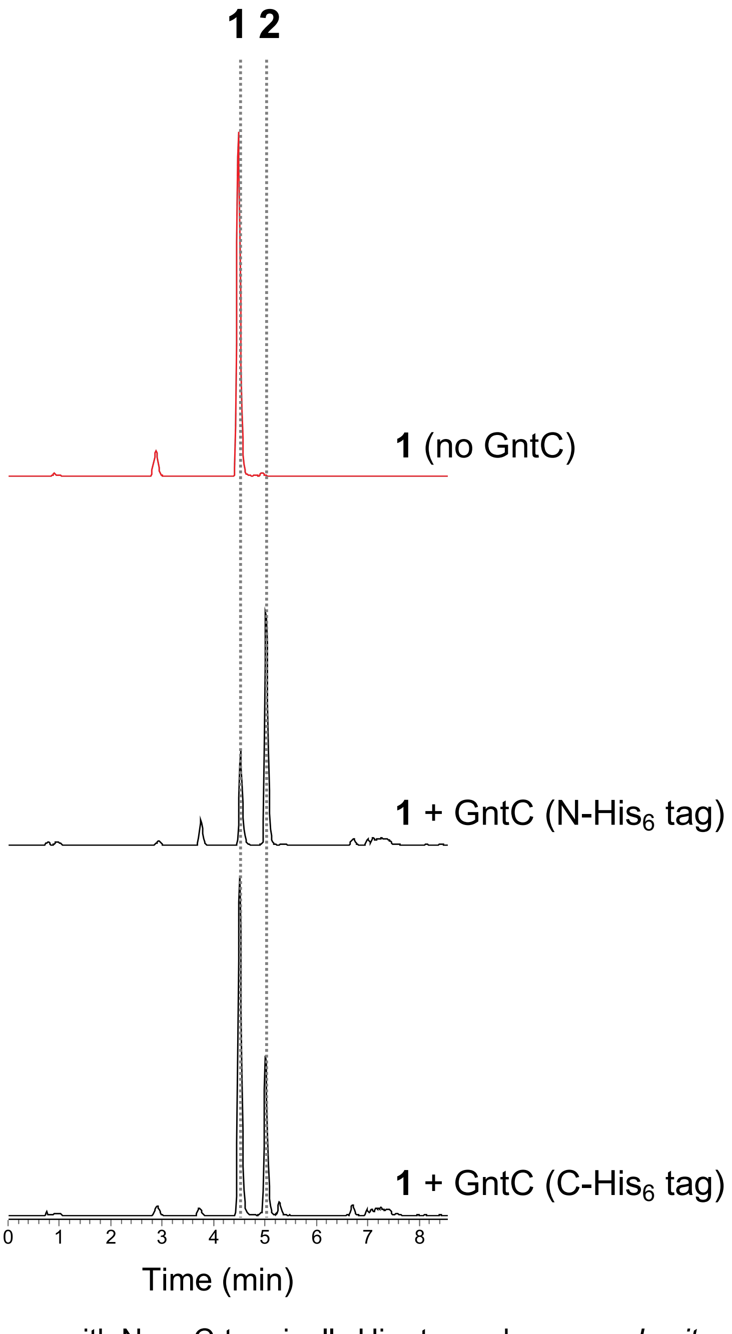

Figure S5: *In vitro* GntC assays with N- or C-terminally His₆-tagged enzyme. *In vitro* enzyme assays were set up as previously described over 16 hours without enzyme (top trace), N-terminally $His_6$-tagged GntC (middle trace), and C-terminally tagged GntC (bottom trace). Relative intensities of positive mode extracted ion chromatograms from UPLC-MS traces were extracted for derivatized **1** and **2** ($[M+H]^+$ 443.16, 425.15 ± 0.50 *m/z* respectively). Both versions of the enzyme successfully produced **2** *in vitro*, showing the negligible impact of the $His_6$-tag on GntC catalysis.

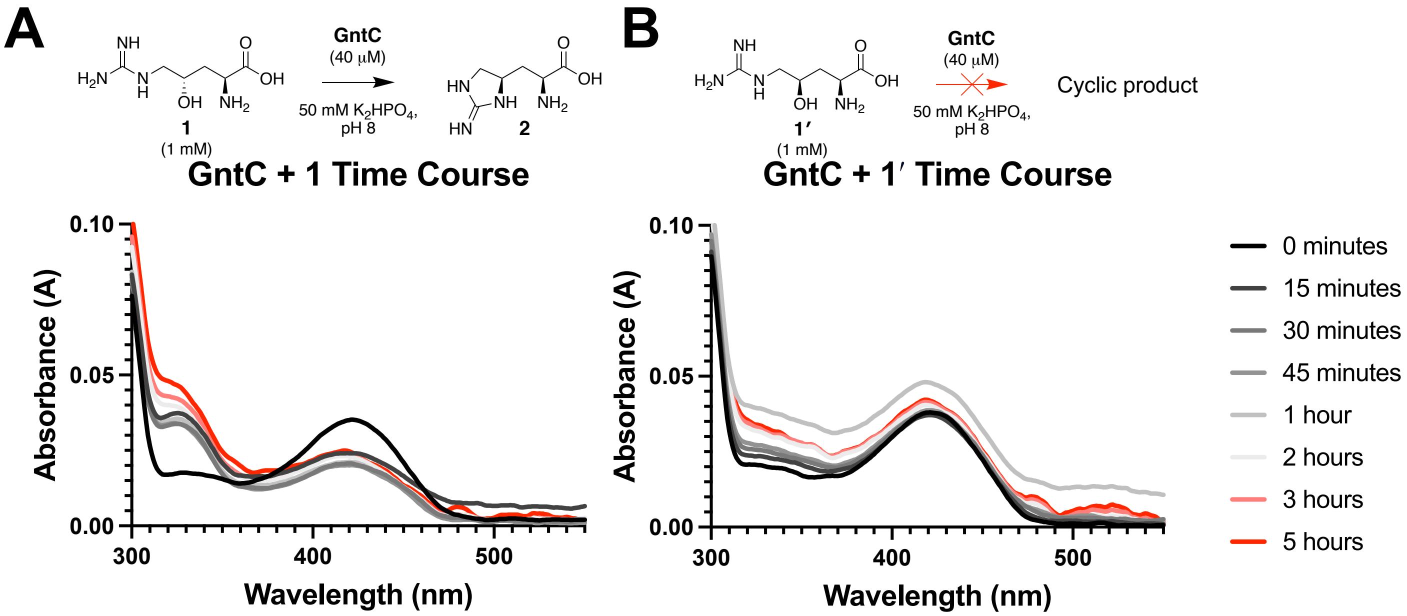

Figure S6. UV-Vis spectra of GntC assays with substrate **1** and diastereomer **1′**. *In vitro* enzyme assays were incubated at room temperature for up to 5 hours with 1 mM substrate, 40 µM GntC and 50 mM $K_2HPO_4$ buffer (pH 8.0) and no additional PLP. UV-Vis absorption spectra were taken prior to the addition of substrate **1** (left) or **1′** (right) and then at the time points listed.

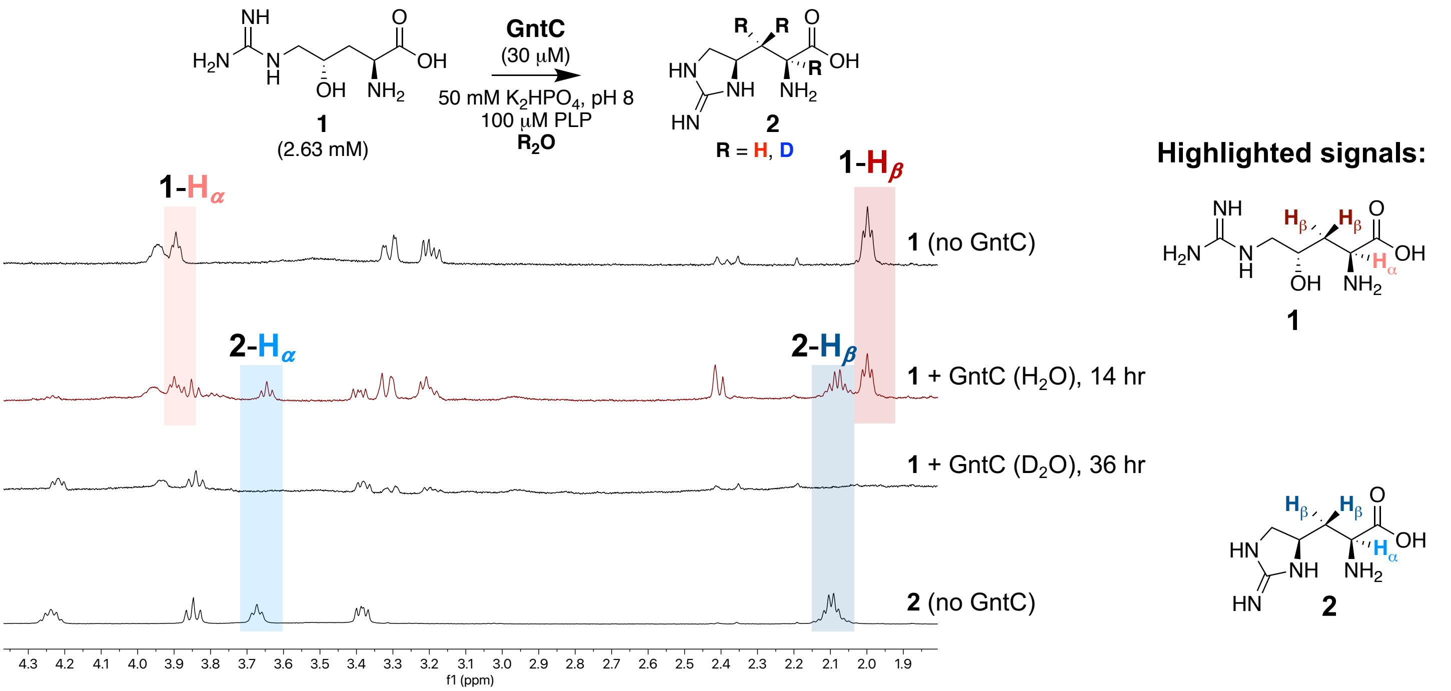

Figure S7: *In vitro* [1]H NMR GntC assay with substrate **1** in $H_2O$ and $D_2O$. *In vitro* enzyme assays were set up in a ~100:1 substrate **1**:GntC molar ratio and analyzed using 500 MHz [1]H NMR. In comparison to no enzyme controls with substrate **1** (top trace) and product **2** (bottom trace), characteristic signals for **1** and **2** are present after prolonged incubation with GntC in $H_2O$ buffer (second trace). As depicted by the red and blue boxes, α- and β- hydrogen signals for both **1** and **2** are substantially diminished under buffered $D_2O$ conditions (third trace).

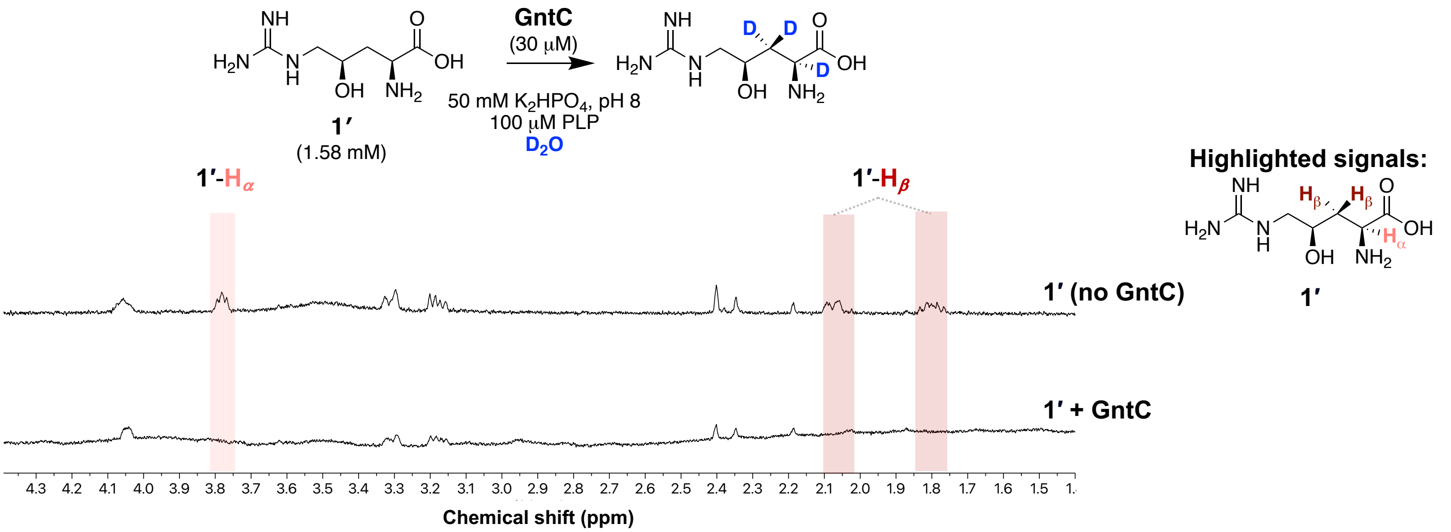

Figure S8: *In vitro* $^1$H NMR GntC assay with substrate diastereomer **1′** in $D_2O$. *In vitro* enzyme assays were set up in a ~50:1 substrate diastereomer **1′**:GntC molar ratio in buffered $D_2O$ and analyzed using 500 MHz $^1$H NMR. Following a 20 hour incubation of substrate diastereomer **1′** without (top trace) and with GntC (bottom trace), a substantial decrease in α- and β-hydrogen signals for **1′** were observed without any obvious cyclic or alternative products.

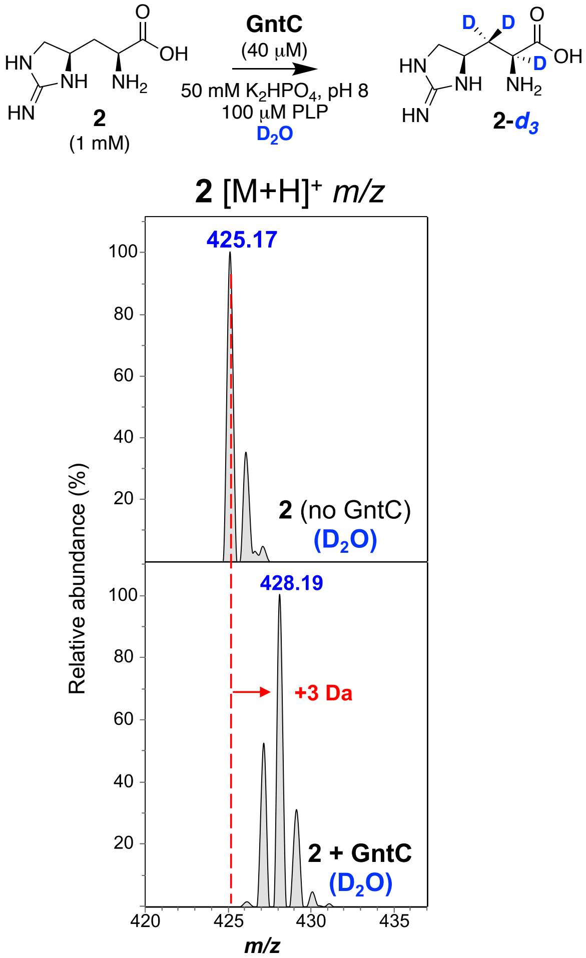

Figure S9: *In vitro* GntC assay with product L-enduracididine (**2**) in $D_2O$. *In vitro* incubation of **2** for 16 hours without (top) and with GntC (bottom) in the presence of in $D_2O$ conditions. Following Marfey's derivatization and UPLC-MS analysis, up to three 3 deuterium incorporations occur in the product (**2-*d*₃**) based on the mass spectrometry envelope.

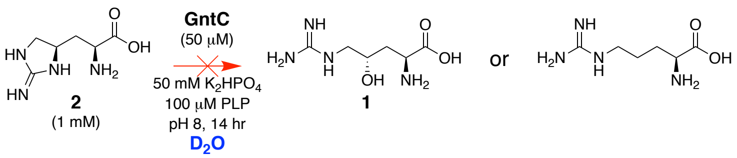

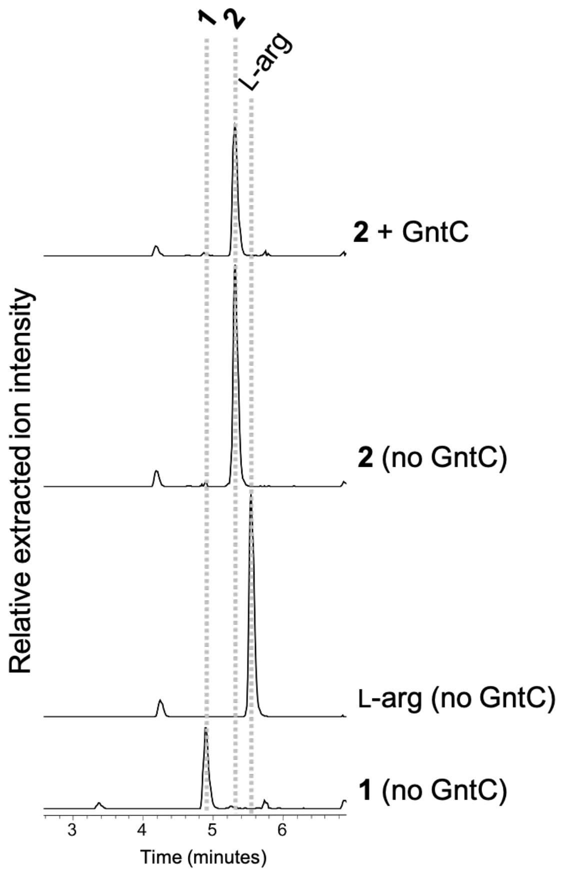

Figure S10: *In vitro* GntC assay to assess overall reaction reversibility. *In vitro* enzyme assays were set up with product **2** and incubated for 14 hours under buffered $K_2HPO_4$ in $D_2O$. Relative intensities of positive mode extracted ion chromatograms from UPLC-MS traces were extracted for all reasonably deuterated isotopologs of derivatized **2**, **1**, and L-arginine (425.15, 426.15, 427.16, 428.16, 443.16, 444.16, 445.16 ± 0.50 *m/z*). No obvious differences were observed in the presence (top trace) or absence (second trace) of GntC incubation, indicating that the overall GntC reaction is not reversible in *in vitro* conditions.

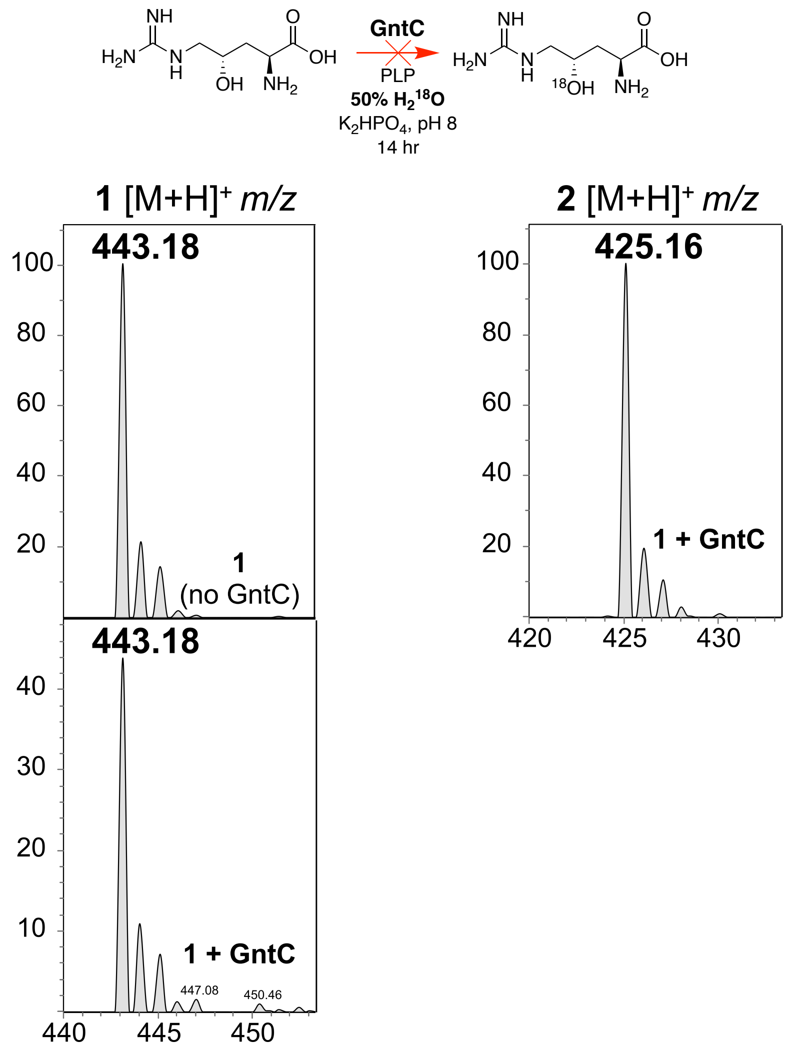

Figure S11: *In vitro* GntC assay in 50% $H_2{}^{18}O$ to assess dehydration reversibility. *In vitro* enzyme assays were set up as previously described in 50% $H_2{}^{18}O$ buffered conditions. Following Marfey's derivatization and UPLC-MS analysis, no difference in isotopic mass distribution was observed for starting material **1** when comparing the no enzyme (top left) and with GntC (bottom left) experiments. This indicates that the dehydration step is not reversible under *in vitro* GntC assay conditions.

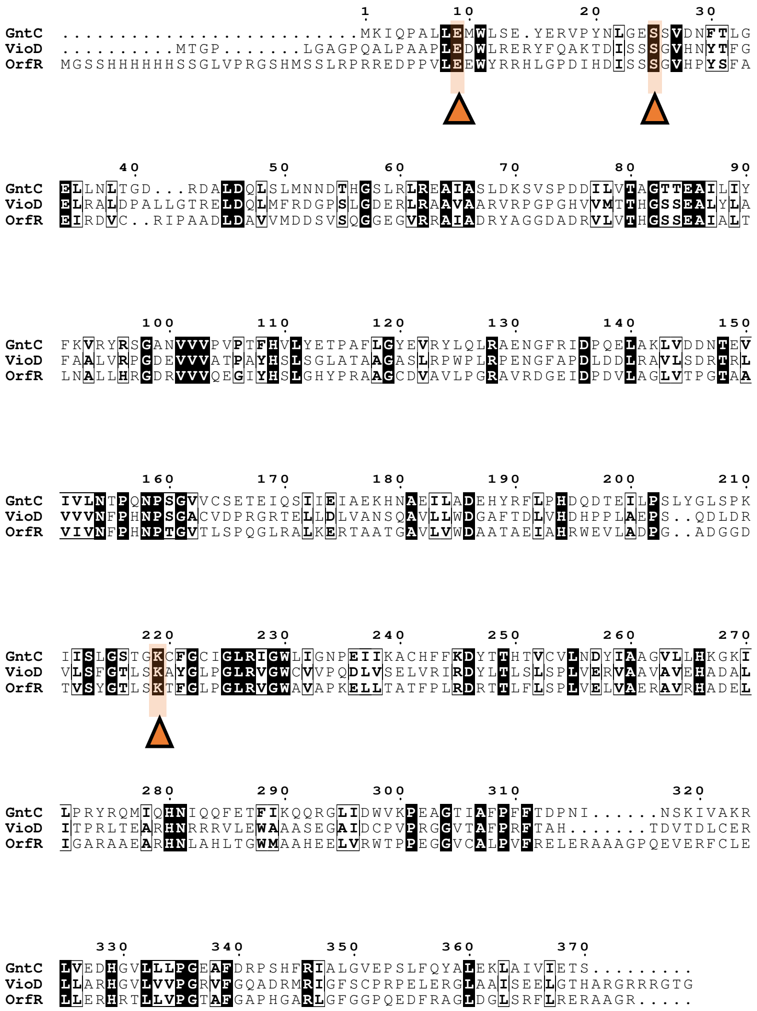

Figure S12: Sequence alignment of GntC to PLP-dependent capreomycidine cyclases. Enzymes listed are GntC (PDB ID: 8FFT), VioD, and OrfR (PDB ID: 4M2M, Chain A). Alignment was generated using Clustal Omega and viewed using ESPript 3.0. Sequence numbering follows the GntC sequence, and highlighted residues are orange with a triangle underneath them. Conserved residues include a glutamate residue towards the N-terminus of the sequences (GntC E9), a serine residue conserved amongst GntC, OrfR, and VioD (GntC S25), and a lysine residue (GntC K219). NCBI accession numbers for all sequences are listed in Table S1.

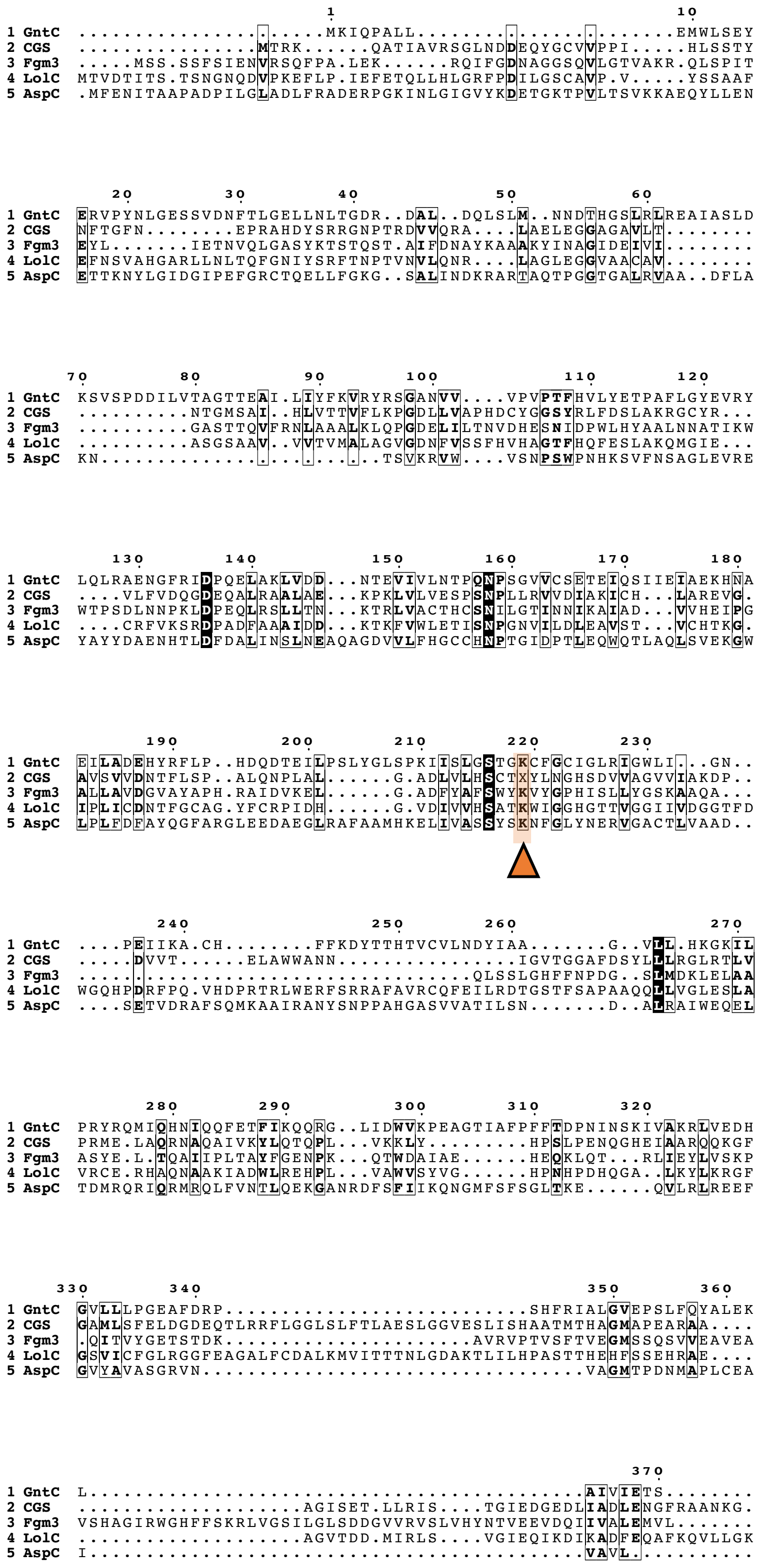

Figure S13: Sequence alignment of GntC to PLP-dependent intermolecular γ-substitution enzymes and aspartate aminotransferase AspC. The sequences were aligned in Clustal Omega and visualized in ESPript 3.0. Residues of interest are highlighted in orange, and the sequence numbering follows GntC. GntC shares minimal sequence homology to other γ-substitution enzymes, such as PLP-specific residues D186 which allows for deprotonation of the nitrogen in the pyridoxal ring of PLP, and K219, which keeps PLP in the enzyme active site. NCBI accession numbers for all sequences are listed in Table S1.

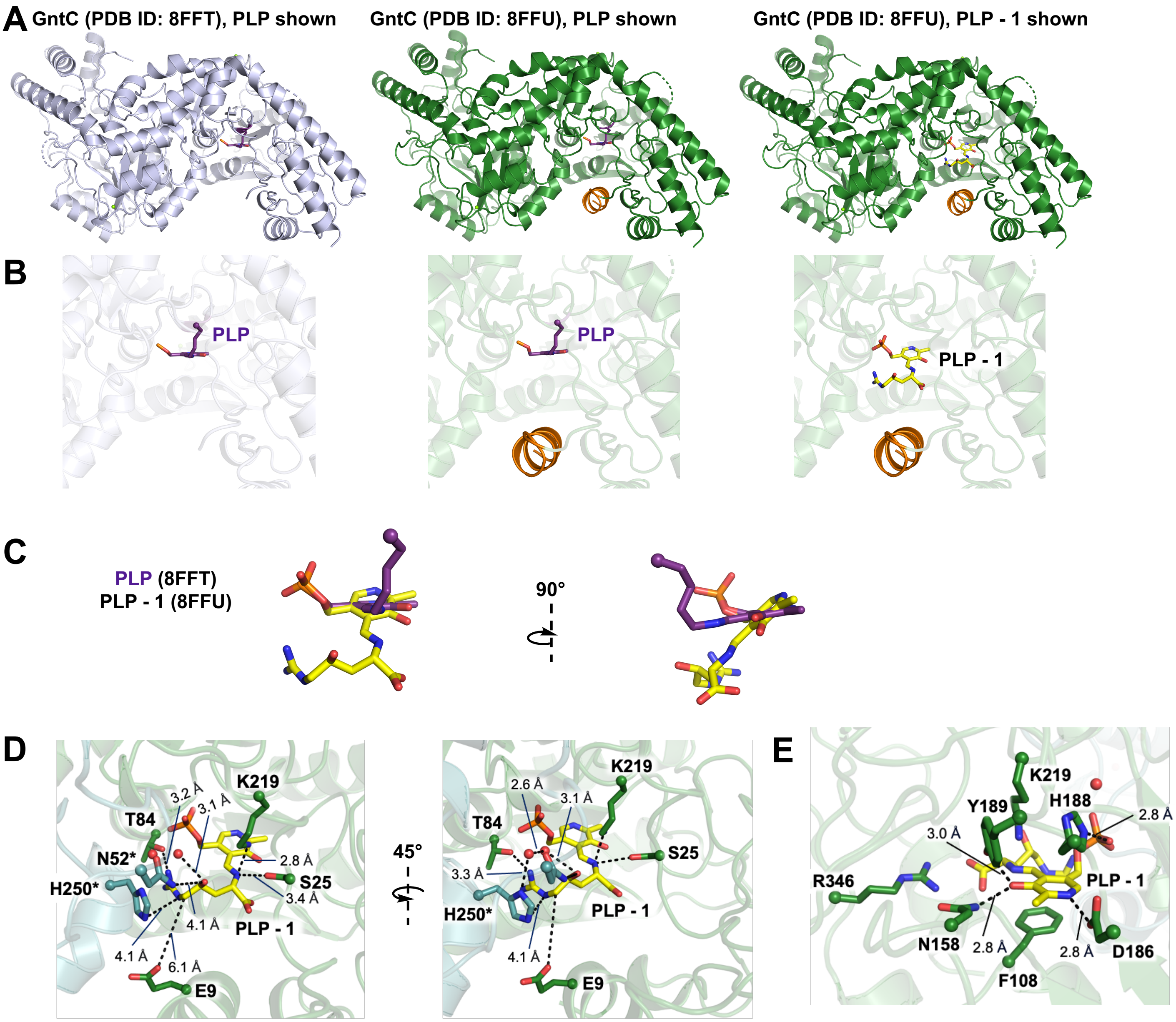

Fig. S14. Structural comparison of GntC with PLP and PLP-**1** substrates. (A) Structural comparison of GntC with bound PLP (gray, PDB ID: 8FFT) and with bound PLP-**1** ligand (green, PDB ID: 8FFU). Full view of GntC crystal structure dimers reveal minimal differences in quaternary structure except for an additional ordered α-helix (orange) at the N-terminus of GntC with bound PLP-**1**. GntC (8FFU) crystallized with two states of PLP: the internal (middle column, colored purple) and external aldimine (right column, colored yellow) intermediates. (B) Zoomed in view of GntC active site reveals PLP remains in similar positions in GntC and the additional α-helix in GntC (8FFU) helps shape the active site. (C) Overlay of the PLP internal aldimine with PLP-**1** (the external aldimine) reveals a 30° change in orientation between the two species, with the ncAA component of **1** positioned closer to this ordered α-helical region. (D) Detailed view of GntC active site with hydrogen bonding distances between residues proposed to interact with PLP-**1**. Residues from the other monomer (colored teal) are denoted with an asterisk. (E) Additional 180° rotation of the GntC active site from depiction D to highlight residues that interact with the PLP cofactor, including F108, N158, D186, H188, Y189 and R346.

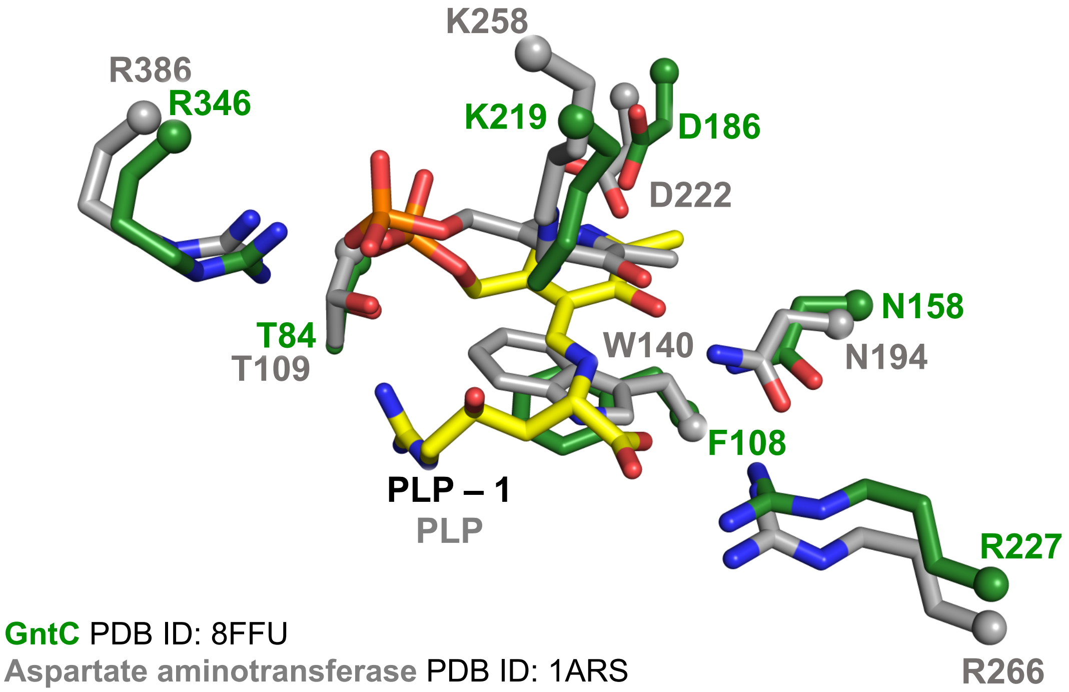

Figure S15: Active site comparison of PLP-**1** GntC to aspartate aminotransferase. Structure alignment of GntC (green) to aspartate aminotransferase (gray, PDB ID: 1ARS). Conserved residues between GntC and an aminotransferase include the canonical lysine residue needed for the internal aldimine (K219 in GntC). Some differences include a tryptophan residue that resides below the PLP in aspartate aminotransferase (W140) versus phenylalanine (GntC F108).

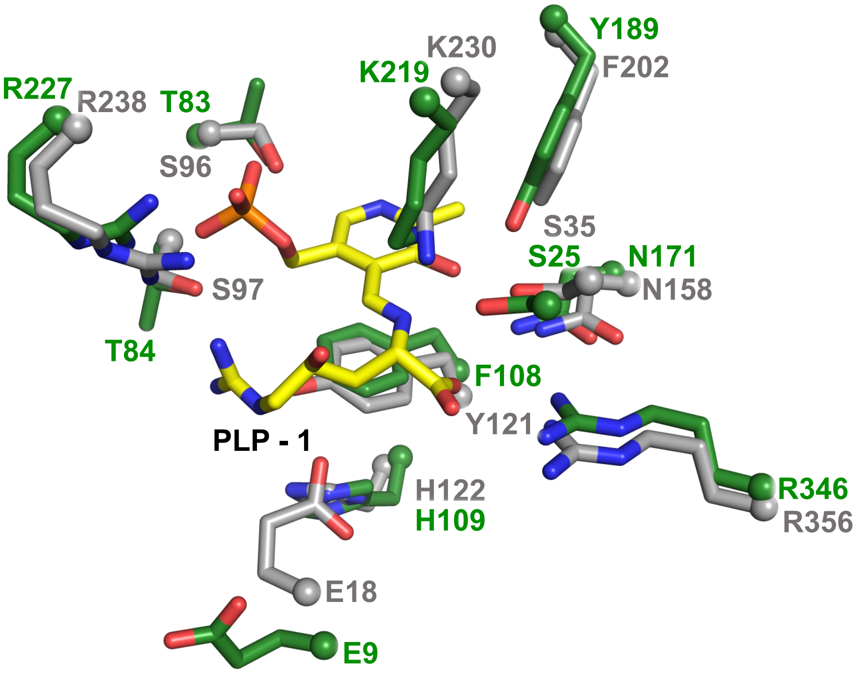

**GntC** PDB ID: 8FFU
**VioD**: AlphaFold 2.0 model

Figure S16: Active site comparison of PLP-**1** GntC and VioD AlphaFold model. Structure alignments of GntC (green, PDB ID: 8FFU) and modeled VioD (gray, structure generated with AlphaFold 2.0). Structural alignments indicate a lysine residue is conserved (GntC K219) for external aldimine formation of the two PLP-dependent enzymes. Both structures also conserve two arginine residues (GntC R346 and GntC R227), and a histidine residue (GntC H108) that sits farther away from the pyridoxal ring of PLP. Intriguingly, GntC utilizes F108 to $\pi$ stack with the PLP cofactor, whereas this residue is replaced with a tyrosine residue (Y121) in VioD. A glutamate residue (GntC E9) is conserved between both structures, which is also conserved in PLP-dependent OrfR and revealed in the sequence alignments (Figure S12). An aromatic residue is also present which is a tyrosine residue (Y189) in GntC, and a phenylalanine residue (F202) in VioD. GntC Y189 is 3 Å from the hydroxyl group of PLP in the PLP-1 adduct and is 2.3 Å from the same hydroxyl group when PLP is present as an internal aldimine.

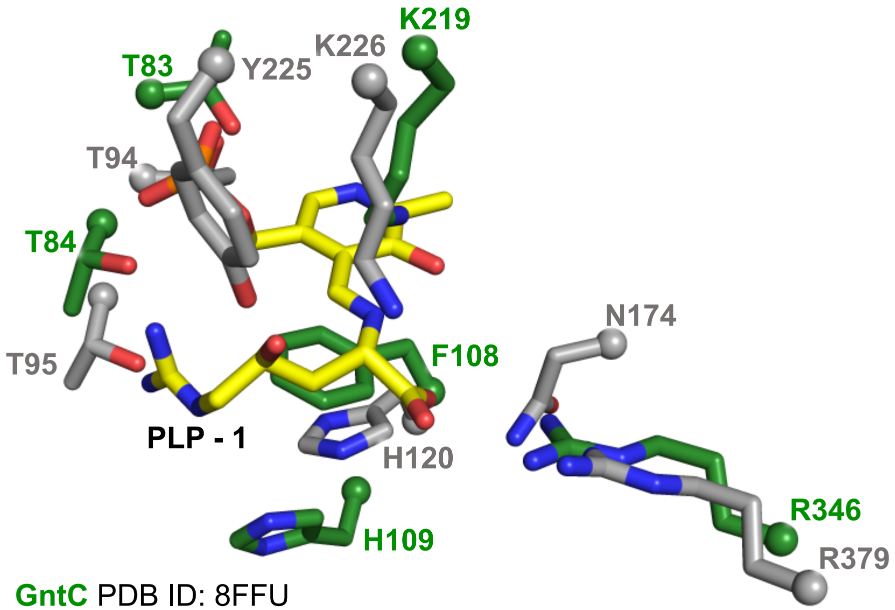

**GntC** PDB ID: 8FFU
**Fgm3**: AlphaFold 2.0 model

Figure S17: Active site comparison of PLP-**1** GntC and Fgm3 AlphaFold model. Structure alignments of GntC (green, PDB ID: 8FFU) and Fgm3 (gray, generated using AlphaFold 2.0).[14] Structural alignments indicate a lysine residue (GntC K219) for internal aldimine formation, threonine residue (GntC T84) for coordinating to the substrate guanidine group, and histidine residue (GntC H109) are conserved. Although both enzymes utilize **1** as a substrate, the Fgm3 structure contains a tyrosine residue (Y225) that could coordinate with the γ-hydroxy group of the substrate that GntC does not contain to putatively participate in retro-aldolase activity.

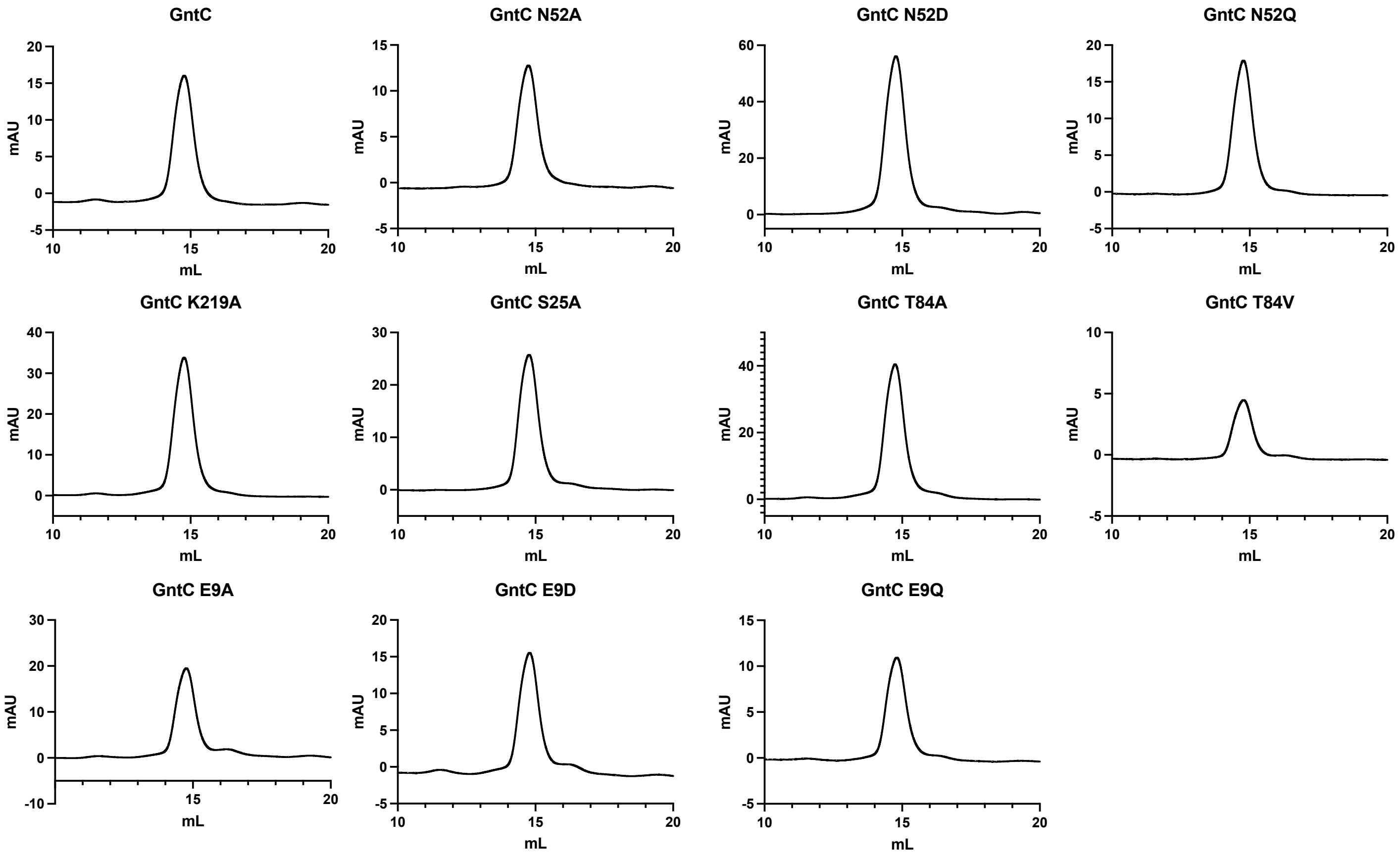

Figure S18. Analytical size exclusion chromatograms of wild-type GntC and mutants. GntC and variants were expressed and purified using the protocol detailed in the Methods and Materials section. Samples were diluted to 4 mg/mL before injection onto an analytical size exclusion column. No obvious retention time difference was observed between GntC and its site-directed mutants, suggesting similar homodimeric forms.

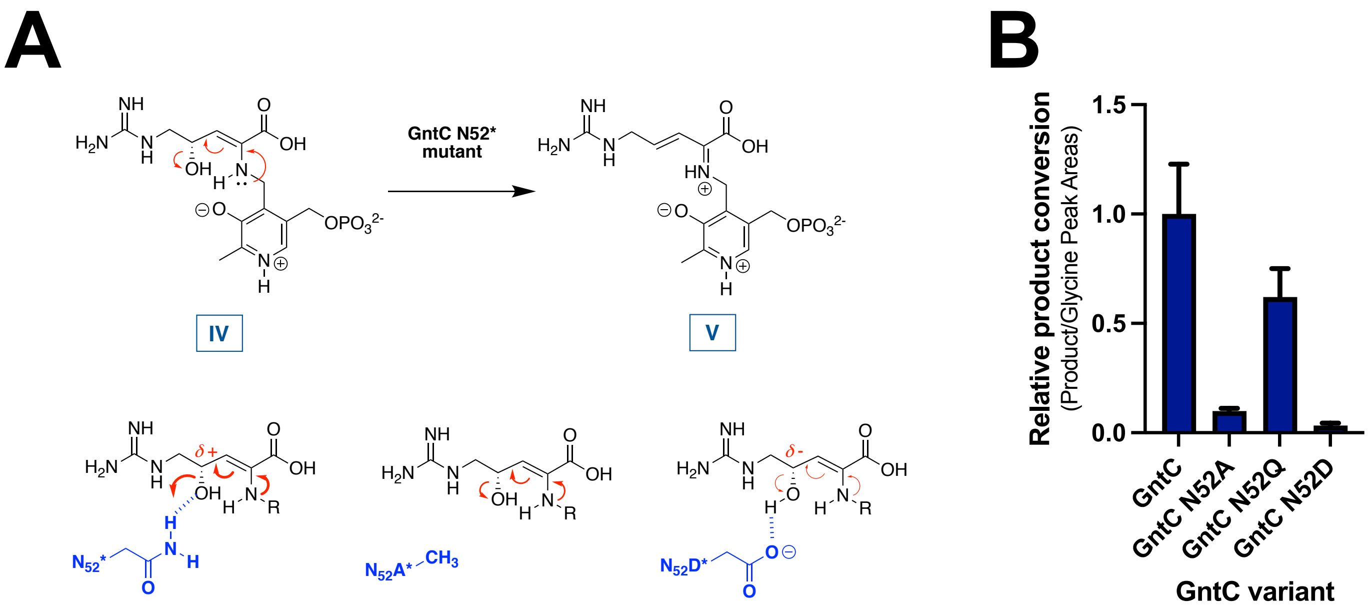

Figure S19. Proposed rationale for how GntC N52* enhances γ-hydroxy elimination. (A) Putative role of N52 on directly (or through a coordinated water molecule not depicted) facilitating the elimination of the γ-hydroxy group through a hydrogen bond interaction. The absence of a hydrogen bond (N52A) or presence of an anionic residue (N52D) would theoretically decrease the dehydration step. (B) Product conversion of GntC N52* mutants compared to GntC were assessed via LCMS analysis with a 500 μM internal glycine standard and performed in triplicate. The N52Q rescue of activity supports the significance of an amide side chain at this position for efficient production of **2**.

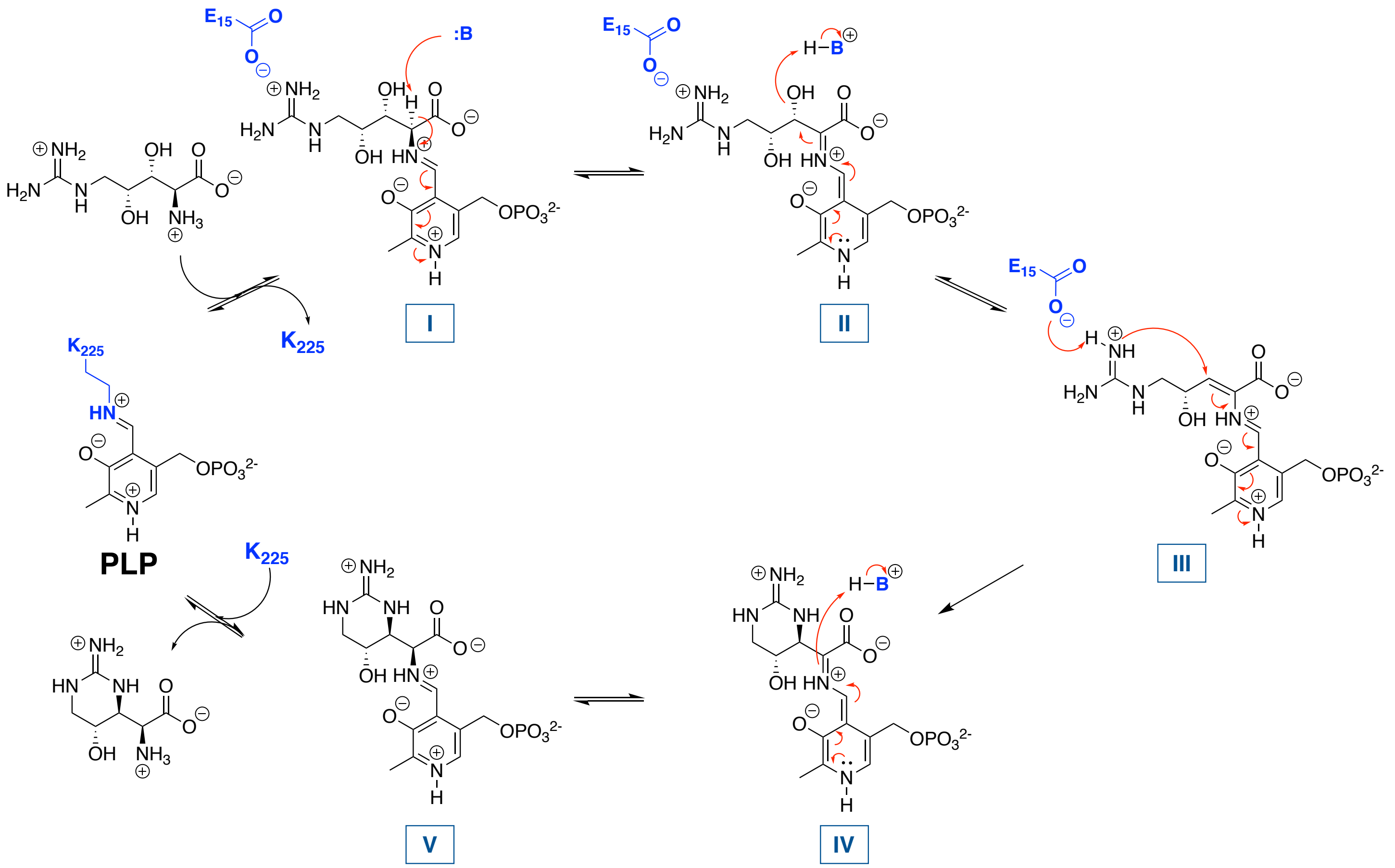

Figure S20. PLP-dependent OrfR capreomycidine cyclase is mechanistically distinct from GntC.[15] A similar external aldimine and α-hydrogen deprotonation occur (intermediate I), however the quinonoid intermediate II is proposed to eliminate the β-hydroxy group. This generates extensively conjugated intermediate III and restores aromaticity within the PLP-cofactor. A diastereoselective nucleophilic addition of the E15-deprotonated guanidine moiety at the β-position is controlled by OrfR to generate the capreomycidine six-membered ring (intermediate IV), which is then reprotonated at the α-position and external aldimine V has its ncAA moiety released from the enzyme through the action of active site K225.

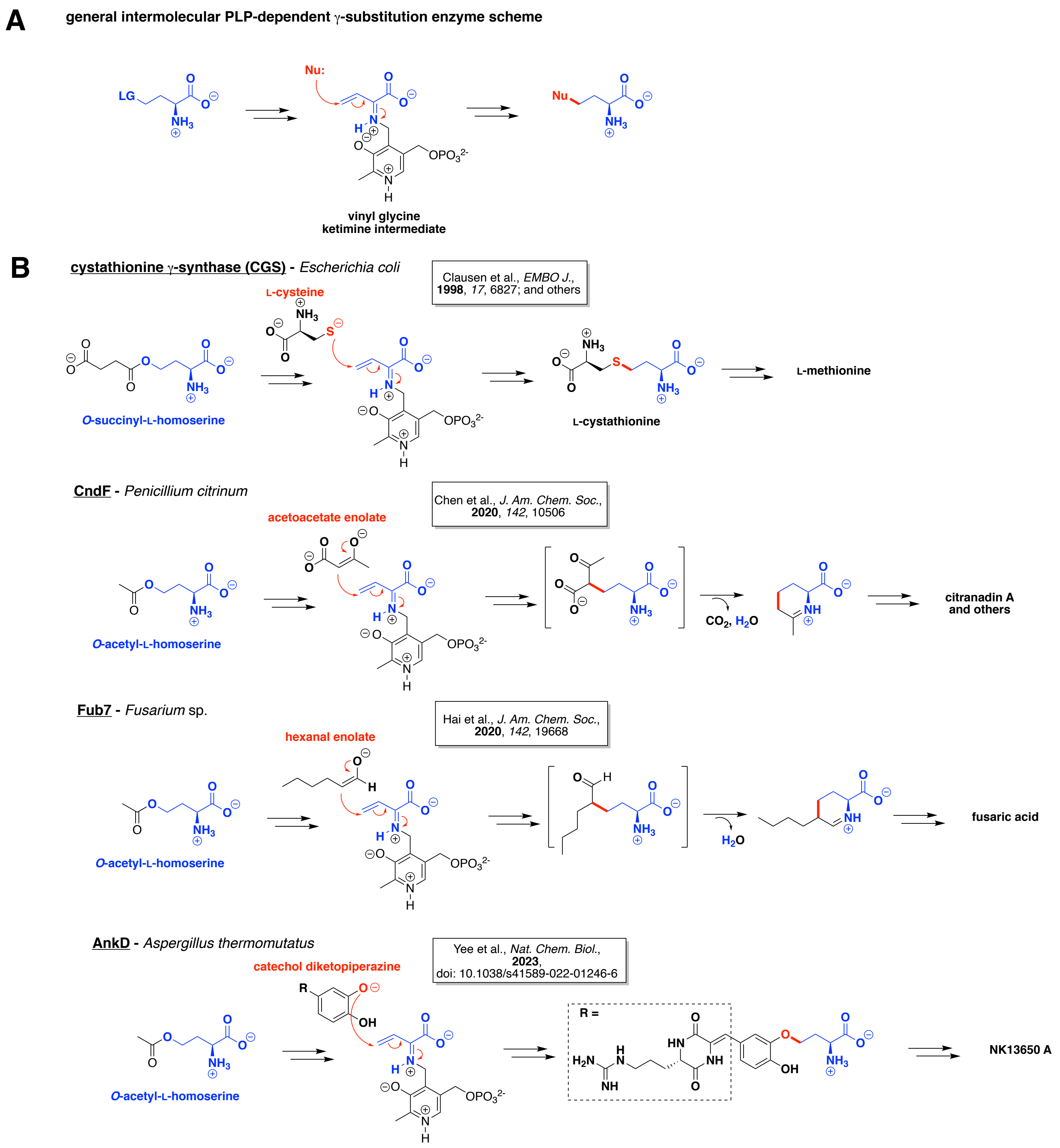

A
general intermolecular PLP-dependent γ-substitution enzyme scheme
LG
Nu:
Nu
vinyl glycine
ketimine intermediate
B
cystathionine γ-synthase (CGS) - Escherichia coli
Clausen et al., EMBO J., 1998, 17, 6827; and others
L-cysteine
O-succinyl-L-homoserine
L-cystathionine
L-methionine
CndF - Penicillium citrinum
Chen et al., J. Am. Chem. Soc., 2020, 142, 10506
acetoacetate enolate
O-acetyl-L-homoserine
CO2, H2O
citranadin A
and others
Fub7 - Fusarium sp.
Hai et al., J. Am. Chem. Soc., 2020, 142, 19668
hexanal enolate
O-acetyl-L-homoserine
H2O
fusaric acid
AnkD - Aspergillus thermomutatus
Yee et al., Nat. Chem. Biol., 2023, doi: 10.1038/s41589-022-01246-6
catechol diketopiperazine
O-acetyl-L-homoserine
R =
NK13650 A

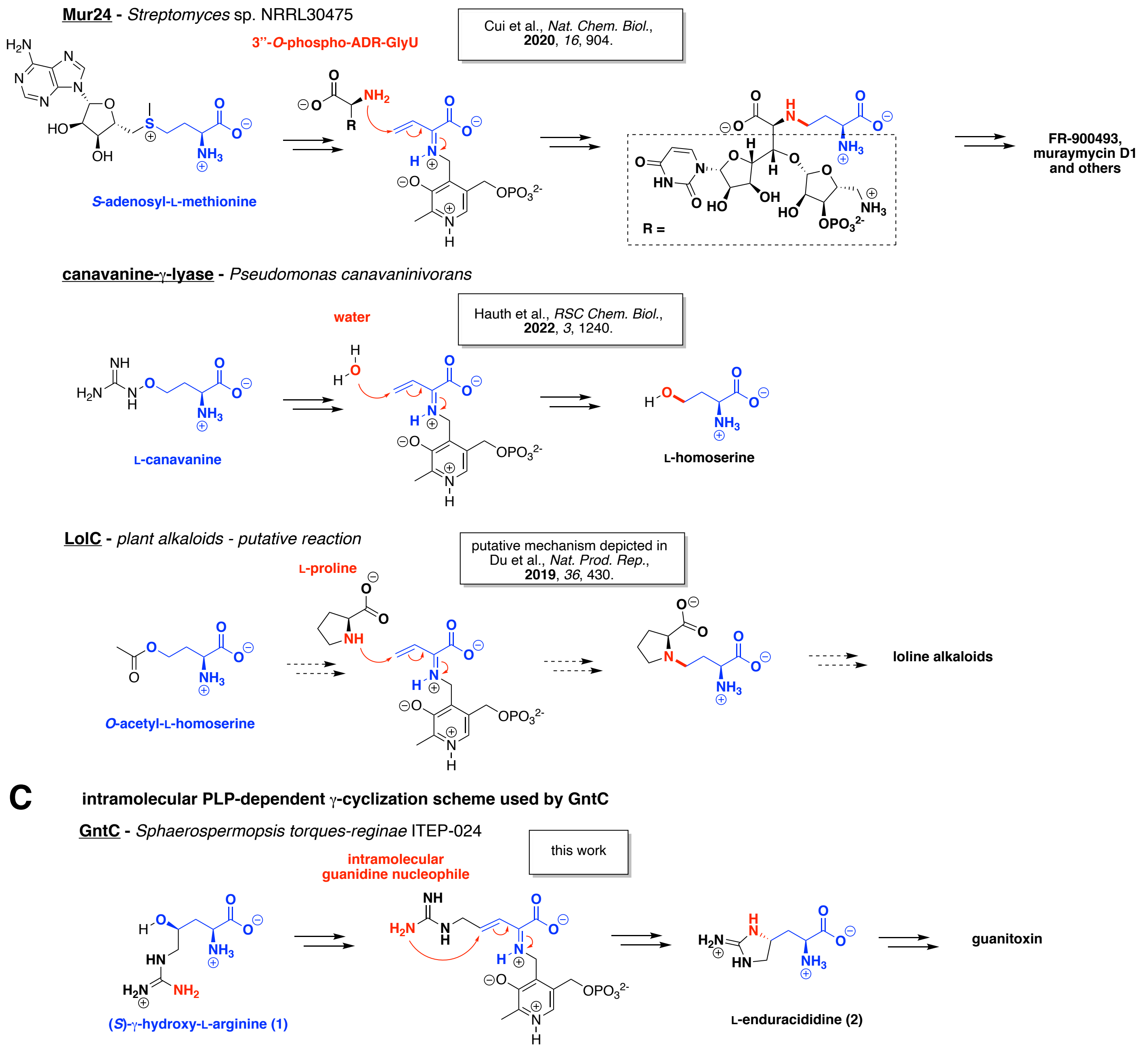

Figure S21. PLP-dependent intermolecular γ-substitution reactions from diverse biosynthetic pathways are functionally similar to the intramolecular GntC reaction. A) General PLP-dependent intermolecular γ-substitution reaction scheme. An activated γ-functionalized amino acid (blue) is enzymatically converted to a convergent vinyl glycine ketimine intermediate. An intermolecular nucleophile (red) adds to the γ-position via a Michael addition, eventually generating a γ-substituted ncAA product. B) Literature examples of intermolecular γ-substitution enzymes from diverse biosynthetic pathways that have been functionally characterized or putatively would occur through this mechanism.[12,16–21] C) Comparison of the intramolecular γ-substitution strategy adapted by GntC in the production of **2**.

Table S1: NCBI accession codes for protein sequences used in this study.

| | Organism | NCBI Accession Number |
|---|---|---|
| GntC | *Sphaerospermopsis torques-reginae ITEP-024* | QYX30483.1 |
| VioD | *Streptomyces vinaceus* | AAO66428.1 |
| OrfR | *Streptomyces lavendulae* | WP_030227708.1 |
| CGS | *Escherichia coli (strain K12)* | WP_001417137.1 |
| Fgm3 | *Fusarium graminearum PH-1* | A0A1C3YKE0.1 |
| LolC | *Neotyphodium sp. PauTG-1* | ABQ57528.1 |
| AspC | *Escherichia coli* | WP_000462687 |
| CndF | *Penicillium citrinum* DSM1997 | From literature reference[15] |
| Aminotransferase | *Streptomyces suis 89-1591* | ZP_03625122.1 |

Table S2: Data collection and refinement statistics of GntC

| | GntC | GntC with **1** bound |
|---|---|---|
| PDB ID | 8FFT | 8FFU |
| Beamline | ALS 8.2.1 | SSRL 12-2 |
| Wavelength (Å) | 0.99992 | 0.97946 |
| Space Group | $P2_1$ | $P2_1$ |
| Cell dimensions (Å) | a = 60.86, b = 158.16, c = 73.23, β = 90.01° | a = 61.06, b = 158.49, c = 73.60, β = 90.02° |
| Resolution | 73.2-2.10 (2.11-2.10) | 40.0-2.04 (2.16-2.04) |
| # unique reflections | 75093 (778) | 87584 (14110) |
| Completeness (%) | 93.1 (97.3) | 98.6 (98.8) |
| Redundancy | 5.2 (4.6) | 5.8 (5.9) |
| <I/sigI> | 10.9 (2.2) | 13.6 (1.9) |
| $R_{sym}$ | 0.115 (0.706) | 0.094 (0.786) |
| $CC_{1/2}$ | (0.57) | (0.71) |
| Resolution | 73.23-2.10 | 36.80-2.04 |
| # unique reflections | 75,048 | 87,583 |
| $R_{work}$ (%) / $R_{free}$ (%) | 22.0/24.2 | 18.0/21.1 |
| RMS bond length (Å) | 0.020 | 0.004 |
| RMS bond angles (Å) | 0.744 | 0.727 |
| # of atoms | | |
| Protein Atoms | 10975 | 11231 |
| Lysine-PLP | 96 | 96 |
| PLP-(*S*)-4-hydroxy-arginine | - | 56 |
| Water molecules | 103 | 163 |
| $Mg^{2+}$ | 5 | 4 |
| Average B-factor (Å$^2$) | | |
| Protein Atoms | 32.8 | 31 |
| Lysine-PLP | 29.1 | 27.3 |
| PLP-(*S*)-4-hydroxy-arginine | - | 27.1 |
| Water molecules | 26.4 | 27.1 |
| $Mg^{2+}$ | 30.4 | 29.7 |
| Ramachandran plot | | |
| Favored (%) | 97.52 | 97.73 |
| Allowed (%) | 2.41 | 2.27 |
| Outliers (%) | 0.07 | 0.00 |
| Rotamer outliers (%) | 0.75 | 0.57 |

Table S3: Residues modeled in each chain of GntC (residues 1-370)

| | GntC | GntC with **1** bound |
|---|---|---|
| Chain A | 18-194, 200-369,<br>K219 is covalently bound with PLP | 6-194, 198-369, K219 is covalently bound with PLP (20% occupancy), PLP with **1** bound (80% occupancy) |
| Chain B | 17-194, 198-369,<br>K219 is covalently bound with PLP | 17-369, K219 is covalently bound with PLP (100% occupancy) |
| Chain C | 19-194, 200-369,<br>K219 is covalently bound with PLP | 6-194, 198-367, K219 is covalently bound with PLP (20% occupancy), PLP with **1** bound (80% occupancy) |
| Chain D | 18-369,<br>K219 is covalently bound with PLP | 17-369, K219 is covalently bound with PLP (100% occupancy) |

Table S4: Primers used in this study

| Name | Primer sequence |
|---|---|
| GntC-6xHis-Cterm-F | CTTTAAGAAGGAGATATACCATGAAGATCCAGCCGGCG |
| GntC-6xHis-Cterm-R | GGTGGTGGTGGTGCTCGAGGCTGGTTTCAATAACGATCGCCAG |
| GntC-N52A-F | GAGCCTGATGGCCAACGATACCCATGG |
| GntC-N52Q-F | GAGCCTGATGCAGAACGATACCCATG |
| GntC-N52D-F | GAGCCTGATGGACAACGATACCC |
| GntC-N52X-R | AGTTGGTCCAGCGCATCA |
| GntC-T84A-F | CGCGGGTACCGCCGAAGCGATCC |
| GntC-T84A-R | GTAACCAGGATATCGTCCGGGCTCACGCTCTTATCCAGG |
| GntC-T84S-F | CGCGGGTACCTCCGAAGCGATCC |
| GntC-T84S-R | GTAACCAGGATATCGTCCGGGCTCACGC |
| GntC-T84V-F | CGCGGGTACCGTCGAAGCGATCC |
| GntC-T84V-R | GTAACCAGGATATCGTCCGGGCTCACG |
| GntC-K219A-F | TAGCACCGGCGCATGCTTTGGTT |
| GntC-K219A-R | CCCAGGCTAATGATCTTC |
| GntC-S25A-F | CCTGGGCGAGGCCAGCGTTGACA |
| GntC-S25A-R | TTATACGGCACACGTTCG |
| GntC-E9A-F | GCGCTGCTGGCGATGTGGCTG |
| GntC-E9A-R | CGGCTGGATCTTCATATGGCTGCCGCG |
| GntC-E9D-F | CGCTGCTGGATATGTGGCTGA |
| GntC-E9D-R | CCGGCTGGATCTTCATATGGCTG |
| GntC-E9Q-F | GGCGCTGCTGCAGATGTGGCT |
| GntC-E9Q-R | GGCTGGATCTTCATATGGCTGCCGC |
| GntC-H250A-F | CTACACCACCGCCACCGTTTGCG |
| GntC-H250A-R | TCTTTAAAGAAGTGGCAC |

**Chemical Synthesis**

Synthesis of γ-deuterated substrate **1**-$d_\gamma$

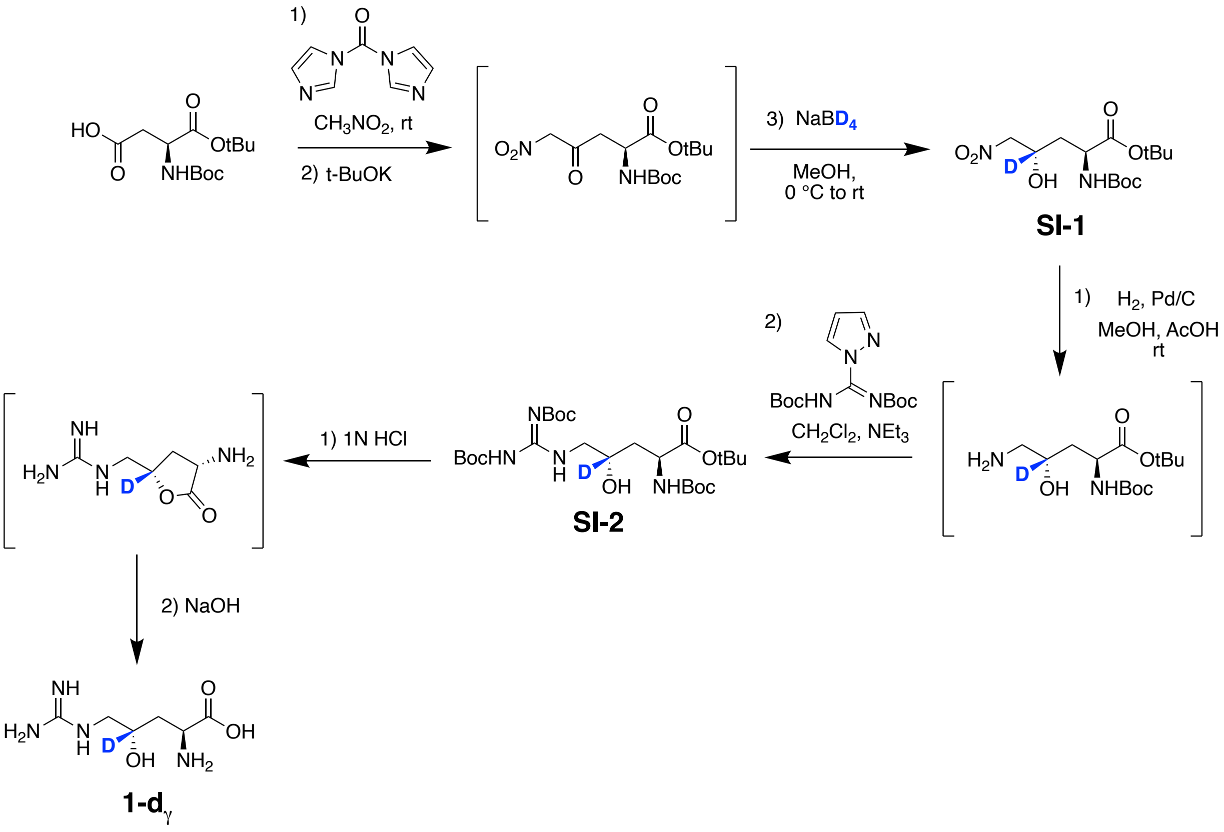

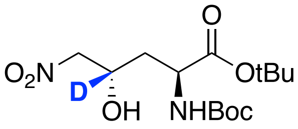

**SI-1**

The procedure to synthesize SI-1 was adapted from literature references.[1,22] Boc-L-Asp-OtBu (1.002 g, 3.46 mmol) was dried and concentrated with toluene (15 mL) *in vacuo* to remove residual water. Nitromethane (20 mL) and 1,1′-carbonyldiimidazole (0.582 g, 3.59 mmol) were added sequentially and stirred at room temperature for 45 minutes under Ar gas. Potassium tert-butoxide (0.783 g, 6.98 mmol) was added and stirred at room temperature for an additional 3 hours. The reaction mixture was quenched by the addition of 50% glacial acetic acid (50 mL), then extracted using ethyl acetate (3 x 50 mL). Pooled organic layers were washed with water (50 mL), saturated sodium bicarbonate (50 mL), water again (50 mL), then brine (50 mL). The organic layers were dried over $MgSO_4$, filtered, then concentrated *in vacuo*. The crude product was dried twice with toluene co-evaporations, then resuspended in methanol (10 mL). Sodium borodeuteride (0.1499 g, 3.58 mmol) was added portion wise at 0 °C and let reach room temperature over the course of 25 hours. The reaction was quenched with 1 N HCl until the pH was 3-4. The reaction was concentrated *in vacuo*, resuspended in water (20 mL), then extracted with ethyl acetate (3 x 25 mL). The organic layers were pooled, washed with brine (20 mL), dried over $MgSO_4$, filtered, then concentrated *in vacuo*. The crude product was purified via a silica flash column with a 4-6% diethyl ether:dichloromethane eluent system to separate the product diastereomers. A second column with a 90:5:5 toluene:THF:ethyl acetate eluent system was used to separate the diastereomer products from impurities that co-eluted from the initial column to give a white solid (0.343 g, 30%). $^1$H NMR (500 MHz, $CDCl_3$) δ 5.42 (d, *J* = 7.9 Hz, 1H), 4.70 (s, 1H), 4.49 (d, *J* = 12.4 Hz, 1H), 4.45 – 4.37 (m, 1H), 4.33 (d, *J* = 12.5 Hz, 1H), 1.95 (dd, *J* = 13.4, 3.1 Hz, 1H), 1.59 (d, *J* = 7.0 Hz, 1H), 1.53 (d, *J* = 12.5 Hz, 1H), 1.47 (d, *J* = 1.4 Hz, 9H), 1.46 (d, *J* = 1.4 Hz, 10H). $^{13}$C NMR (126 MHz, $CDCl_3$) δ 171.12, 157.32, 83.22, 81.35, 79.97, 79.89, 65.03 (*J* = 21.3 Hz), 50.71, 38.65, 28.36, 28.11; HRMS (MALDI) Calculated for $C_{14}H_{25}DN_2O_7Na$ 358.1695, found 358.1701 $(M+Na)^+$.

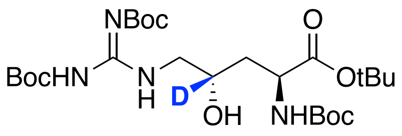

**SI-2**

This reaction was adapted from a literature reference.[1] An aliquot of SI-1 (0.327 g, 0.98 mmol) was suspended in methanol (15 mL) and acetic acid (0.056 mL, 0.976 mmol). This solution had 10% Pd/C (0.117 g, 1.10 mmol) added to it, and was flushed with Ar gas, then was left under a $H_2$ atmosphere for 16 hours. The reaction mixture was filtered through a Celite pad then concentrated *in vacuo*. The crude mixture was resuspended in toluene (20 mL) and N,N′-di-Boc-1H-pyrazole-1-carboxamidine (0.310 g, 1.00 mmol) and triethylamine (0.68 mL, 4.88 mmol) added sequentially. This solution was heated to 55 °C and stirred for 20 hours. The reaction was quenched with saturated aqueous $NH_4Cl$ (25 mL), and the organic components were extracted with ethyl acetate (3 x 25 mL), then dried over magnesium sulfate, and filtered prior to concentrating *in vacuo*. The crude reaction was purified with silica flash chromatography using an eluent of 4:1 hexanes:ethyl acetate + 0.1% triethylamine. Fractions were pooled, then concentrated *in vacuo* to yield the desired product as a white solid (0.297 g, 56%). $^1$H NMR (499 MHz, $CDCl_3$) $\delta$11.45 (s, 1H), 8.72 (s, 1H), 5.46 (d, *J* = 7.8 Hz, 1H), 4.88 (s, 1H), 4.36 (t, *J* = 9.2 Hz, 1H), 3.70 (dd, *J* = 13.9, 6.5 Hz, 1H), 3.27 (d, *J* = 13.9 Hz, 1H), 1.88 (dt, *J* = 13.6, 3.2 Hz, 1H), 1.57 (d, *J* = 12.3 Hz, 2H), 1.49 (q, *J* = 4.3, 3.8 Hz, 20H), 1.46 (s, 9H), 1.44 (s, 9H). $^{13}$C NMR (126 MHz, $CDCl_3$) $\delta$ 171.78, 163.37, 156.84, 153.06, 83.33, 82.55, 80.52, 79.51, 66.62 (*J* = 20.0 Hz), 51.30, 46.45, 38.89, 28.41, 28.19, 28.12, 27.97; HRMS (MALDI) Calculated for $C_{25}H_{45}DN_4O_9$ 548.3401, found 548.3409 $(M+H)^+$.

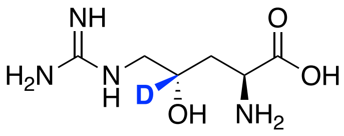

**1-d$_\gamma$**

A solution of 1 N HCl (10 mL) was added to SI-2 (0.107 g, 0.195 mmol). The reaction was stirred for 2 hours, then concentrated *in vacuo*. The crude reaction mixture was resuspended in water (10 mL) and NaOH (0.039 g, 0.98 mmol), then 1 N HCl was added dropwise to the reaction until the pH reached 7, then lyophilized overnight to give a white solid (0.112 g, quantitative). $^1$H NMR (500 MHz, $D_2O$ + 0.1% MeOH) $\delta$ 4.13 (s, 1H), 3.59 – 3.47 (m, 2H), 3.42 (d, *J* = 14.7 Hz, 1H), 2.22 (s, 2H).$^{13}$C NMR (126 MHz, $D_2O$ + 0.1% MeOH) $\delta$ 174.01, 157.42, 66.71 (*J* = 21.3 Hz), 52.62, 49.00, 46.97, 46.88, 33.29; HRMS (MALDI) Calculated for $C_6H_{13}DN_4O_3$ 192.1202, found 192.1207 $(M+H)^+$.

**NMR and Compound Characterization**

**SI-1:**

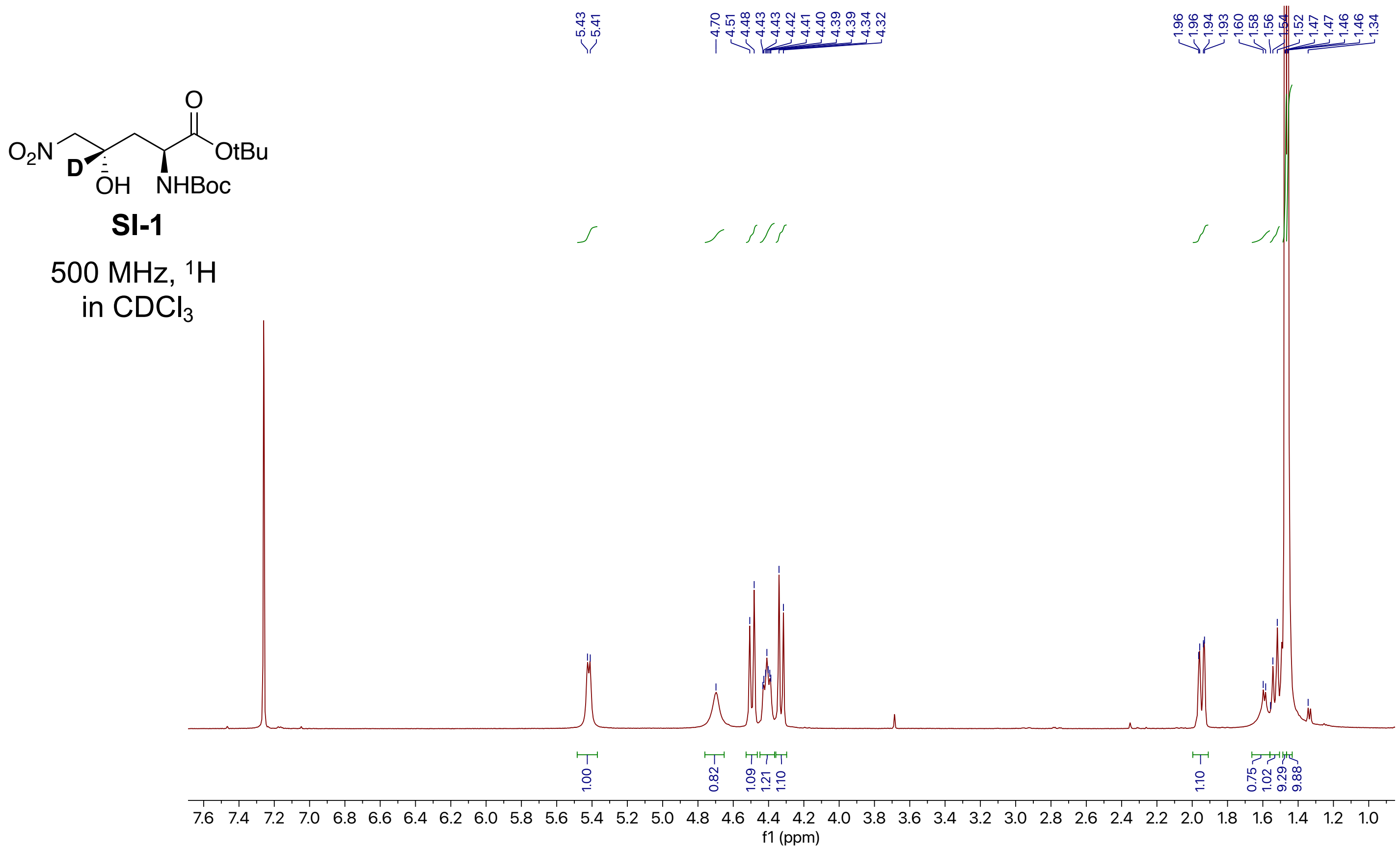

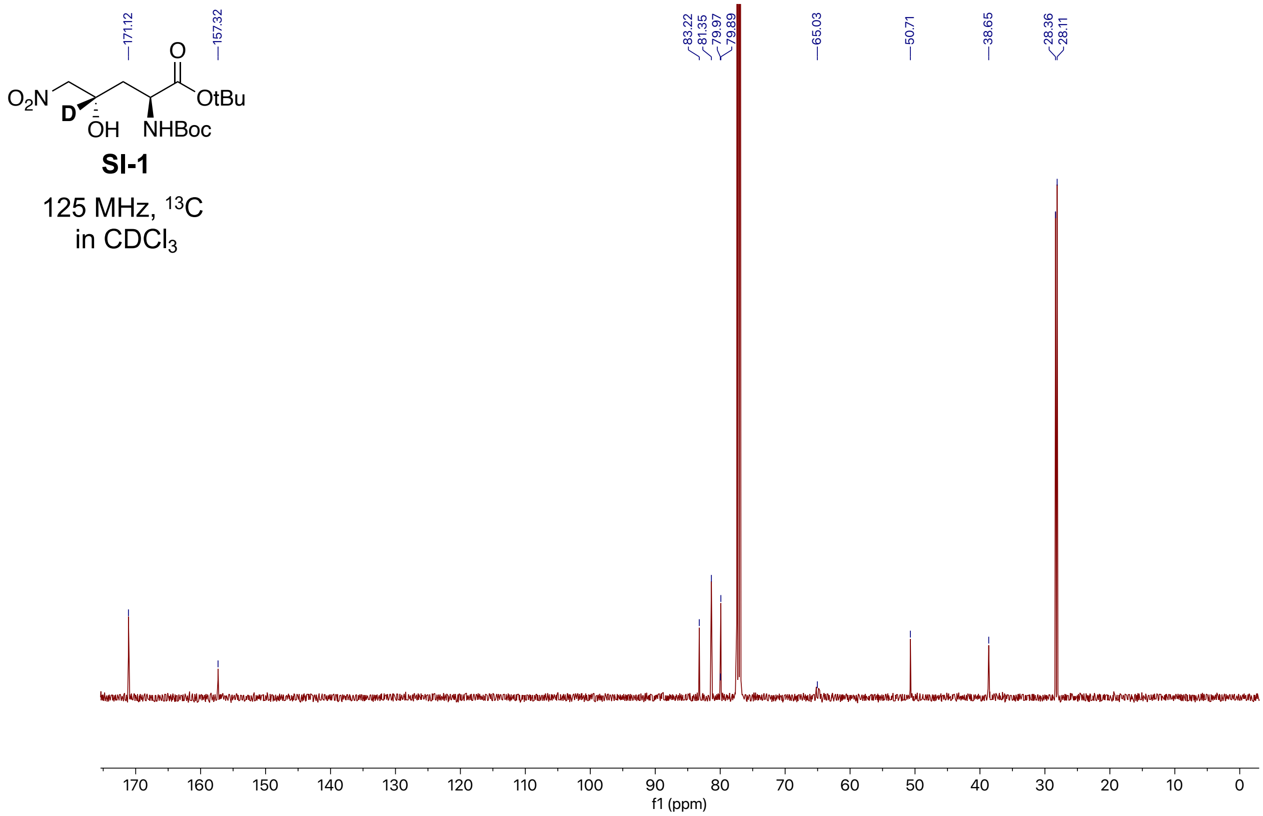

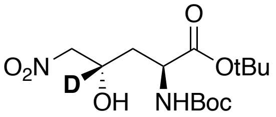

**SI-1**

500 MHz, COSY
in $CDCl_3$

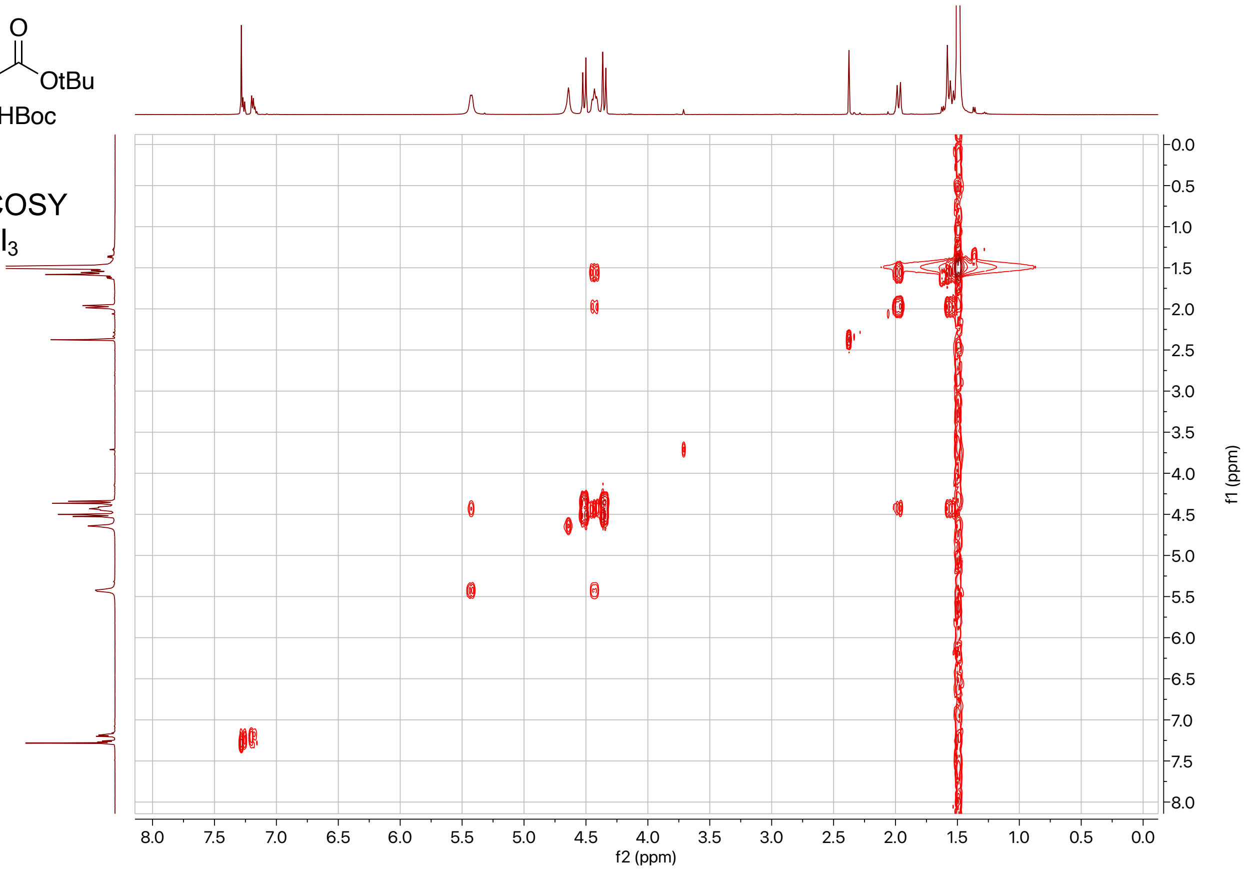

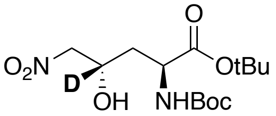

**SI-1**

500 MHz, HSQC
in $CDCl_3$

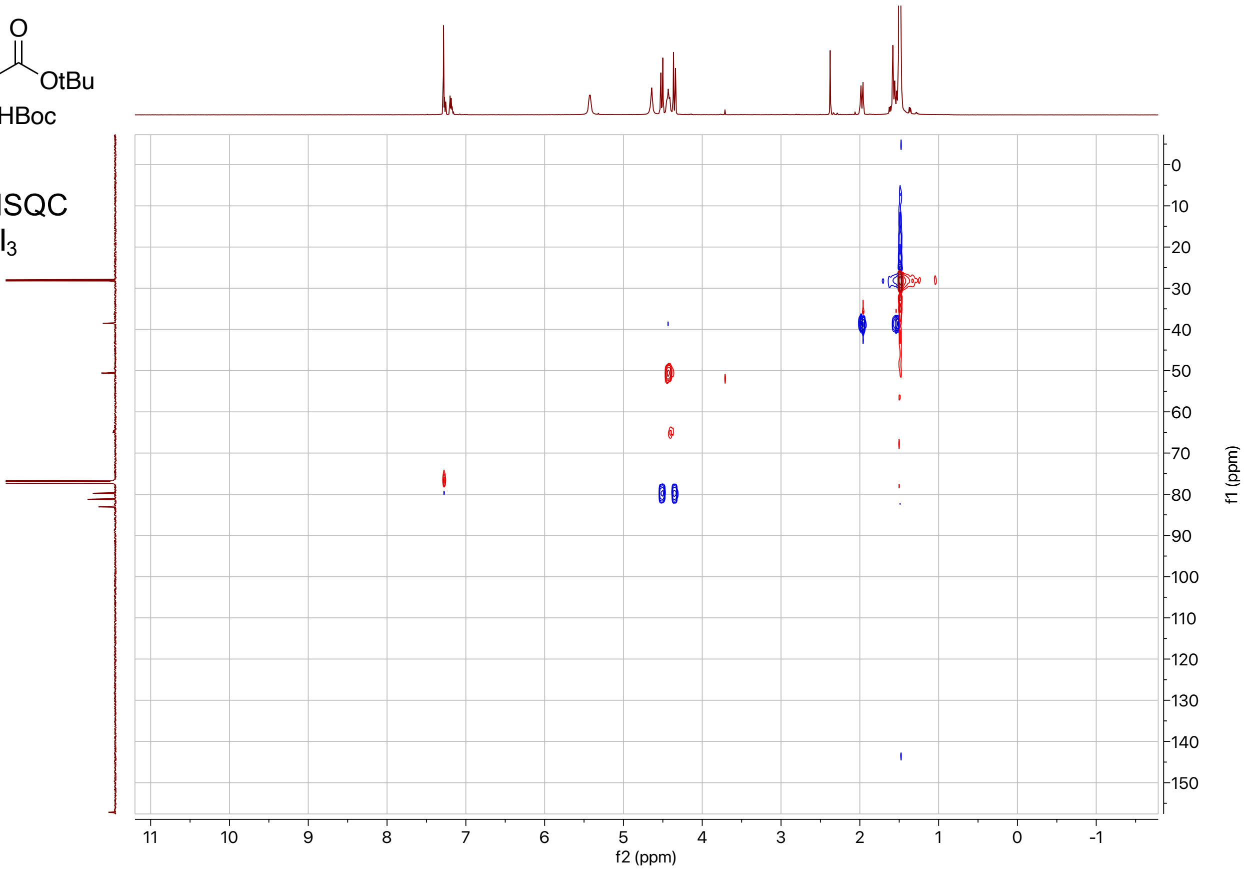

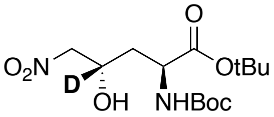

**SI-1**

500 MHz, HMBC
in $CDCl_3$

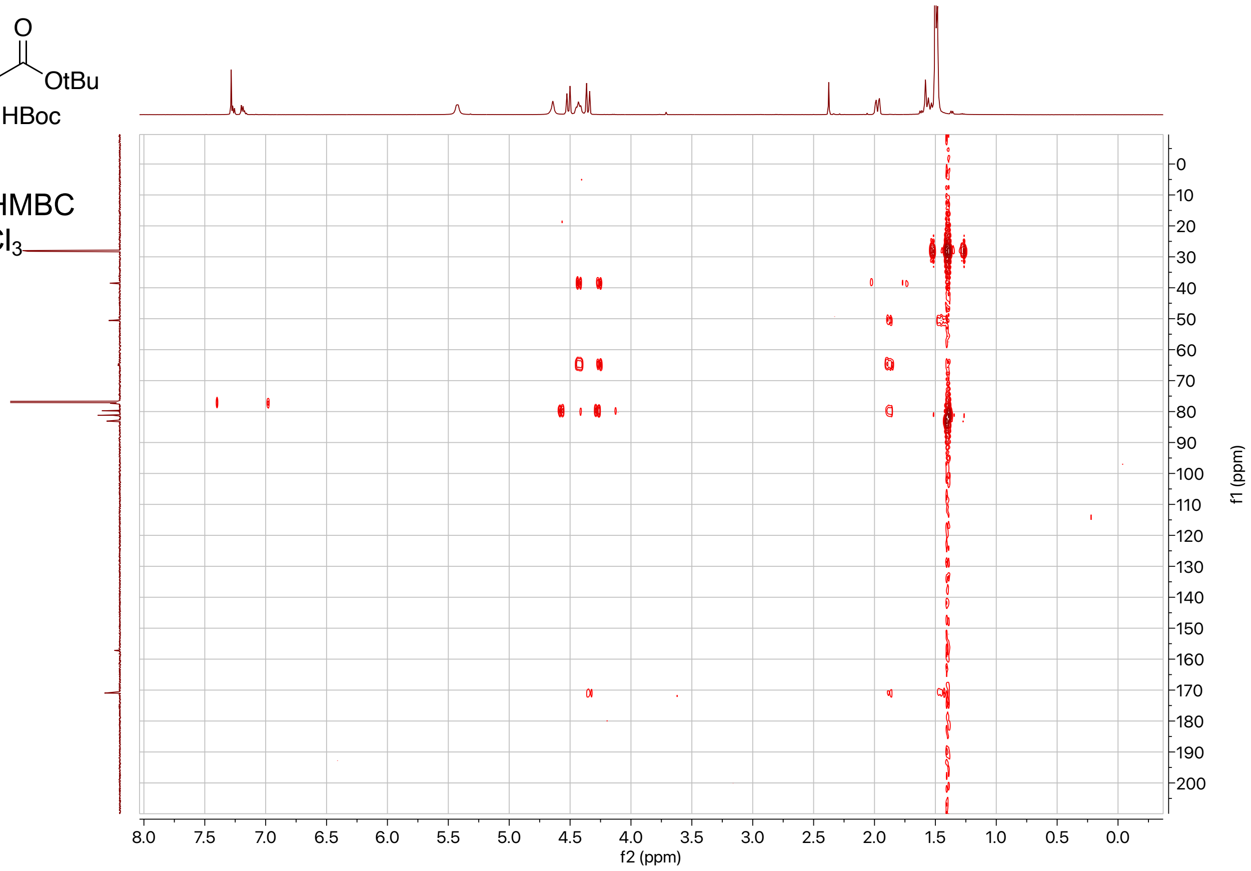

**SI-2:**

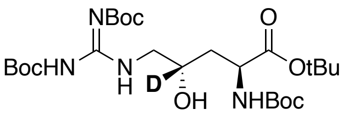

**SI-2**

500 MHz, $^{1}H$
in $CDCl_3$

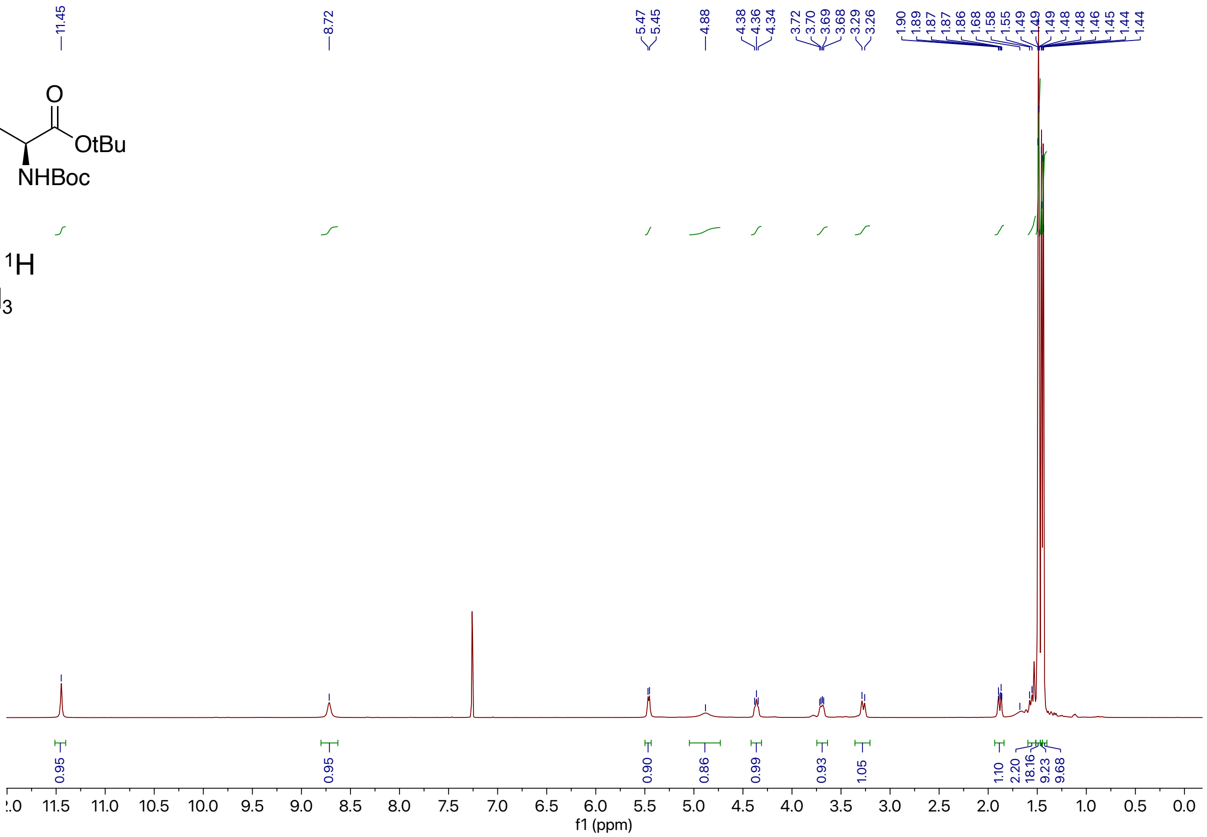

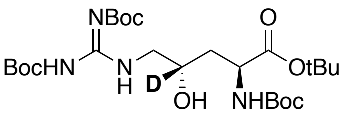

**SI-2**

500 MHz, $^{1}H$
in $CDCl_3$

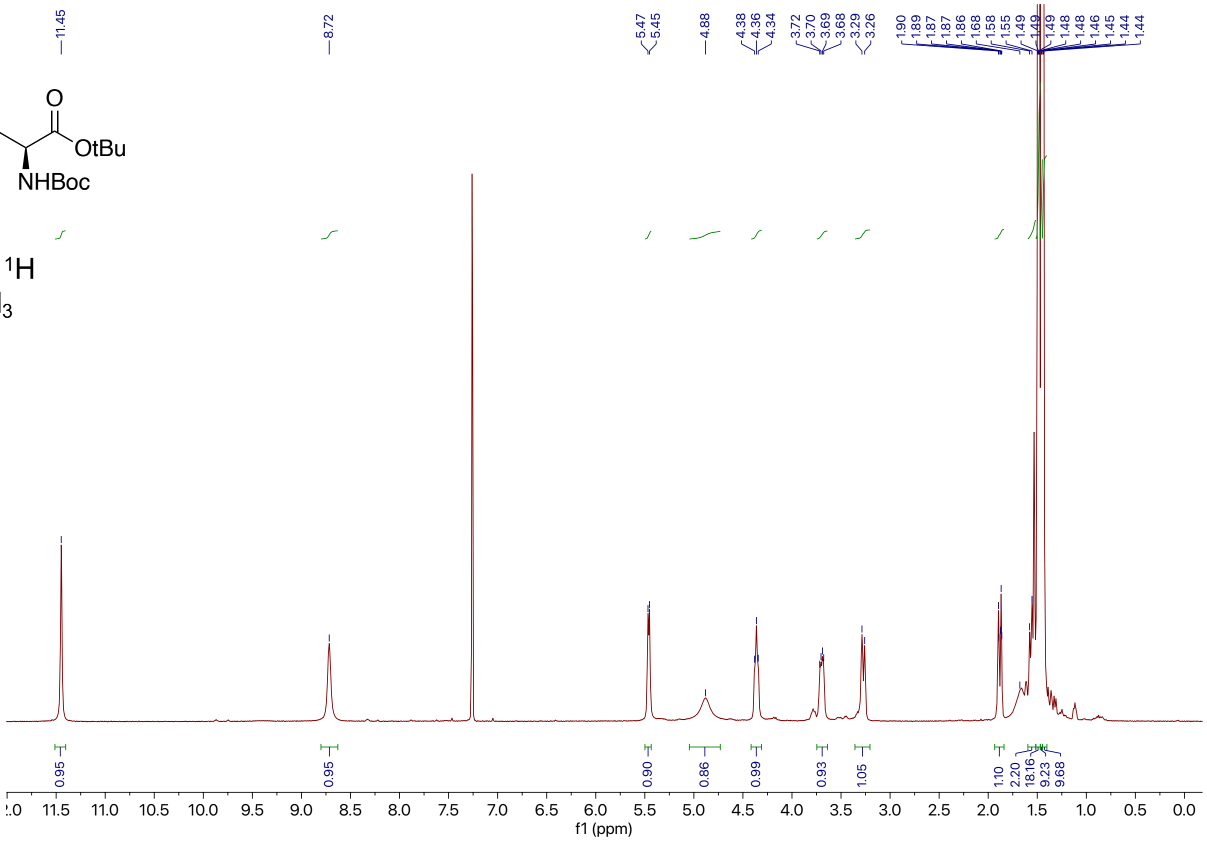

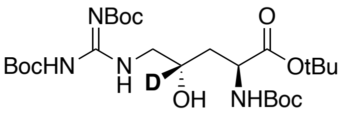

**SI-2**

125 MHz, $^{13}$C
in $CDCl_3$

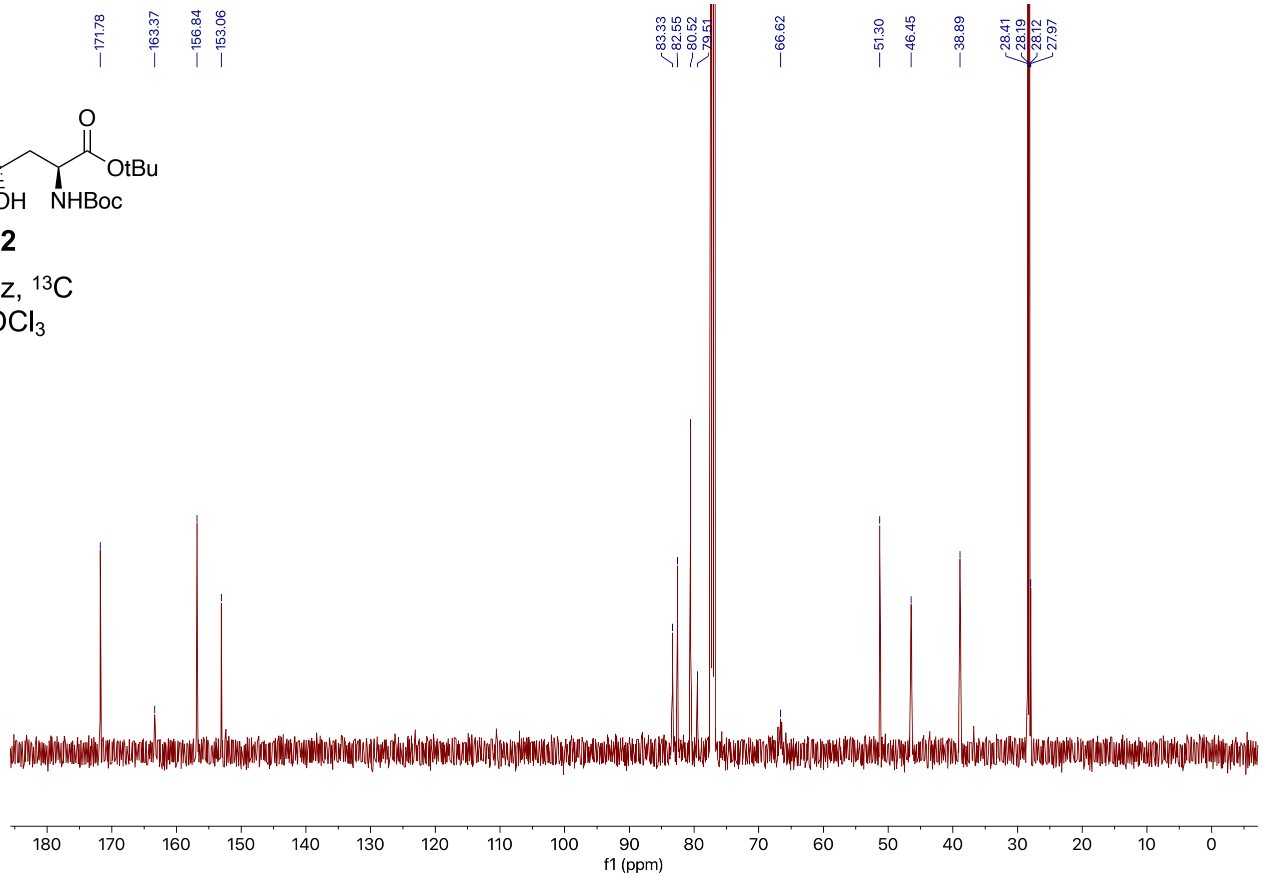

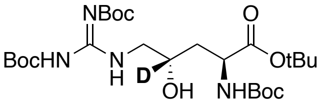

**SI-2**

125 MHz, $^{13}$C
in $CDCl_3$

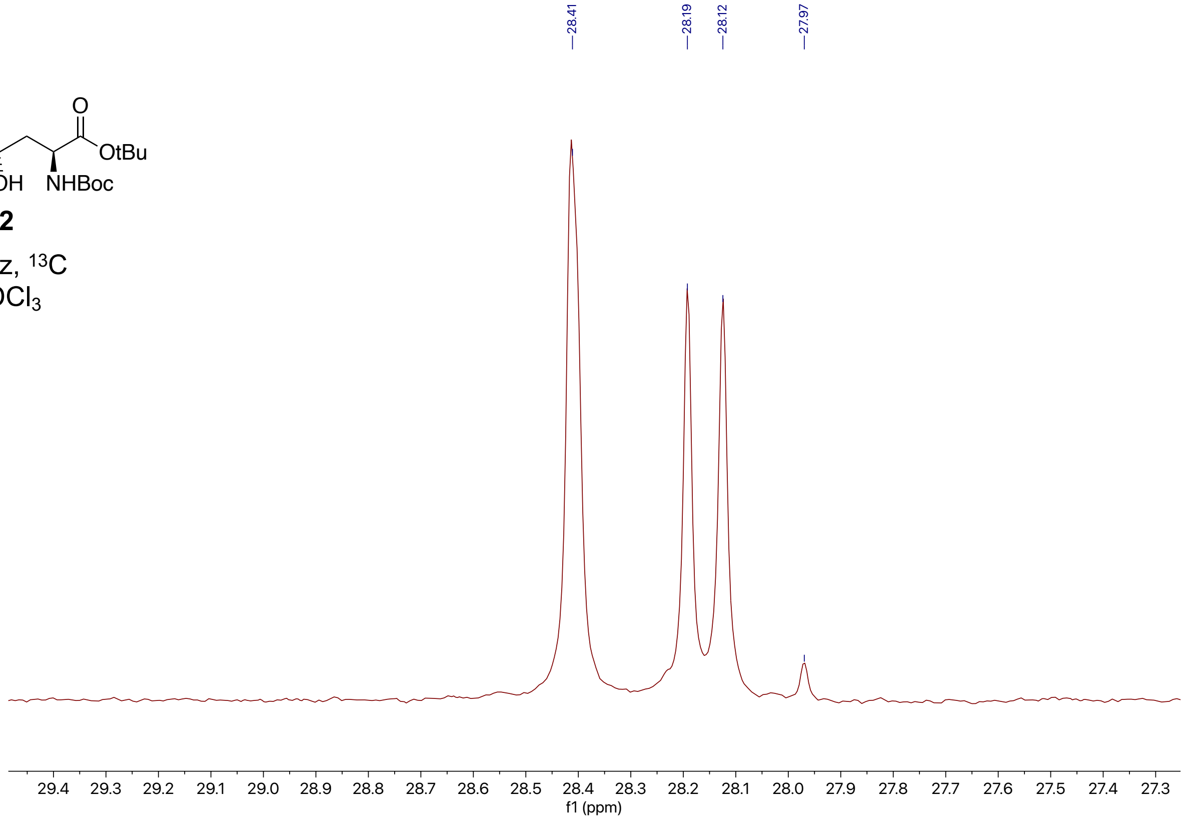

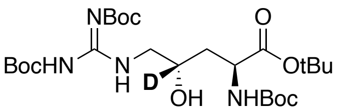

**SI-2**

500 MHz, COSY
in $CDCl_3$

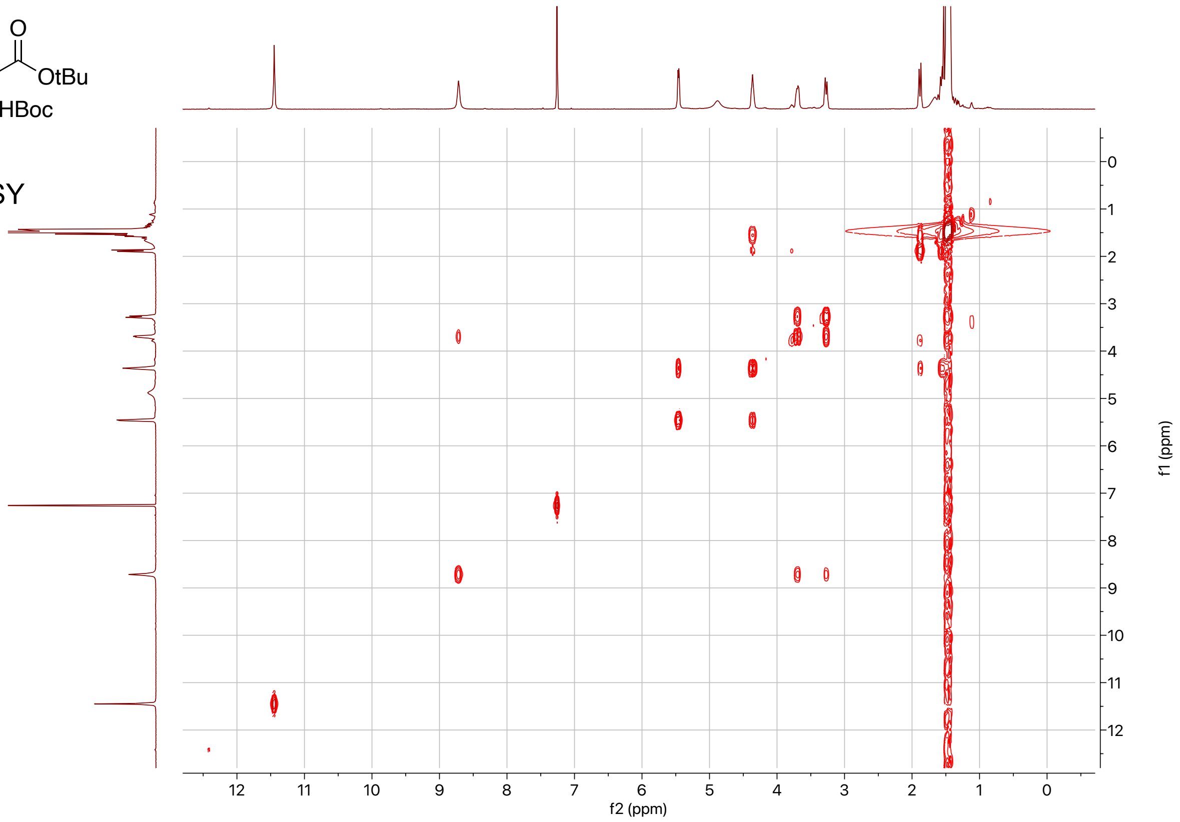

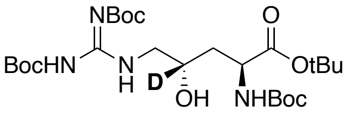

**SI-2**

500 MHz, HSQC
in $CDCl_3$

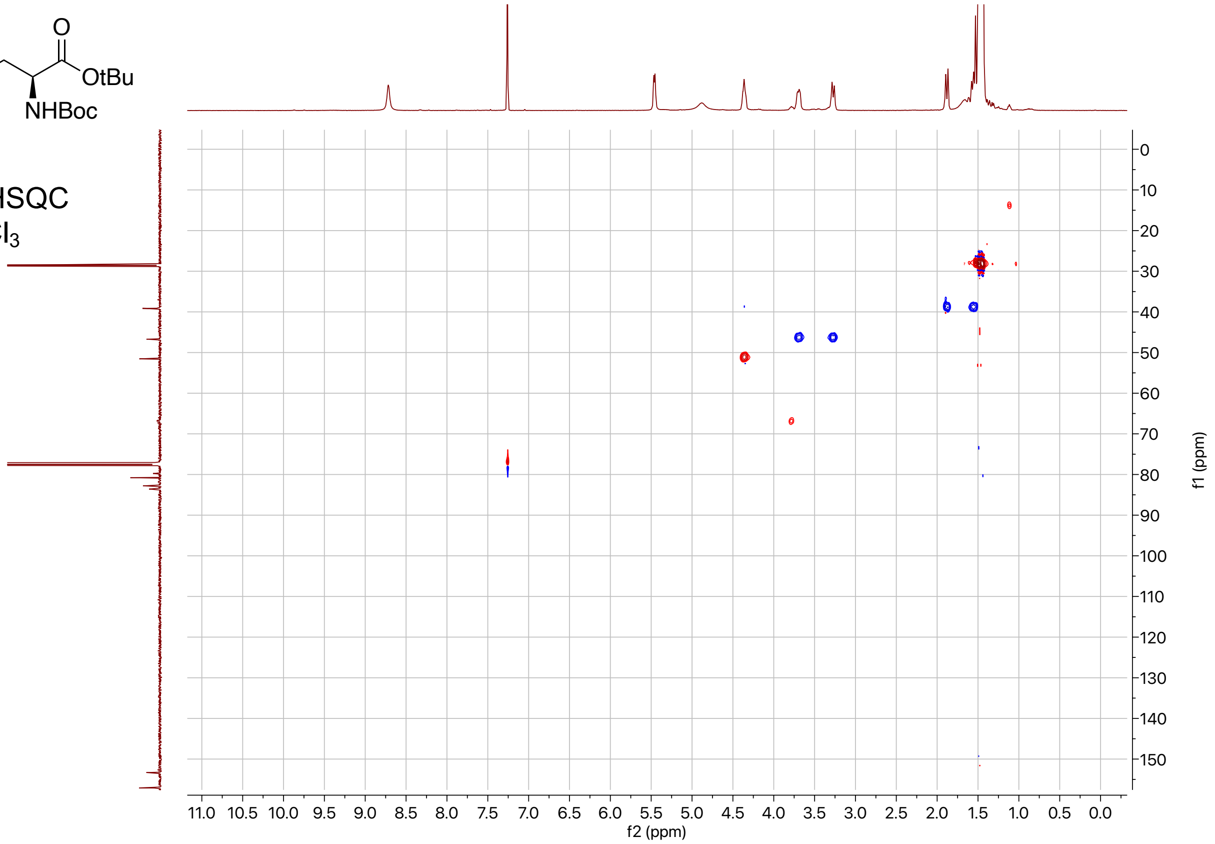

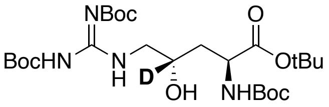

**SI-2**

500 MHz, HMBC
in $CDCl_3$

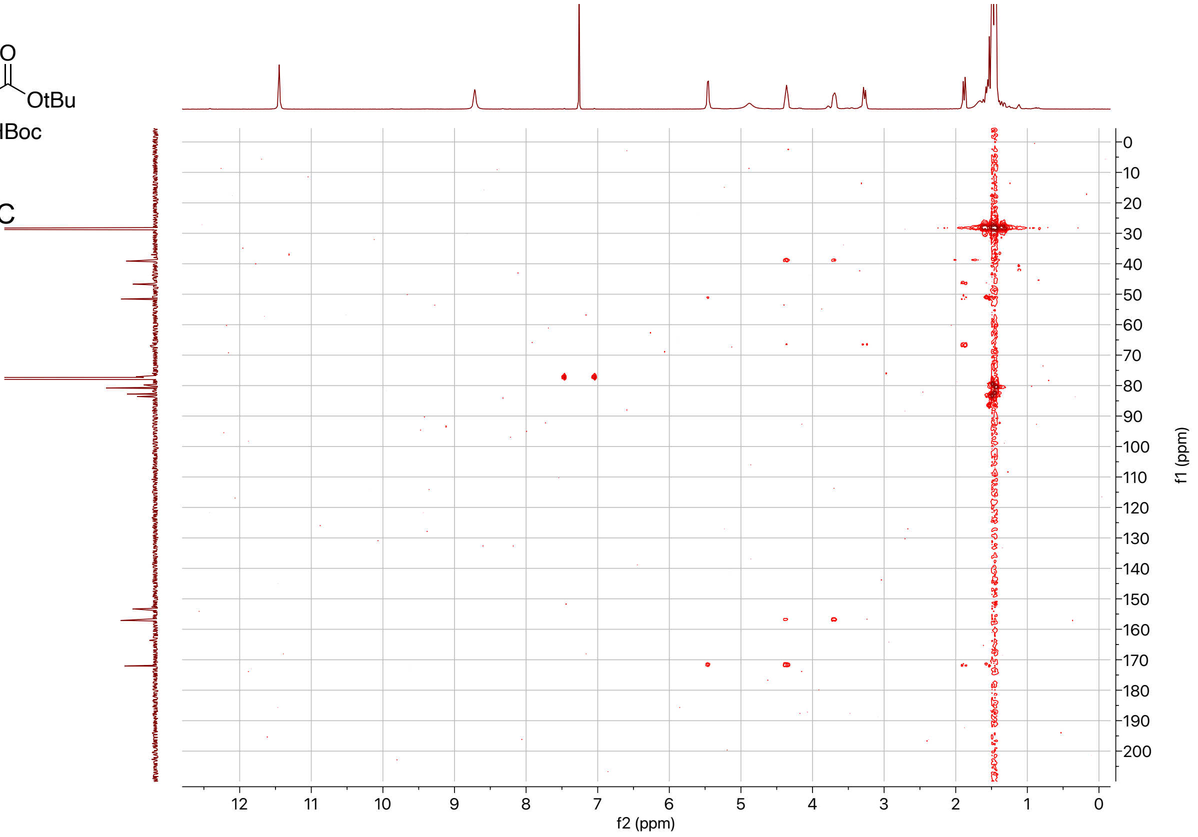

**1-$d_\gamma$:**

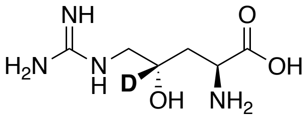

**1-$d_\gamma$**

500 MHZ, $^{1}H$
in $D_2O$ + 0.1% MeOH

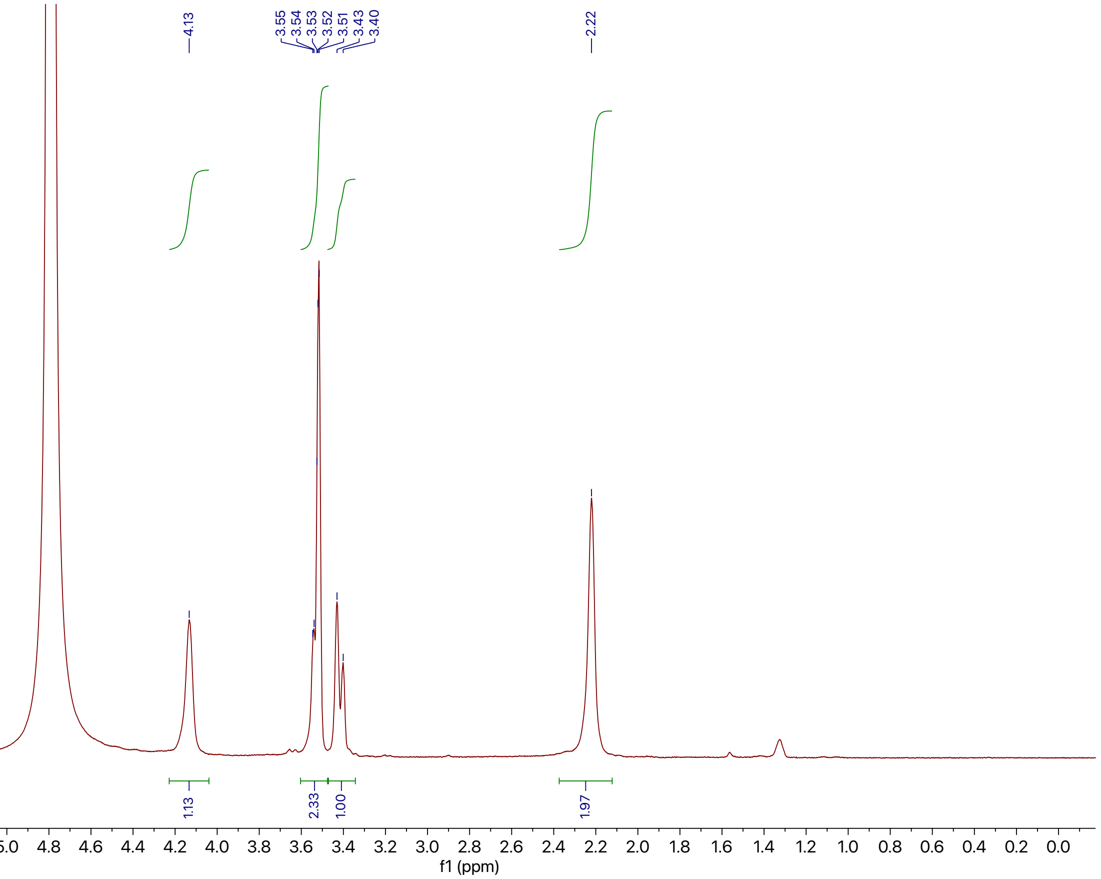

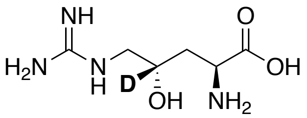

**1-$d_\gamma$**

125 MHZ, $^{13}C$
in $D_2O$ + 0.1% MeOH

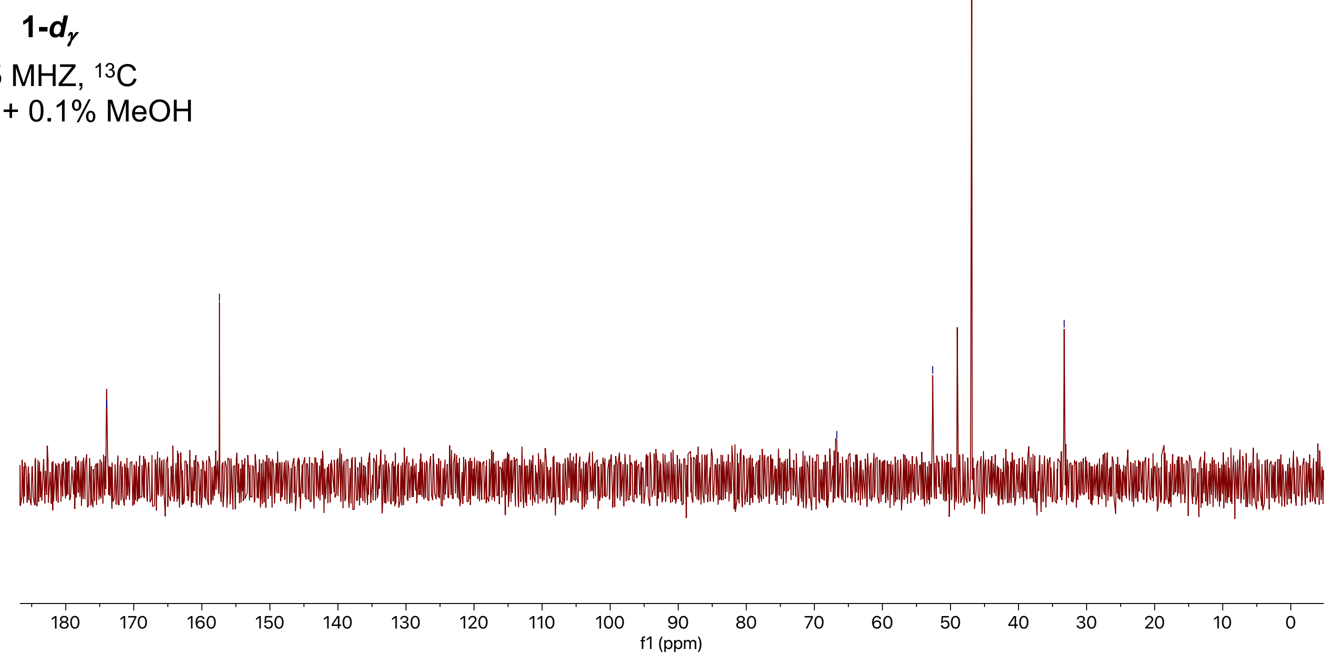

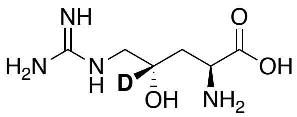

**1-*d*$_\gamma$**

500 MHz, COSY
in $D_2O$ + 0.1% MeOH

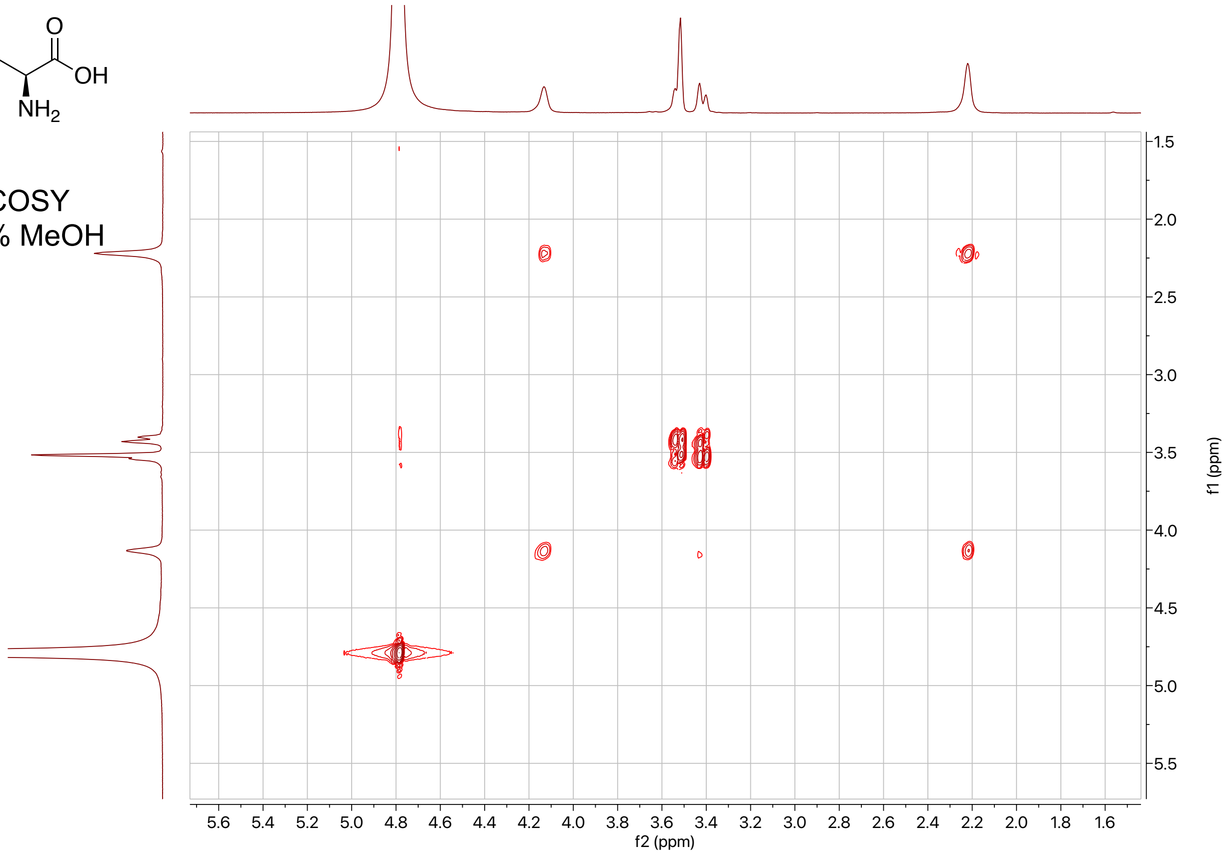

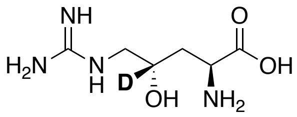

**1-*d*$_\gamma$**

500 MHz, HSQC
in $D_2O$ + 0.1% MeOH

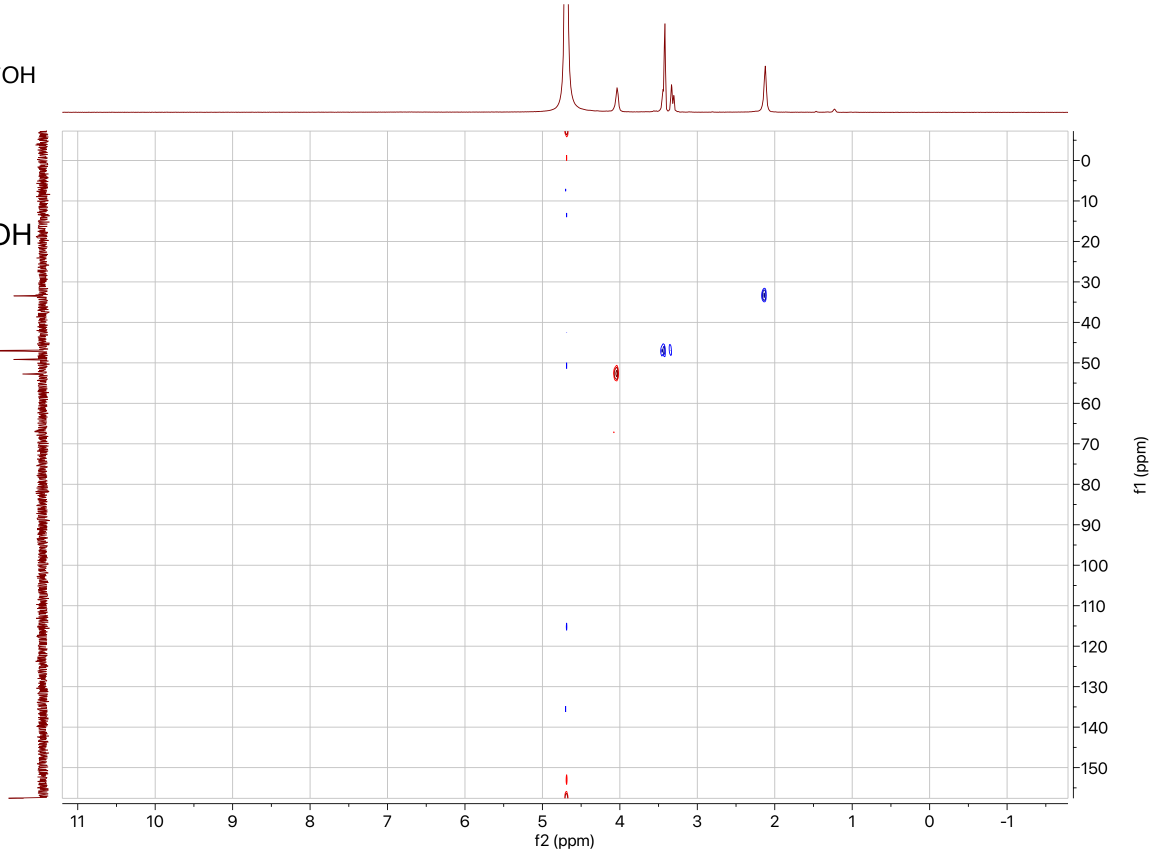

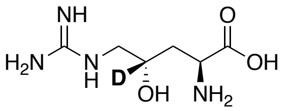

**1-*d*$_\gamma$**

500 MHZ, HMBC
in $D_2O$ + 0.1% MeOH

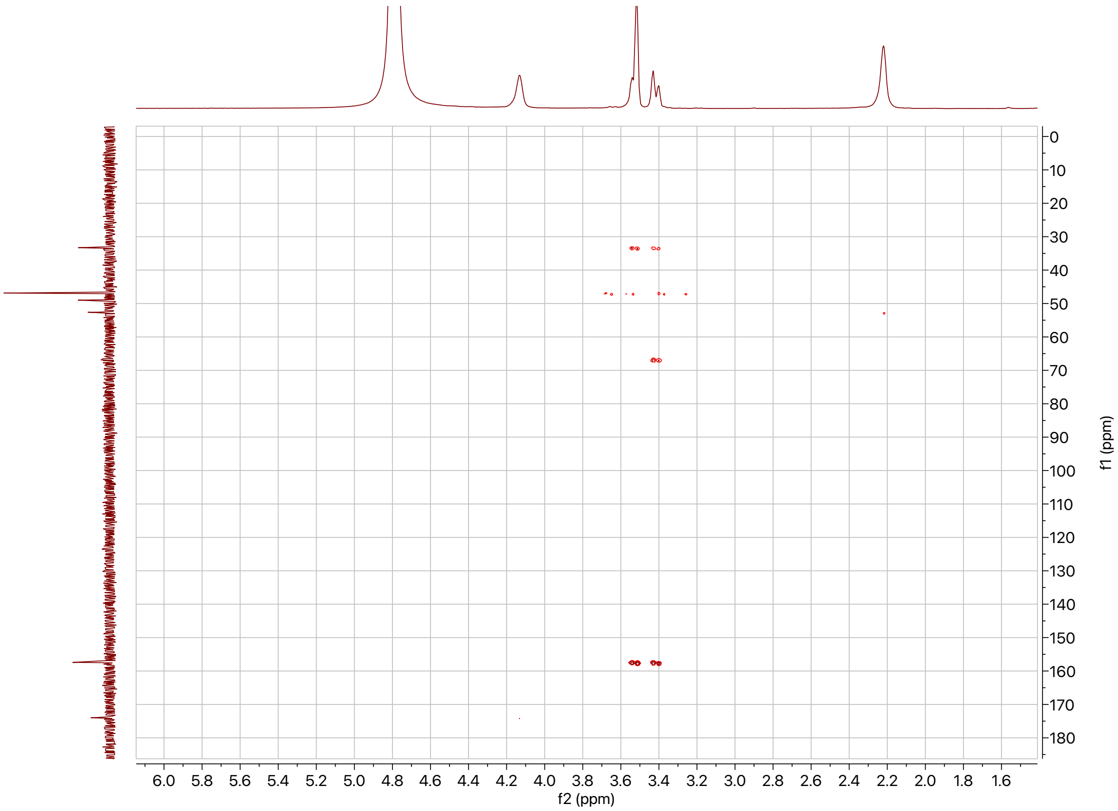

### References

(1) Lima, S. T.; Fallon, T. R.; Cordoza, J. L.; Chekan, J. R.; Delbaje, E.; Hopiavuori, A. R.; Alvarenga, D. O.; Wood, S. M.; Luhavaya, H.; Baumgartner, J. T.; Dörr, F. A.; Etchegaray, A.; Pinto, E.; McKinnie, S. M. K.; Fiore, M. F.; Moore, B. S. Biosynthesis of guanitoxin enables global environmental detection in freshwater cyanobacteria. *J. Am. Chem. Soc.* **2022**, *144* (21), 9372–9379.

(2) Chun, S. W.; Narayan, A. R. H. Biocatalytic, stereoselective deuteration of $\alpha$-amino acids and methyl esters. *ACS Catal.* **2020**, *10* (13), 7413–7418.

(3) Kumar, P.; Meza, A.; Ellis, J. M.; Carlson, G. A.; Bingman, C. A.; Buller, A. R. L-Threonine transaldolase activity is enabled by a persistent catalytic intermediate. *ACS Chem. Biol.* **2021**, *16* (1), 86–95.

(4) Vonrhein, C.; Flensburg, C.; Keller, P.; Sharff, A.; Smart, O.; Paciorek, W.; Womack, T.; Bricogne, G. Data processing and analysis with the *AutoPROC* toolbox. *Acta Crystallogr. D Biol. Crystallogr.* **2011**, *67* (4), 293–302.

(5) Kabsch, W. *XDS*. *Acta Crystallogr. D Biol. Crystallogr.* **2010**, *66* (2), 125–132.

(6) Liebschner, D.; Afonine, P. V.; Baker, M. L.; Bunkóczi, G.; Chen, V. B.; Croll, T. I.; Hintze, B.; Hung, L.-W.; Jain, S.; McCoy, A. J.; Moriarty, N. W.; Oeffner, R. D.; Poon, B. K.; Prisant, M. G.; Read, R. J.; Richardson, J. S.; Richardson, D. C.; Sammito, M. D.; Sobolev, O. V.; Stockwell, D. H.; Terwilliger, T. C.; Urzhumtsev, A. G.; Videau, L. L.; Williams, C. J.; Adams, P. D. Macromolecular structure determination using x-rays, neutrons and electrons: Recent developments in *Phenix*. *Acta Crystallogr. Sect. Struct. Biol.* **2019**, *75* (10), 861–877.

(7) Bunkóczi, G.; Echols, N.; McCoy, A. J.; Oeffner, R. D.; Adams, P. D.; Read, R. J. *Phaser.MRage*: Automated molecular replacement. *Acta Crystallogr. D Biol. Crystallogr.* **2013**, *69* (11), 2276–2286.

(8) Bunkóczi, G.; Read, R. J. Improvement of molecular-replacement models with *Sculptor*. *Acta Crystallogr. D Biol. Crystallogr.* **2011**, *67* (4), 303–312.

(9) Terwilliger, T. C.; Grosse-Kunstleve, R. W.; Afonine, P. V.; Moriarty, N. W.; Zwart, P. H.; Hung, L.-W.; Read, R. J.; Adams, P. D. Iterative model building, structure refinement and density modification with the *PHENIX AutoBuild* Wizard. *Acta Crystallogr. D Biol. Crystallogr.* **2008**, *64* (1), 61–69.

(10) Emsley, P.; Lohkamp, B.; Scott, W. G.; Cowtan, K. Features and development of *Coot*. *Acta Crystallogr. D Biol. Crystallogr.* **2010**, *66* (4), 486–501.

(11) Hemscheidt, T.; Burgoyne, D. L.; Moore, R. E. Biosynthesis of anatoxin-a(*s*). (2*S*,4*S*)-4-Hydroxyarginine as an intermediate. *J. Chem. Soc. Chem. Commun.* **1995**, *2*, 205–206.

(12) Chen, M.; Liu, C.-T.; Tang, Y. Discovery and biocatalytic application of a PLP-dependent amino acid $\gamma$-substitution enzyme that catalyzes C–C bond formation. *J. Am. Chem. Soc.* **2020**, *142* (23), 10506–10515.

(13) Robert, X.; Gouet, P. Deciphering key features in protein structures with the new ENDscript server. *Nucleic Acids Res.* **2014**, *42* (W1), W320–W324.

(14) Jumper, J.; Evans, R.; Pritzel, A.; Green, T.; Figurnov, M.; Ronneberger, O.; Tunyasuvunakool, K.; Bates, R.; Žídek, A.; Potapenko, A.; Bridgland, A.; Meyer, C.; Kohl, S. A. A.; Ballard, A. J.; Cowie, A.; Romera-Paredes, B.; Nikolov, S.; Jain, R.; Adler, J.; Back, T.; Petersen, S.; Reiman, D.; Clancy, E.; Zielinski, M.; Steinegger, M.; Pacholska, M.; Berghammer, T.; Bodenstein, S.; Silver, D.; Vinyals, O.; Senior, A. W.; Kavukcuoglu, K.; Kohli, P.; Hassabis, D. Highly accurate protein structure prediction with AlphaFold. *Nature* **2021**, *596* (7873), 583–589.

(15) Chang, C.-Y.; Lyu, S.-Y.; Liu, Y.-C.; Hsu, N.-S.; Wu, C.-C.; Tang, C.-F.; Lin, K.-H.; Ho, J.-Y.; Wu, C.-J.; Tsai, M.-D.; Li, T.-L. Biosynthesis of streptolidine involved two unexpected intermediates produced by a dihydroxylase and a cyclase through unusual mechanisms. *Angew. Chem. Int. Ed.* **2014**, *53* (7), 1943–1948.

(16) Clausen, T.; Huber, R.; Prade, L.; Wahl, M. C.; Messerschmidt, A. Crystal structure of *Escherichia coli* cystathionine $\gamma$-synthase at 1.5 Å resolution. *EMBO J.* **1998**, *17* (23), 6827–6838.

(17) Hai, Y.; Chen, M.; Huang, A.; Tang, Y. Biosynthesis of mycotoxin fusaric acid and application of a PLP-dependent enzyme for chemoenzymatic synthesis of substituted L-Pipecolic Acids. *J. Am. Chem. Soc.* **2020**, *142* (46), 19668–19677.

(18) Yee, D. A.; Niwa, K.; Perlatti, B.; Chen, M.; Li, Y.; Tang, Y. Genome mining for unknown–unknown natural products. *Nat. Chem. Biol.* **2023**.

(19) Cui, Z.; Overbay, J.; Wang, X.; Liu, X.; Zhang, Y.; Bhardwaj, M.; Lemke, A.; Wiegmann, D.; Niro, G.; Thorson, J. S.; Ducho, C.; Van Lanen, S. G. Pyridoxal-5′-phosphate-dependent alkyl transfer in nucleoside antibiotic biosynthesis. *Nat. Chem. Biol.* **2020**, *16* (8), 904–911.

(20) Hauth, F.; Buck, H.; Stanoppi, M.; Hartig, J. S. Canavanine utilization via homoserine and hydroxyguanidine by a PLP-dependent γ-lyase in *Pseudomonadaceae* and *Rhizobiales*. *RSC Chem. Biol.* **2022**, *3* (10), 1240–1250.

(21) Du, Y.-L.; Ryan, K. S. Pyridoxal phosphate-dependent reactions in the biosynthesis of natural products. *Nat. Prod. Rep.* **2019**, *36* (3), 430–457.

(22) Giltrap, A. M.; Dowman, L. J.; Nagalingam, G.; Ochoa, J. L.; Linington, R. G.; Britton, W. J.; Payne, R. J. Total synthesis of teixobactin. *Org. Lett.* **2016**, *18* (11), 2788–2791.
